# Oxylipins and Other Natural Products Produced by UV-sterilization Impact Commercial Oyster Larvae Production in a New England Estuary

**DOI:** 10.64898/2026.07.29.738347

**Authors:** Bethanie R Edwards, Meredith White, Scarlett Schroeder, Amanda Clapp, Bill Mook, Rich Smith, Andy Stevenson, Steve Zimmerman

## Abstract

Here, oyster larval abnormalities within a hatchery were linked to a common UV sterilization technique used in aquaculture. Because of the known link between phytoplankton oxylipins and zooplankton mortality, we hypothesized that UV-treatment of seawater results in the production of oxylipins that inhibit larval digestion of microalgae. We used lipidomics to observe changes in the organic compounds dissolved in estuarine seawater. UV-treatment resulted in an increase in oxylipins associated with cyanobacteria, fungi, and macroalgae in 2020, whereas oxylipins typically produced by diatoms were more abundant in the 2021 UV-treatments. Oxylipin concentrations were higher in 2020, when the hatchery reported the most severe issues. Experiments directly testing the impact of oxylipins showed exposing larvae to exogenous compounds resulted in the same digestive syndrome as the endogenous suite of compounds produced by UV. Further investigation of the lipidomes revealed that oxylipins were only one class of compounds linked to UV-treatment, with secondary metabolites associated with higher plants, a wide range of pharmaceuticals, and anthropogenic compounds also increasing under UV. Future efforts will explore the sources of these compounds, the mechanisms by which they inhibit oysters, and the extent of this emerging environmental problem for marine ecosystems globally.

**Graphical Abstract:** a) Experiments exploring the dissolved metabolome of hatchery seawater treated by 1) filtration + UV-sterilization or 2) by filtration only, show UV pretreatment leads to larval death due to a digestive syndrome while filtering-only lead to shell deformities and longer metamorphosis times. b) Lipid-focused mass spectrometry and comparative metabolomic analysis suggest the causative agents are oxylipins either from phytoplankton, bacterioplankton, or macroalgal detritus less than 1uM; while anthropogenic compounds like terephthalic acid and pharmaceutical quinolones were likely liberated from submicron microplastics or organic matter to which they are known to absorb.

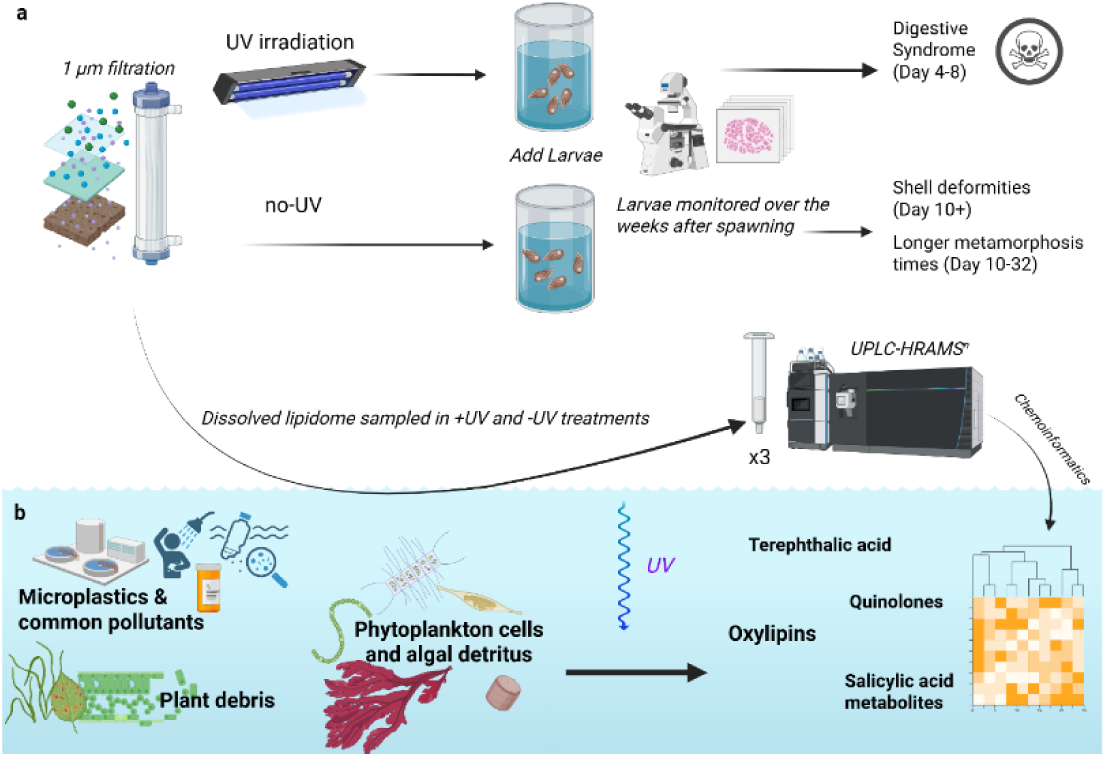

## 1. Introduction

Roughly sixty-seven percent of all oysters produced in Maine come from the Damariscotta River, making the system important for the local economy and cold-water aquaculture nationally (Maine Department of Marine Resources (MEDMR), 2020). Mook Sea Farm (MSF) is a vertically integrated oyster farm located on the Damariscotta River in Walpole, Maine, USA. The hatchery at MSF produces oyster (*Crassostrea virginica*) seed (1-12 mm juveniles) for grow-out to market size on their own leases, as well as for sale to over 100 oyster farms along the East Coast from Maine to North Carolina. We begin by outlining an aquaculture problem observed at the MSF hatchery in the Introduction and then expand on the lipidomic analysis used to determine the causative agents in the rest of the manuscript.

In the MSF hatchery, the standard process for over 20 years has been that larvae are held in tanks filled with 1 μm filtered and UV-irradiated (∼52,000 μW/s/cm^2^) estuary water (average 29 ppt). This pretreatment removes pathogens and provides controlled conditions for larval growth. While common UV pretreatment is not used by all bivalve hatcheries (Helm & Bourne, 2004).

### 1.1 Digestive syndrome in oyster larvae

Starting with the first spawning of the season in January 2020, Mook Sea Farm noticed abnormal larval development presenting as a digestive syndrome, which resulted in a decline in seed sales to 56% of the 2019 values (**table 1**; **figure 1**). The larvae developed normally to the D-stage veliger at 2 d of age, but by day 4, it was obvious through microscopic examination that they could not digest their microalgal food after ingestion (White et al., 2022). Undigested microalgal cells were visible in the stomachs of the larvae, but no digested material was visible in the digestive glands (**figure 1a**). By day 8, the larvae should have doubled their size, yet little to no growth occurred (**figure 1a**). The hatchery was forced to discard over 100,000,000 larvae from the first spawn and began a months-long investigation to understand the cause of the digestive syndrome. Histology performed on a new spawn of larvae in February 2020 showed necrosis of the gastric epithelium and debris in the gastric lumen (**figure 2a,b**), consistent with contaminants or toxins entering the larval digestive system.

**Figure 1.**
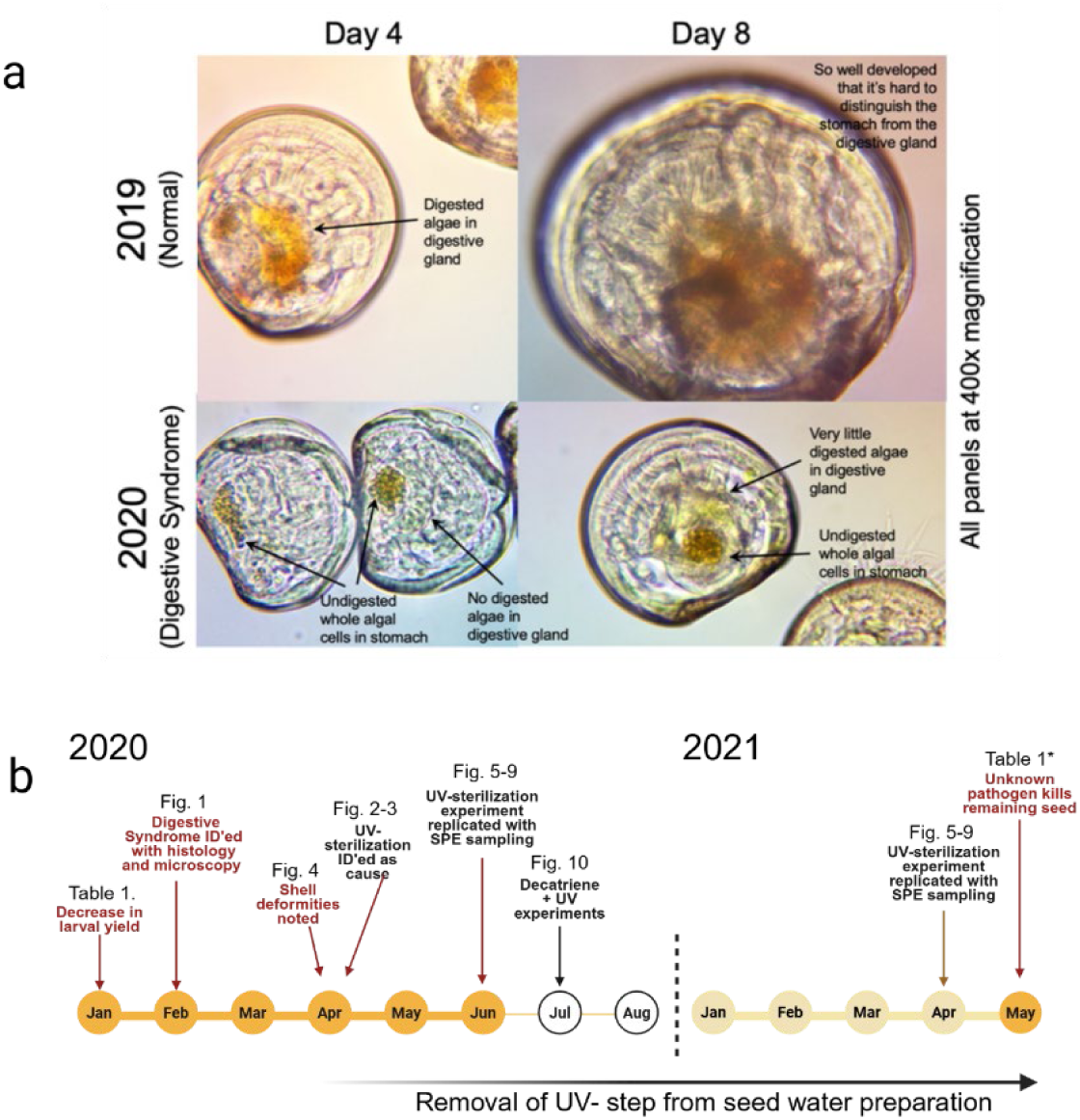
Evidence of UV induced digestive syndrome and timeline. **A)** Comparison of normal oyster larvae produced in 2019 vs. abnormal larvae produced in 2020. All images are at the same scale and were taken under 400x magnification. Normal day 4 larvae from 2019 showed dark golden color of digested microalgae food in their digestive gland, with little to no undigested algal cells in the stomach. In 2020, day 4 larvae had large numbers of undigested whole algal cells in their stomachs, with no digested algae in their digestive glands, symptoms that became known as the larval digestive syndrome. These larvae also showed minor shell deformities. These impaired larvae in 2020 showed little growth from day 4 to day 8 and were obviously smaller than a normal day 8 larva from 2019. The 2020 day 8 larvae showed a small amount of digested material in their digestive gland, while the normal 2019 day 8 larvae have developed to the point where it is hard to distinguish the stomach from the digestive gland. **B)** Timeline of events and experimentation over the 2020 and 2021 spawning season with the corresponding figures and tables with data.

**Figure 2.**
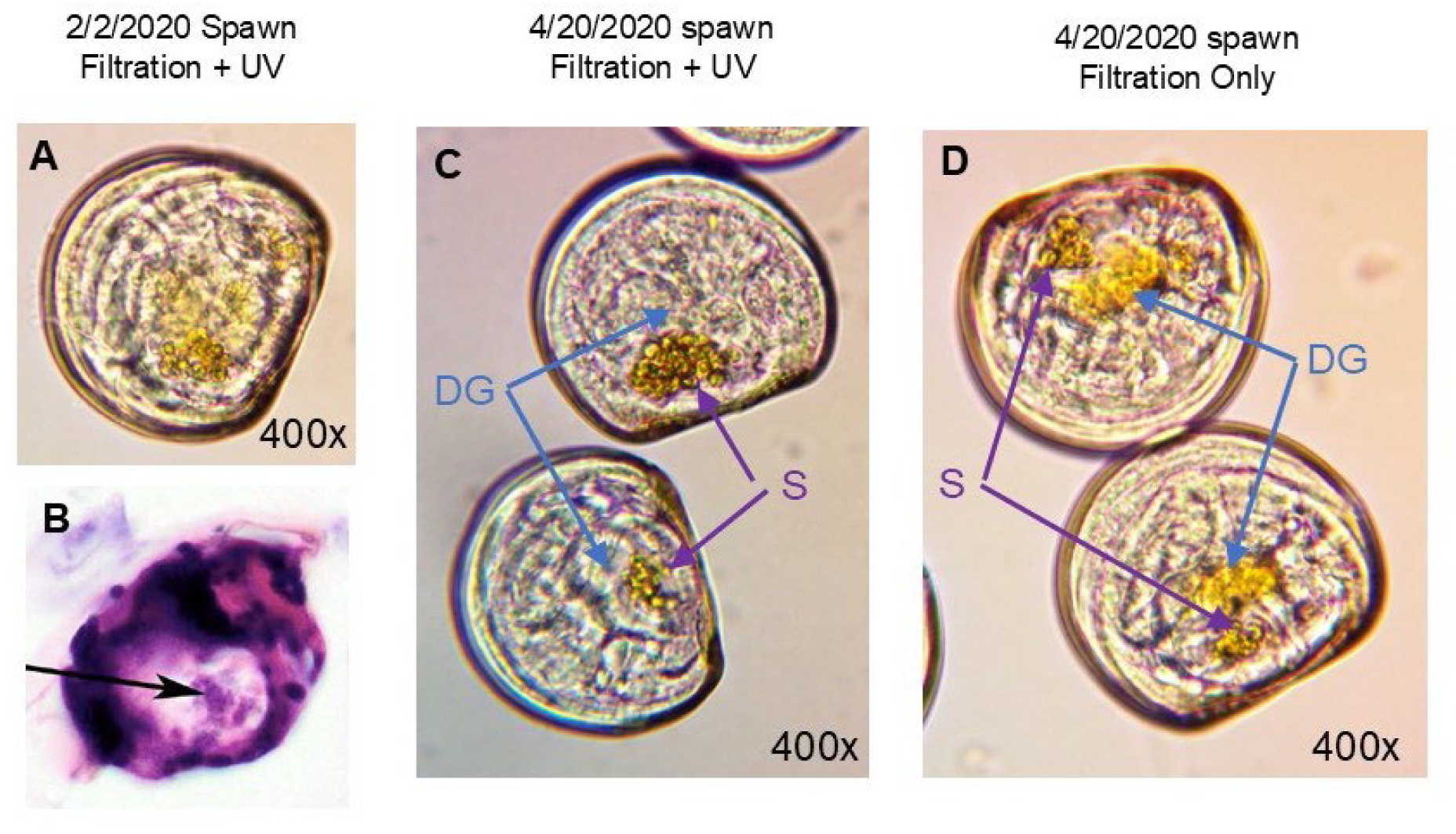
Two day old oyster larvae from spawns on a,b) February 2, 2020 and c,d) April 20, 2020 at the Mook Sea Farm Hatchery, exposed to 1 μm filtration and ∼52,000 μW/s/cm^2^ UV irradiation (**A,B,C**) or exposed to 1 μm filtration only (**D**). Micrographs in **A**, **C**, and **D** are at the same scale and were taken under 400x magnification. Larvae from the February 2 spawn were examined through histology (**B**), which showed necrosis of the gastric epithelium and debris in the gastric lumen (black arrow). The April 20 spawn was an experimental spawn to help identify the cause of the digestive syndrome seen since early January, 2020. Larvae exposed to UV irradiation (**C**) showed the same symptoms of the digestive syndrome that had been noted since January 2020, undigested microalgal cells in the stomach (S), with little to no digested algal material in the digestive gland (DG). Larvae exposed to filtration only (**D**) had significantly fewer undigested algal cells in the stomach and showed the golden color of digested algal material in the digestive gland, which is what is typically seen in healthy 2-day old oyster larvae.

**Table 1.** Shortfalls in Mook Sea Farm hatchery production in 2020 and 2021, relative to 2019, one of the best production years on record. Exact revenue in US dollars and number of oyster seed sold (millions) are proprietary. Revenue is not linearly related to the number of seed sold because the price per 1000 seed varies based on the size of the seed.

|  | 2019 | 2020 | 2021 |
|---|---|---|---|
| Revenue from oyster seed sales (\$) | 100% | 56.0% | 21.1% |
| Number of oyster seed sold | 100% | 84.9% | 34.0% |

On April 20, 2020, we determined that the UV treatment was related to the digestive syndrome (**figure 1b**). Larval digestion was compared between the standard protocol (1 µm filtration and UV irradiation) and a modified technique of 1 µm filtration only. Larvae in the filtration-only (no-UV) treatment were able to digest their microalgal food, while the larvae exposed to water that was filtered and UV-sterilized were unable to digest their food (**figure 2c,d**). The filtration-only treatment was adopted by the hatchery for the remainder of the season, but because it was adopted so late in the production season, revenue from seed sales in 2020 was still severally impacted (**table 1**).

### 1.2 Shell deformities and lagging metamorphosis

Later in the 2020 season, when the larvae were able to digest their food, grow, and metamorphose into juveniles (seed), the hatchery staff noticed two additional changes from previous years: shell deformities and a longer time to metamorphosis. Older (>10 d) larvae developed novel and severe shell deformities (**figure 3**), named duckbill and cinnamon roll deformities by hatchery staff. Shell deformities are somewhat common in larvae that are 1-4 days old but have not been noted in older larvae at MSF.

**Figure 3.**
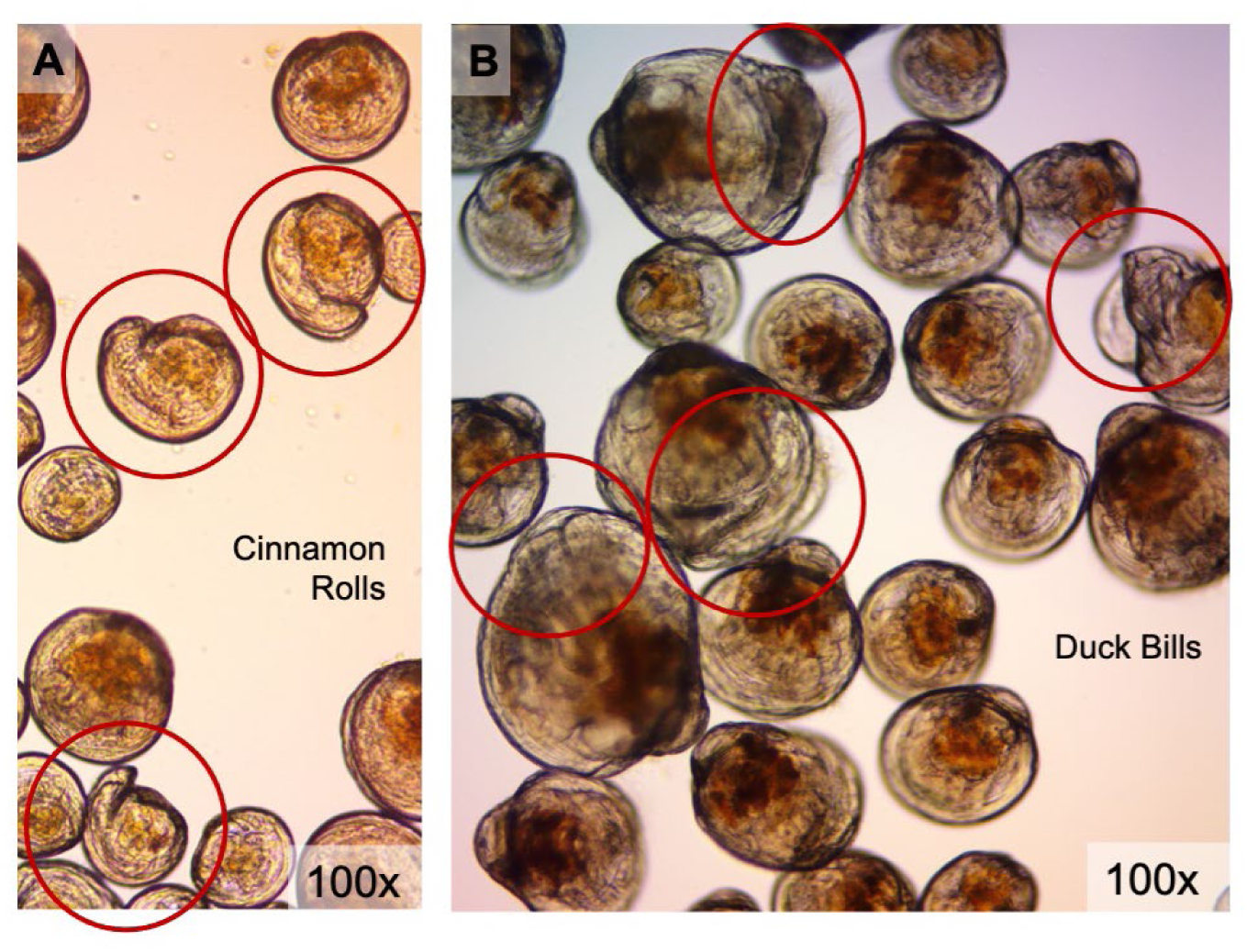
After the hatchery adopted the filtration-only seawater treatment in late April, 2020, larvae were able to digest their microalgal food and develop into later-stage larvae, but staff noticed novel shell deformities that developed starting on day 10. These 10-day old larvae show what hatchery staff dubbed ‘cinnamon roll’ (**A**) and ‘duck bill’ (**B**) shell deformities, highlighted by red circles. Images are at the same scale and were taken under 100x magnification. The disparate sizes of larvae in these images is also outside of the standard production practice of of culling the smallest larvae at each water change to maintain a population of similar-sized, ‘best performing’ larvae. Because so much of the early production season was lost, the hatchery had to keep all sizes of larvae to maintain numbers needed for sales.

Additionally, larvae produced in previous years were competent to ‘set’ (metamorphose to juveniles) by day 14; the cohorts produced using the filtration-only treatment took as long as 24 days to reach setting competency (**Table 2**). As the hatchery season progressed, filtration-only water was used, and the symptoms of the larvae improved. Larvae from a spawn on June 19, 2020, took 18 days to reach setting competency and showed severe duckbill shell deformities (**figure 3**) after day 10. Larvae from a spawn on June 30, 2020, took 16 days to reach setting competency and showed few very minor shell deformities. Larvae from a spawn on July 14, 2020, took the normal 14 days to reach setting competency and did not have any shell deformities. The decline in larval symptoms indicates that the exogenous factor from the ambient estuary water also declined from mid-June to mid-July 2020.

**Table 2.** Age of setting competency for larval cohorts spawned March 6, 2020 – July 14, 2020. A well performing larval cohort should be fully competent to set by day 14, which was only accomplished for the final cohort of the season. 10 spawns were completed prior to March 6, none of which reached setting competency. Setting age improved consistently following the June 1, 2020 spawn. **The cohort spawned on April 20 was switched to filtered-only (no UV) water on April 22. *All cohorts spawned after May 1 were raised in filtration-only water.

| Spawn Date | Age of setting competency<br>(days post fertilization) |
|---|---|
| 03/06/20 | 16-24 |
| 03/17/20 | 16-26 |
| 03/30/20 | 19-30 |
| 04/09/20 | 19-22 |
| 04/20/20** | 14-20 |
| 05/01/20* | 14-18 |
| 05/21/20* | 16-20 |
| 06/01/20* | 15-24 |
| 06/19/20* | 13-18 |
| 06/30/20* | 12-16 |
| 07/14/20* | 12-14 |

In January 2021, the hatchery continued seed production using filtration-only water while conducting small-scale experiments on the temporal variability of the digestive syndrome and UV-induced chemical changes. Although UV-related impacts were milder than in 2020, filtration-only methods were maintained as a precaution, successfully yielding tens of millions of seed through early May. In mid-May 2021, ambient estuary temperatures rapidly rose from 10.5 °C to 16 °C over three weeks (Figure 4), causing complete mortality in the previously healthy seed. Pathology by Kennebec River Biosciences attributed the die-off to Pseudomonas sp. and Vibrio sp., which proliferated unchecked without UV sterilization (<1 μm). Consequently, 2021 seed sales revenue fell to 21.1% of 2019 levels (Table 1).

**Figure 4.**
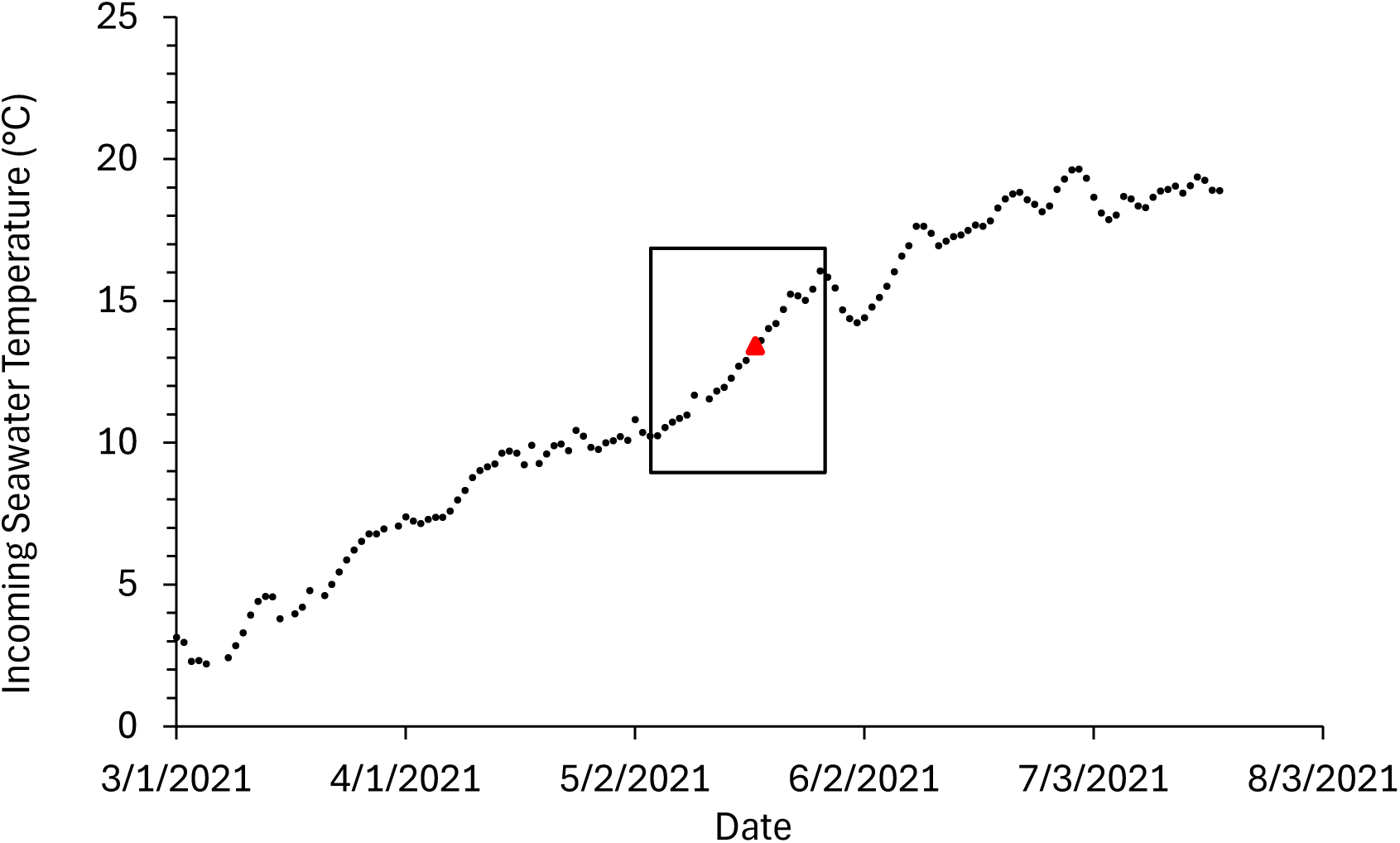
Incoming seawater temperature at the Mook Sea Farm Hatchery from March to July, 2021. The box indicates a rapid increase in incoming water temperature from 10.5 °C to 16 °C over three weeks. The red triangle indicates the first date at which mortality of seed was observed. Pathology completed by Kennebec River Biosciences indicated moderate to heavy *Pseudomonas* sp. and *Vibrio* sp. growth on the affected seed.

Following initial larval failures in January 2020, staff conducted over 85 experiments, ruling out broodstock, microalgae, packing plant runoff, filter bags, UV bulbs, tidal dynamics, temperature, pH, pCO2, and dissolved oxygen. Because symptoms improved after water changes, a chemical contaminant was suspected. Eliminating UV sterilization resolved the issue, leading researchers to hypothesize that UV photooxidation was generating oxylipins—chemical signals that are ubiquitous across the Domains of Life (bacteria and eukaryotes; archaea have not yet been investigated). Humans produce oxylipins, although they are typically referred to as lipid mediators and prostaglandins, as do land plants, where oxylipins, like jasmonic acid, play a critical role in immune response and grazing defense (Serhan et al. 2008, Spite et al. 2014, Yang et al. 2019). In marine systems, oxylipins are produced by certain species of phytoplankton and macroalgae as a stress response(Andreou et al., 2009; Jagusch et al., 2020). They can also be produced abiotically by reactive oxygen species (ROS) or UV irradiation (Barbosa et al., 2016; Collins et al., 2018).

Oxylipin chemical signaling is largely alleopathic in higher trophic levels of marine systems. Oxylipins produced by diatoms cause teratogenic effects on copepod nauplii and decrease copepod egg hatching success by as much as 80% (Brugnano et al., 2016; Carotenuto et al., 2011; Miralto et al., 1999; Taylor et al., 2007). Copepods from the Adriatic Sea showed increased expression of key stress-related genes (e.g., heat-shock proteins, catalase, glutathione S-transferase, and aldehyde dehydrogenase) when oxylipin concentrations were high (Lauritano et al., 2016). Exposure to micromolar oxylipin concentrations also leads to developmental abnormalities in sea urchins, and alters the expression of genes involved in stress, skeletogenesis, development, and detoxification (Esposito et al., 2020). Oxylipins produced by dinoflagellates exposed to reactive oxygen species (ROS) inhibit the epithelial cells of fish gills (Mardones et al., 2022). Furthermore, experiments on juveniles and adults oysters showed that peroxidation of lipids from the microalgae *Prorocentrum* resulted in hemocyte apoptosis (Wikfors et al., 2008), suggesting that oxylipins are toxic to shellfish.

Thus, phytoplankton-derived oxylipins were the principal analytes investigated as the causative agent impacting planktivorous oyster larvae produced on the Damariscotta River Estuary. Corresponding with spawns in June 2020 and February 2021, we conducted experiments to test whether oxylipins were present in UV-treated water from the Damariscotta River. We also conducted an experiment in July 2020 to determine whether exposing larvae to exogenous oxylipins would result in the same digestive issues.

## 2. Methods

### 2.1 Experiments exploring effects of UV on dissolved organic matter and larval development

In June 2020, an experiment was designed to test for phytoplankton-derived oxylipins in two seawater treatments: treatment 1) seawater was passed through a 1 μm filter as is standard with no UV irradiation and treatment 2) seawater was filtered and then recirculated through the UV sterilizer for 18 h. Dissolved lipids were extracted from each of the two treatments using a solid-phase extraction cartridge. For each water treatment, triplicate Waters™ Oasis HLB cartridges were primed according to the manufacturer’s instructions before 150–200 mL of each treatment was filtered through a cartridge in triplicate. The HLB cartridges were placed in whirl-pak bags, flushed with N_2_ gas, and sealed before shipping on dry ice to UC-Berkeley for analysis.

Additionally, water from the two treatments was used to prepare 15 L buckets for larval bioassays (one bucket per treatment), using 4-day old larvae stocked at a density appropriate to their size, in order to make qualitative larval digestive observations. The larvae used for this experiment were raised in 1 μm filtered seawater for four days prior to the experiment. The larvae were fed according to the standard practices. Larvae were observed by microscopy one and two days following exposure to these water treatments. An in-depth description of the larval culturing techniques and experimental design for larval microscopy and histology can be found in the Supplemental Materials and Methods.

The UV experiment was repeated in February 2021. Triplicate SPE samples were taken for dissolved lipid analysis, as described above. Again, water from the two treatments was used to prepare 15 L buckets for larval bioassays (one bucket per treatment), using 2-day old larvae stocked at a density appropriate to their size.

### 2.2 Dissolved organic matter analysis using untargeted lipidomics

The extraction, chromatographic, mass spectrometric, and chemoinformatic methods are described in full detail within the supplemental methods. Briefly, organics were extracted from filtered seawater using a solid phase extraction (SPE) cartridge. A deuterated internal standard was added during elution of the dissolved lipidome from the SPE cartridge and used along with the volume filtered to normalize the resulting data. Data was collected using reverse phase chromatography paired with high resolution mass spectrometry with data dependent aquistion of ms2. Our chemoinformatic pipeline included annotating features with Lipid and Oxylipin Biomarker Screening Through Adduct Hierarchy Sequences (LOBSTAHS) and MS-DIAL workflows (Ulloa et al., 2025). Please see **figure S1** for our lipidomic workflow, and **table 3** for details of our data reduction through the annotation process.

**Table 3.** Mass spectrometric data reduction using two different branches of the chemoinformatics pipeline present in Supplemental Figure 2.

| XCMS-CAMERA |  |  |
|---|---|---|
| Peaks |  | 1259 |
| Features after removing isotopologues |  | 1085 |
| Manually verified peaks |  | 220 |
|  | LOBSTAHS Annotated | 56 |
|  | FFA + Oxylipins | 19 |
| MS-DIAL |  |  |
| Peaks |  | 3070 |
| ms<sup>2</sup> manually verified | MS-DIAL LipidBlast | 78 |
|  | MS-DIAL Negative Publicly Available Database v17 | 61 |
|  | Waitrose Mammalian Oxylipins Database | 8 |
|  | Edwards Lab CFMID *in silico* marine oxylipin database v1 | 11 |

### 2.3 July 2020 Decatriene + UV experiment to assess larval development and histology

The MSF hatchery designed an experiment to test whether the addition of a common oxylipin, decatrienal, to seawater produced similar effects in the larvae *e.g.* an inability to digest microalgae after ingestion. Owing to the COVID-19 pandemic, only 1,5,9-decatriene was available for purchase during our experiment. It is a fatty acid with the same number of carbons and double bonds as decatrienal. Therefore, we devised an experiment including UV to produce oxylipins from a small fatty acid dissolved in otherwise benign estuarine seawater.

Using embryos from the July 14, 2020, spawn (the final spawning of the season), four experimental seawater treatments were set up in 100 L tanks: 1) 1 μm filtration only (control); 2) 1 μm filtration followed by ∼52 μW/s/cm^2^ UV sterilization; 3) 1 μm filtration followed by the addition of 1,5,9-decatriene to 0.001 mg L^-1^ followed by ∼52 μW/s/cm^2^ UV sterilization, allowing UV irradiation to photo-oxidize the decatriene; 4) 1 μm filtration followed by the addition of 1,5,9-decatriene to 0.001 mg L^-1^, serving as a control for treatment 3.

Embryos were stocked at 20 embryos mL^-1^ in one 100 L tank per treatment, with food introduced on day 1, and then daily as is standard. On days 2, 4, and 6, the water was replaced with water from the appropriate treatment. Larvae from each treatment were observed and photographed on days 2-8. On day 8, larvae from treatments 1, 3, and 4 were fixed in Davidson’s fixative for histological analysis. Unfortunately, not enough larvae from Treatment 2 remained for fixation.

## 3. Results

### 3.1 The effects of UV on oyster larvae during the 2020 and 2021 hatchery seasons

The larval bioassay in June 2020 show UV sterilization of filtered seawater led to symptoms of larval digestive syndrome (**figure 5**). Because these larvae had spent 2-4 days growing in favorable conditions prior to treatment for two days, they were more normally developed than those seen in figure 1 and figure 2 from February 2020. However, the larvae exposed to Treatment 2 (1 μm filtration & recirculated UV) were significantly smaller than Treatment 1 (1 μm filtration only) and lacked umbo development at the hinge. The larvae in Treatment 2 did not have large amounts of undigested algae in their stomachs (as in **figure 2**), but their digestive glands were significantly lighter in color than the larvae in Treatment 1, indicating less digested material.

**Figure 5:**
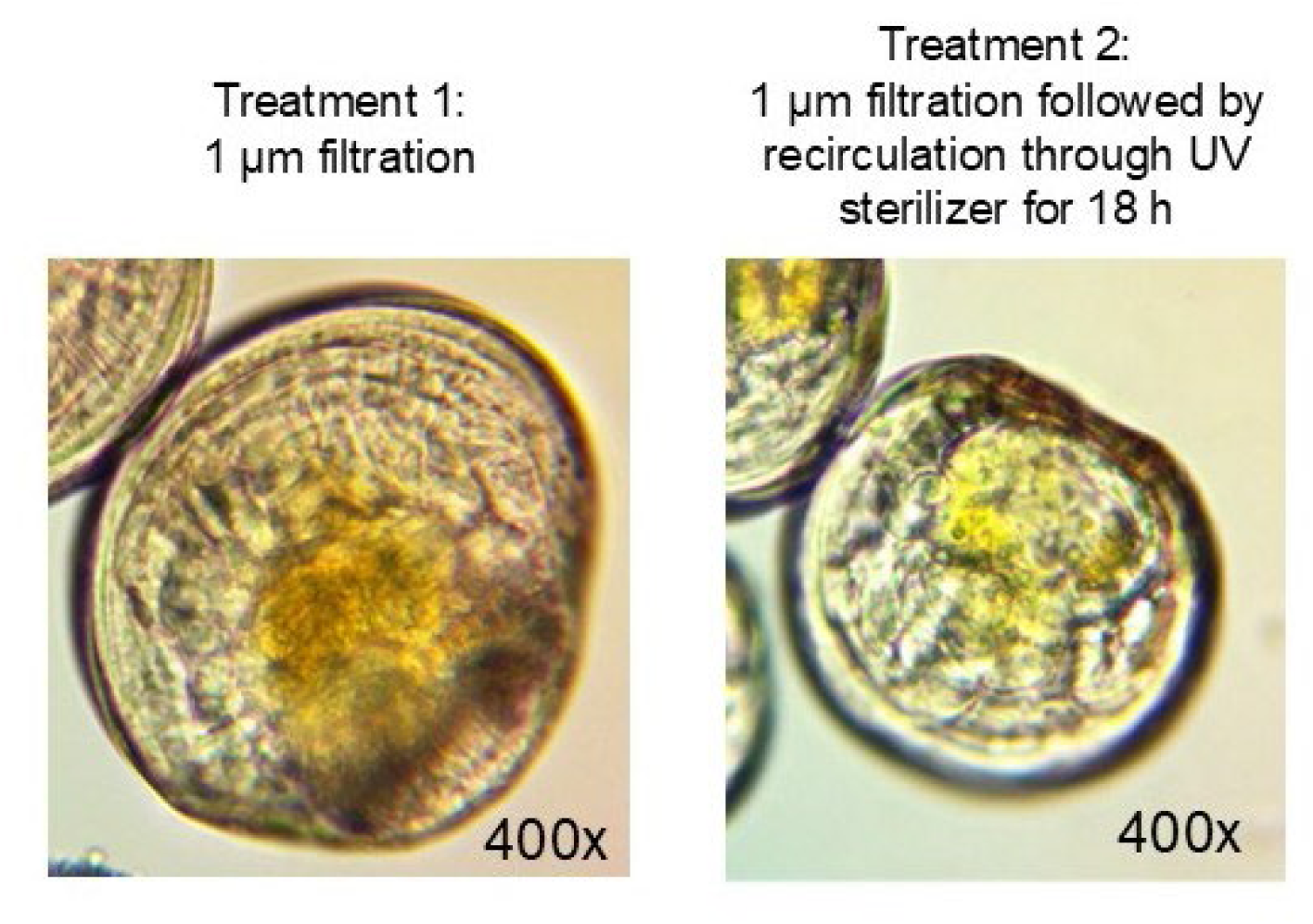
Microscopy of day 6 oyster larvae from the June 2020 UV experiment, after two days exposure to two different water treatments, parallel oxylipin analysis is presented in Figure 6. Prior to treatment exposure, all larvae were raised in 1 μm-filtered seawater. Because all larvae had spent four days growing in favorable conditions prior to treatments for two days, larvae are all more developed than in other figures of affected larvae. However, the larvae exposed to Treatment 2 (recirculated UV) were noticeably smaller without the umbo development at the hinge seen in larvae from Treatment 1. The larvae in Treatment 2 did not have large amounts of undigested algae in their stomachs as previously observed, but their digestive glands are significantly lighter in color (a symptom of the digestive syndrome) than larvae in Treatment 1. Images are at the same scale and were taken under 400x magnification.

### 3.2 Annotation of the Dissolved Lipidome from the Natural Seawater UV-experiments

In total, 220 features were identified in the dissolved lipidome from experiments exposing filtered estuary water to UV. (**figure 6a; table S1a**). Of these 220 features, a boutique database, LOBSTAHS, assigned putative annotations to 56 compounds (**figure 6b**). Eleven of these features had competing lipid assignments; specifically, there was overlap in the chemical formulas and exact masses of free fatty acids with oxidized polyunsaturated aldehydes (PUA+1 oxygen) of the same chain length and double bond equivalent, odd chain mono-acyl glycerides (MAGs) with oxidized fatty acids (+2 oxygens), oxidized betaine lipids with betaine-like lipids, and oxidized lyso phosphytidylcholine (lysoPC) with oxidized lyso phosphytidylglycerol (lysoPG). Thus, both annotations appear for these features and the names are italicized to indicate a lower confidence level of assignment (**figure 6c, f, 8b, 9b**).

**Figure 6.**
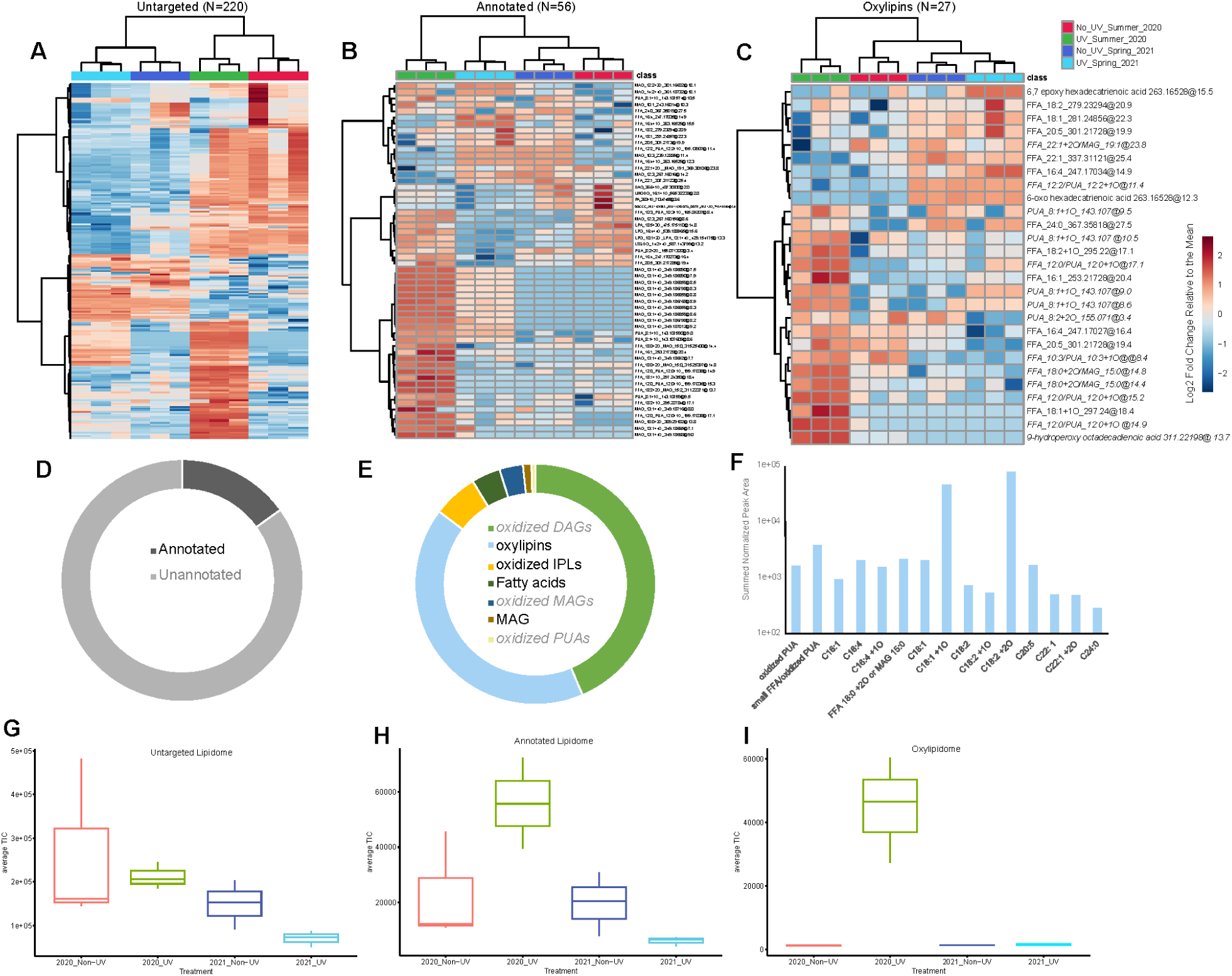
Untargeted and targeted lipidomes. A) heatmap of the relative abundance of untargeted features in Summer 2020 and February 2021 in UV-treated triplicates and nonUV control triplicates. (220 features across twelve samples). B) Heatmap of the relative abundance of features in the Annotated Lipidome from UV experiments in Summer 2020 and February 2021. (56 features across twelve samples.) C) Heatmap of the relative abundance of features annotated as oxylipins from UV experiments in Summer 2020 and February 2021. D) Percentage of the untargeted lipidome total ion chromatogram (TIC) (i.e. the summed peak area of detected features) that was annotated with LOBSTAHs versus unannotated. E) Percentage of the LOBSTAHs Annotated Lipidome TIC represented by each lipid subclass. Compounds in grey italics are less certain putative annotations. The oxidized MAGs we attempted to annotate separately. F) Percentage of the oxylipidome TIC represented by each fatty acid chain length, oxidation state, and oxylipin subclass (i.e. summed peak areas over regioisomers). Average TIC in each triplicate for G) the untargeted lipidome, H) the annotated lipidome, and I) the oxylipidome.

In terms of the total integrated peak area or the total ion chromatogram (TIC), 15% of the TIC was annotated by LOBSTAHs database, which is populated with more than 90,000 individual lipids (Collins et al. 2016; **figure 6d**). Of the LOBSTAHS annotated lipidome, 42% of the TIC was annotated to the oxylipin subclass, 5% as free fatty acids, and 44% was annotated by LOBSTAHs as a single [M-H]^-^ adduct of an oxidized diacylglycerol lipid with 26 carbons, six double bond equivalents, and an extra oxygen moiety in addition to the diacylglycerol backbone (abbreviated DAG 26:6 +1O; **figure 6e**). Further inspection of this feature’s ms^2^ spectra revealed that it was likely an [M+Cl]^-^ adduct of a parent ion with an m/z of 451.3287 [M-H]^-^ (**figure S2**). Our adduct hierarchy model LOBSTAHs includes [M+Cl]^-^ adducts, so this compound is isobaric with DAG 26:6 +1O, but represents another subclass of lipids that was not included in our database. We searched for the monoisotopic mass of the parent mass, M, against the PubChem database and found eight putative chemical formulas representing over 488 potential isomers with 2.5 ppm mass accuracy (**table S2**).

Comparative lipidomics of the untargeted lipidome (Section 3.3) revealed the importance of five unannotated features and five isomers annotated as a monoacylglycerol with 13 carbons, one double bond equivalent, and four extra oxygens (MAG 13:1+4O; **figure 7b**). These compounds, along with the oxidized DAG and oxylipins that dominated the annotated lipidome, were further investigated by querying the ms^2^ fragmentation of these features against the MS-DIAL publicly available negative library (v17) and boutique databases with predicted fragmentation of common diatom oxylipins and iterative fragmentation of common mammalian oxylipins (Ding et al., 2021; Djoumbou-Feunang et al., 2019; Edwards et al., 2024; Tsugawa et al., 2019; Watrous et al., 2019).

**Figure 7.**
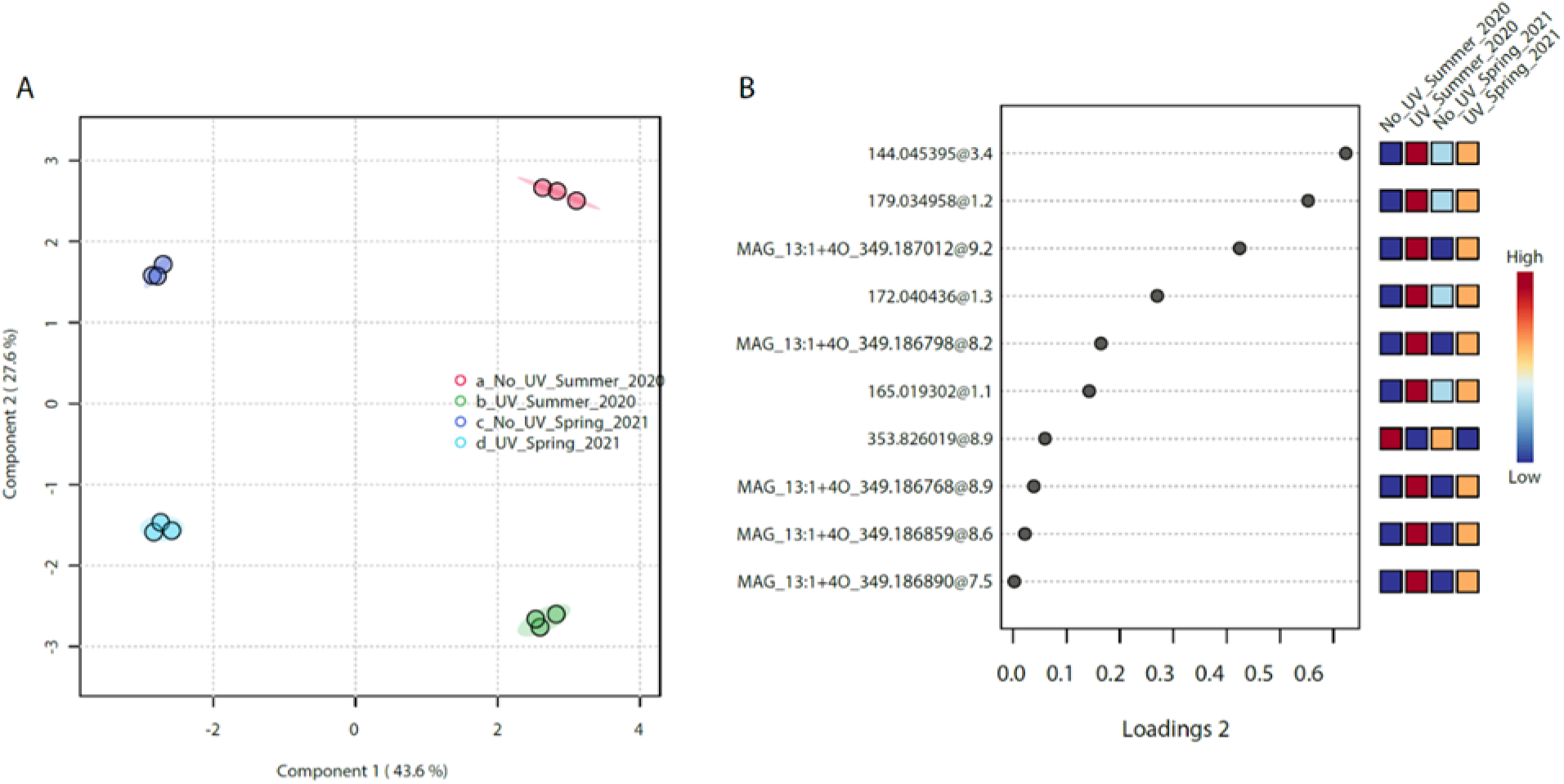
Compounds within the untargeted lipidome associated with UV treatment. A) sPLS-DA sample loading of untargeted lipidomes from waters feeding an oyster hatchery in 2020 and 2021 with and without exposure to UV. B) sPLSDA loading of compounds that were assigned to component 2, which is most distinguished the non-UV treatment from the UV treatment. Averaged relative abundance of compounds across each triplicate displayed in the heatmaps.

In conjunction with our untargeted analysis, four of the five most significant unannotated features were putatively identified by matching their fragmentations to publicly available spectral libraries (**figure 7b**; **table 4**). The compound most significantly “upregulated” by UV in both 2020 and 2021 was putatively identified as a 2-hydroxy quinoline antibiotic (m/z = 144.0454 RT-3.4 min; **table 4**).

**Table 4.**
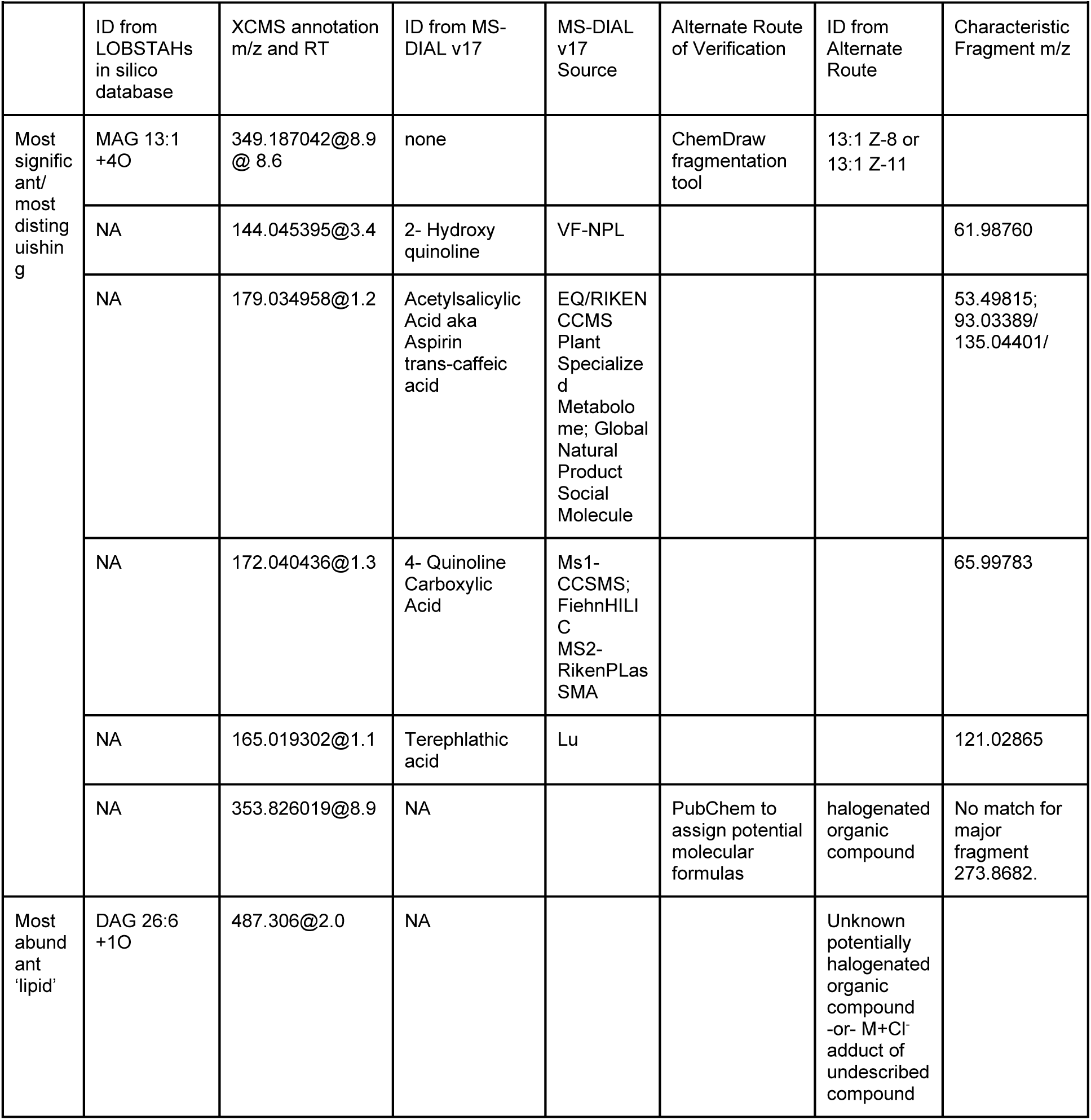
Annotation of the most significant compounds in the manually verified lipidome using MS-DIAL and other methods of ms2 verification, and reassessment of annotation given to the most abundant feature in the LOBSTAH annotated lipidome.

The next most significantly ‘upregulated’ compound (m/z= 179.034958 RT 1.2) had ms^2^ fragments matching both acetylsalicylic acid, more commonly known as aspirin, and trans-caffeic acid. The fourth most upregulated compound was identified as 4-quinoline carboxylic acid based on the ms1 match (m/z 172.0404 RT-1.3). It did not fragment into the characteristic m/z = 128.049 ion found in the Fiehn and CCMS ms^2^ libraries. However, ms^2^ contained a 65.99783 fragment associated with the 4-hydroxy quinolone observed in the RIKEN PlaSMA library. Thus, this compound was likely a quinoline.

The sixth most significant unannotated compound was 165.0194 compound at 1.1 minutes which was identified as terephthalic acid, a plasticizer. The seventh most significant compound, 353.8260 @ RT-8.9 was actually “downregulated” by UV, but its fragmentation did not match any publicly available databases. We attempted to assign an elemental formula to the feature, assuming that we observed an [M-H]^-^ adduct, we found four potential chemical formulas under 2.5 ppm, all containing halogenated compounds, and many of the isomers listed in PubChem have patents. (**table S3**).

### 3.3 UV increases the dissolved concentrations of compounds across untargeted, annotated, and oxylipin-only lipidomes

The untargeted lipidome provides the widest analytical view of which features are the most different, without introducing database bias. To give high abundance and low abundance compounds equal weight, we looked at the relative abundance of each feature compared to the mean abundance across all treatments, represented as a log fold change. This analysis revealed that the 2020 dataset was richer in dissolved organic compounds than the 2021 dataset (**figure 6a**).

An overview of the 220 features observed in the lipidome showed a general increase in the relative abundance of many organic compounds when 1 μm filtered water was exposed to UV (**figure 6a**), but the summed peak area was not significantly different (**figure 6g).** A total of 177 of 220 compounds were significantly different between the UV and non-UV treatments (ANOVA, FDR <0.05, **Supplemental Report 1**). When we considered only the annotated lipids, the overall TIC was significantly higher in the UV treatment, especially with respect to the oxylipins and free fatty acids constituting the oxylipidome (**figure 6j-i**). Further analysis of the annotated lipids and oxylipidome separately revealed a similar trend, with an increase in the relative abundance of many compounds in the UV treatments.

The samples clustered according to year and treatment when Ward hierarchical analysis was applied to the entire untargeted lipidome (N=220; **figure 6a**). Hierarchical clustering of the LOBSTAHS annotated lipidome clustered the samples by treatment and not by year (N=56; **figure 6b**). The LOBSTAHS annotated lipidome of the UV 2021 samples was more similar to that of the UV 2020 samples than to the non-UV samples from 2021. Similar analysis of the oxylipidomes clustered the samples by toxicity with the UV-treated samples from 2020, where the oyster larvae were most impacted, clustering separately from the less-toxic non-UV 2020 samples which were more similar to the far less toxic 2021 samples (N=27; **figure 6c**). With respect to both the annotated lipidome and oxylipidome, UV-treating the water increased the relative abundance of compounds in the dissolved organic pool in both 2020 and 2021; however, the response in 2020 was more intense (**figure 6b,c,h,i**).

#### 3.3.1 Oxidized lipids associated with UV in the LOBSTAHs Annotated Lipidome

Machine Learning (Random Forest) and multivariate analysis (sPLSDA) distinguished the treatments based on their triplicate lipidomes (**figures 7a, 8a, 9a**) and allowed us to identify the specific oxylipins, free fatty acids, and oxidized lipids that were abundant in the oyster hatchery waters when UV pretreatment was applied (**figure 6b**; **figures 7**–**9**). When considering the sPLSDA used to analyze the untargeted lipidome, Component 1 separated the samples based on year and Component 2 separated the lipidomes based on UV treatment. The lipidomic features used to derive Component 2 were higher when UV was applied, and are presented in **figure 7b**, in descending order of importance. The average relative abundance of each compound is shown in the heatmap. Five isomers of MAG 13:1 +4O are among the ten most important annotated compounds in the untargeted lipidome (**figures 7b).** Random forest analysis best captured the differences between the treatments when considering the LOBSTAHS-annotated lipidome alone (**figure 8a, c**), and the same five MAG 13:1+4O compounds were among the most important features (**figure 8b**). The out-of-bag error was zero, which indicated that the random forest classified each lipidome by treatment perfectly (**figure 7c**). The annotated lipidomes were so distinct that fewer than 50 trees were required to converge upon an answer (**figure 7a**).

**Figure 8.**
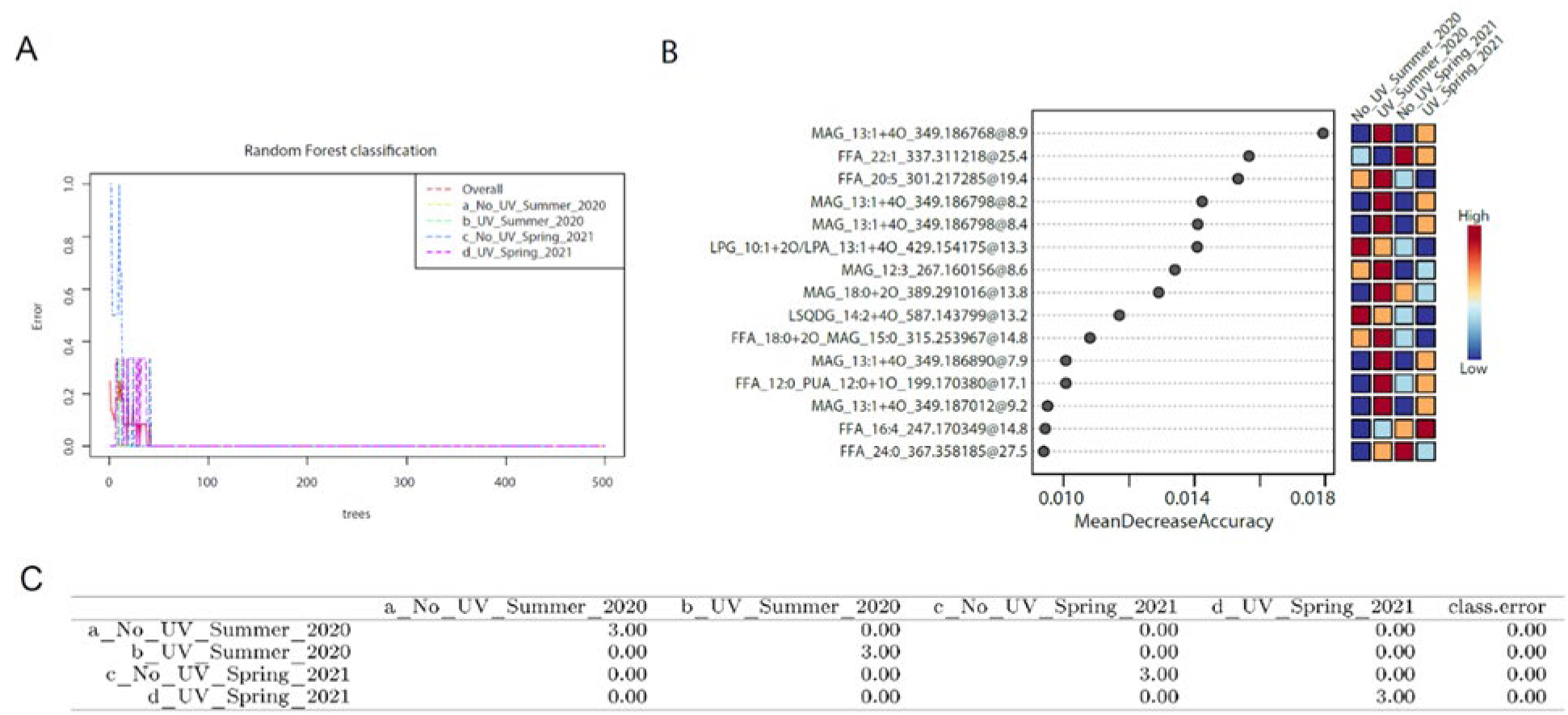
The annotated lipidome was distinct between UV and non-UV treatments in 2020 and 2021. A) Random Forest analysis of the annotated lipidome results showing error with the number of trees. B) Heatmap of the most significant compounds structuring the random forest walk ordered by mean decrease accuracy. The average relative concentration of each compound across biological triplicates is shown in the heatmap. Red hues indicate higher relative abundance and blue hues indicate a lower relative abundance. C) Table with out-of-bag error for each sample class (No-UV and UV Summer 2020, No-UV and UV Spring 2021) showing perfect classification (out of bag error of 0) of triplicate lipidomes.

**Figure 9.**
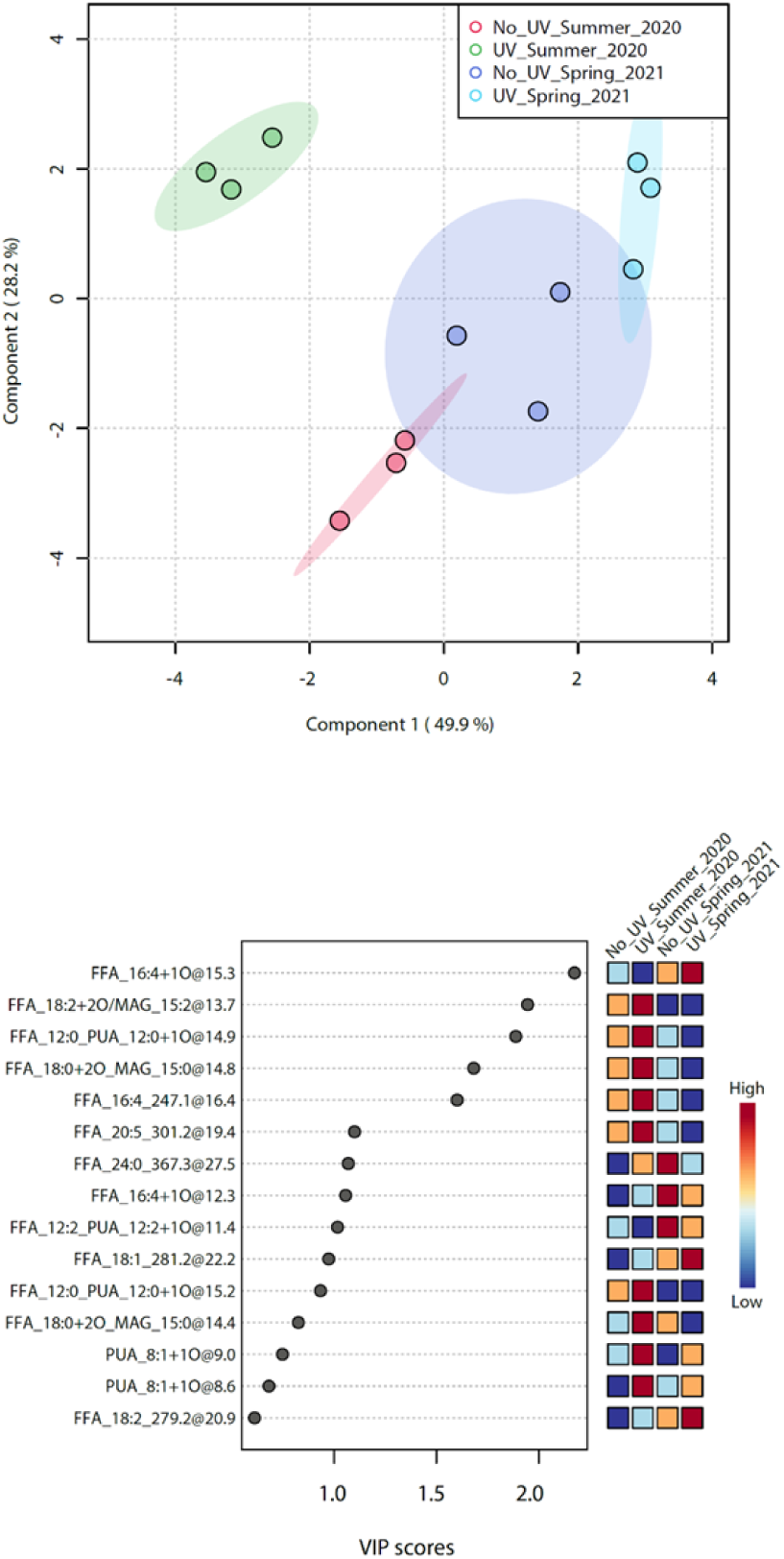
Partial Least Squares Differential Analysis (PLSDA) of the Oxylipidome (N=27 compounds). A) PLSDA loadings of the triplicate sample types on Component 1 and 2, representing 49.9% and 28.2% of the variability in the meta-lipidome, shows UV treatment caused a significant shift in the dissolved oxylipins and free fatty acid pool. B) Eleven compounds were considered significant features for differentiating the lipidome classes with VIP (variable importance projection) scores greater than 1 for Component 1. The heatmap shows the average relative concentration of each compound in each treatment and year. Red hues indicate a higher relative abundance and blue hues indicate a lower relative abundance.

In our dataset, two MAG 13:1 +4O isomers at retention times of 8.2 and 8.6 minutes were abundant enough to produce ms2 fragmentation (**figure S3a-d**). These features in our dataset did not match those of any compounds represented in the negative MS-DIAL database, which is a compilation of both authentic ms2 spectra and in silico libraries across publicly available spectral libraries. Therefore, we explored the fragmentation of four isomers of MAG 13:1 +4O with double bonds at the 2-, 5-, 8-, 11-carbons using the fragmentation tool in ChemDraw and found characteristic fragments for either 13:1 Z-8 or 13:1 Z-11 isomers (143.10649 ms2 fragment observed at both retention times; **figure S3e**). The other isomers that varied significantly with UV were not abundant enough to fragment, but we interpreted them as other isomers based on the retention time of authentic reference standards for a mix of 18:1 oxylipin isomers (**figure S4**). We expect that the earlier retention time is the Z-11 isomer, the latter isomer is the Z-9 isomer, and the other isomers at 9.2 are likely the 2-, 5-isomers of the MAG 13:1 +4O compound. The relative abundance of these isomers increased with UV in 2020 and 2021, but the relative increase in 2020 was approximately four times higher than that in 2021 (**figure S5**).

Three compounds, MAG 12:3, the free fatty acid (FFA) 20:5, and compound co-annotated as an oxidized PUA and FFA 12:0, responded similarly with higher relative concentrations in the UV-treated triplicates and higher overall concentrations in 2020 compared to 2021 (**figure 8b**). The very long-chain FFA 22:1 was distinctive for the experiments in 2021 decreasing upon UV irradiation, whereas that of 16:4 increased.

#### 3.3.2 Effect of UV on the suite of oxylipins and free fatty acids (i.e. the oxylipidome)

The oxylipidome was significantly different between treatments, specifically in June 2020. Most readily, we can see that reflected in the integrated TIC (**figure 6i**), where the average normalized peak area of oxylipins and free fatty acids was much higher after UV pretreatment in 2020 than in the other treatments (>3 orders of magnitude). Of the 27 compounds identified as either oxylipins or free fatty acids, 19 compounds were significantly different between treatments using one-way ANOVA (**figure S6**). Fisher’s ad hoc test was performed. Most of these compounds were more abundant in the UV samples than in the non-UV-treated samples. **Table S4** provides a complete list of treatments that are significantly different from each other.

Multivariate analysis (PLS-DA) agreed with the parametric analysis, pinpointing a suite of oxylipins that were the most distinctive (**figure 9b**). The June 2020 UV treatments had high concentrations of C18:n oxylipins and medium-chain fatty acids (**figures S6k-o** and **8b**). In contrast, the February 2021 UV treatment had a free fatty acid and +1O oxylipin from the 16:4 biosynthesis pathway, as well as some distinctive very long-chain fatty acids and oxylipins (**figures S6q-t** and **9b**). According to the variable importance in projection (VIP) score from the PLSDA, the 18:2 +2O fatty acid was the most important feature in the UV Summer 2020 treatment, and 16:4+1O was the most important feature in the UV February 2021 experiment (**figure 9b**).

We used ms^2^ to obtain a more reliable putative annotation for these oxylipins. Fragmentation of 18:2 +2O across all samples suggested that the compound was 9-hydroperoxy octadecadienoic acid (**figure 10a-c**). Fragmentation of 16:4 +1O at RT-15.3 integrated across all samples (**figure 10g-h**) and from just February 2021 (**figure 10i-j**) suggests that it is an (5,6)-epoxy hexadecatetraenoic acid (**figure 10k**), although this annotation is not as confident because the compound was low in abundance and did not fragment well. This is evident by the even nature of the lower abundance of ms^2^ fragments, which denotes a decrease in our instrument mass accuracy when ms^1^ is ∼5E4 (**figure 10h, j**). There was another isomer of 16:4 +1O (RT=12 min), which was more abundant across all samples and matched the fragmentation of 6-oxo-hexadecatetraenoic acid (**figure 10d-f**), which strengthens the putative annotation of the 16:4 +1O isomer at 15 min as a product of a 6-lipoxygenase biosynthetic pathway.

**Figure 10.**
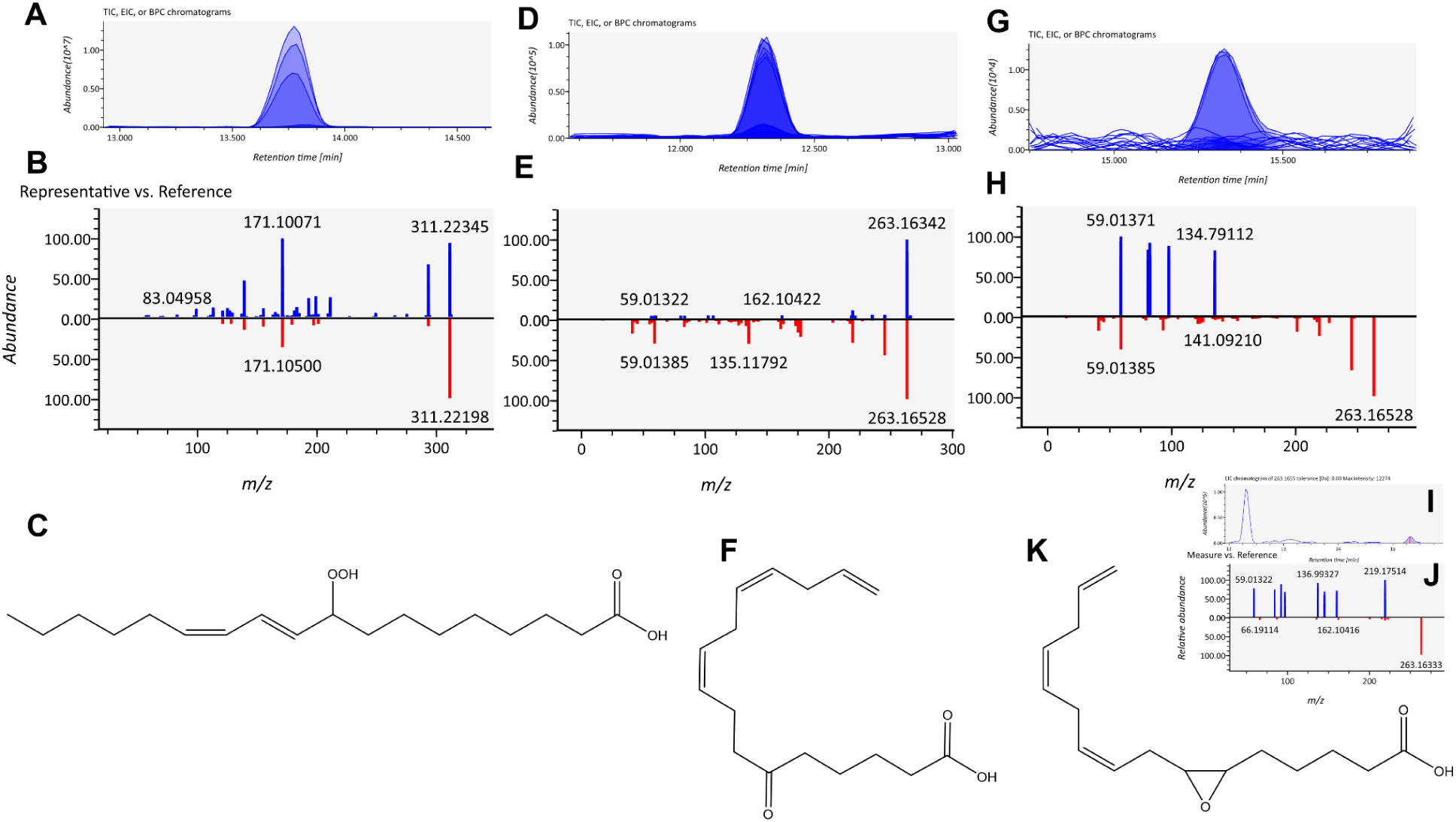
Annotation of non-volatile oxylipins. A-B) Extracted ion chromatogram for m/z 311.22345 @13.7 minutes and ms2 fragmentation compared to in silico fragmentation of C) 9-hydroperoxy octadecadienoic acid. D-E) EIC and ms2 for m/z 263.16528 at 12.3 minutes compared to in silico fragmentation for F) 6-oxo hexadecatrienoic acid. G-H) EIC and ms2 for m/z 263.16528 at 15.3 minutes compared to in silico fragmentation for K) 6,7-epoxy hexadecatrienoic acid. Because only one ms2 fragment matched the reference in H, just the spring 2021 samples were compared in I and J showing there may be different isomers with different fragmentation patterns eluting at 15.5 minutes in the samples from various years and treatments.

### 3.4 Directly testing the effect of oxylipins on oyster larval development

A commercially available oxylipin precursor, decatriene, was used to challenge the recently spawned larvae in July 2020. 1,5,9-decatriene was added to 1 μm-filtered seawater at a concentration of 0.001 mg L^-1^. The decatriene-seawater solution was passed through a UV sterilizer to photooxidize decatriene to decatrienal. Larvae exposed to decatriene + UV seawater for 6-8 days had symptoms of the larval digestive syndrome that were seen in 6 d larvae from the February 2020 spawn (**figure 11a, d**) with similar debris in the gastric lumen (**figure 11f, h**). Notably, the addition of decatriene to 1 μm filtered seawater without UV irradiation exposure did not produce symptoms of larval digestive syndrome (**figure 11e,i**). Nor did UV treatment of filtred seawater during this spawn (**figure 11c**), showing that the added decatriene was UV-oxidized to deleterious compound and not the exogenous lipids in the system at the time. The innocuous nature of UVed seawater in June 2020 but toxicity to larvae in Febrary 2020 also reiterates the seasonal variability in the estuary dissolved organic matter load.

**Figure 11.**
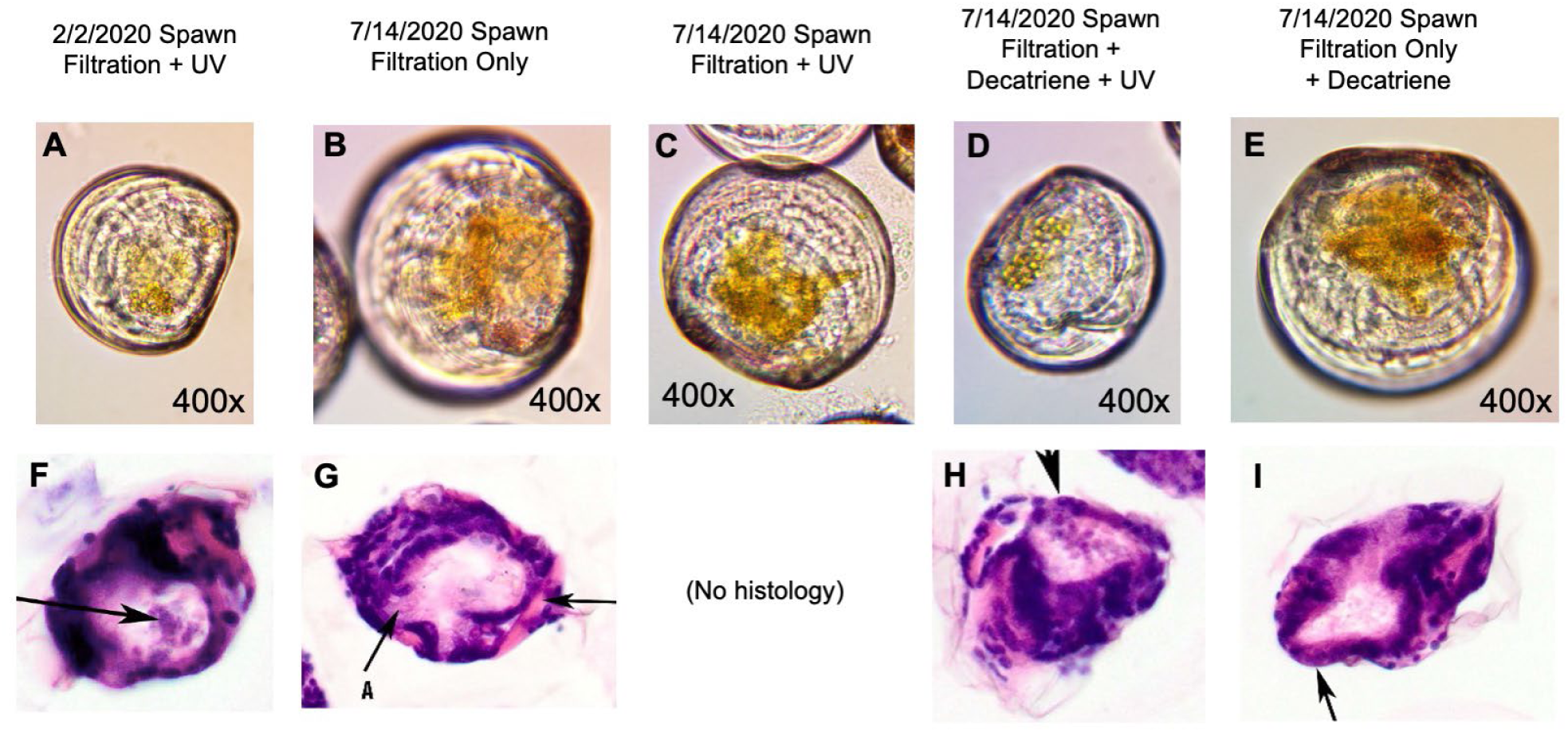
Micrographs (A, B, C, D, E) and histological images (F, G, H, I) of 6-day old (A, B, C, D, E, F) or 8-day old (G, H, I) oyster larvae exposed to various treatments. All micrographs were taken at 400x magnification and are shown at the same scale. The larvae from a February 2, 2020 MSF production spawn using 1 μm filtration and ∼52,000 μW/s/cm^2^ UV irradiation (A, F) show very little digested algal material in the digestive gland, typical of the larval digestive syndrome (A) and debris in the gastric lumen (F). Larvae from a 7/14/2020 experimental spawn exposed only to 1 μm filtration only (B, G) are relatively larger and show a dark golden color of digested algal material in their digestive gland (B) and no debris in the gastric lumen (G). Larvae from a 7/14/2020 experimental spawn exposed to 1 μm filtration and 52,000 μW/s/cm^2^ UV irradiation (C) are a similar size to those exposed to filtration only (B) and have a similar dark color of digested algal material in the digestive gland. Larvae from a 7/14/2020 experimental spawn exposed to 1 μm filtration with 0.001 mg L^-1^ decatriene added, followed by 52,000 μW/s/cm^2^ UV irradiation (D, H) were most similar to those from the 2/2/2020 production spawn, showing symptoms of the digestive syndrome (D), as well as debris in the gastric lumen (H). Larvae from a 7/14/2020 experimental spawn exposed to 1 μm filtration with 0.001 mg L^-1^ decatriene added, with no UV irradiation (E, I) were most similar to larvae from that spawn exposed either to filtration only (B, G) or filtration and UV irradiation (C), with a larger size and digested algal material in the digestive gland (E) and no debris in the gastric lumen (I).

## 4. Discussion

The digestive syndrome observed in larval oysters at Mook Sea Farm in 2020 and 2021 was linked to UV sterilization procedures and the generation of oxylipins and other allelopathic compounds (**figures 6**–**10**). When the UV step was removed, the larval digestion and development returned to near-normal levels. Exposure of a commercially available oxylipin precursor to UV recreated the symptoms and histological patterns observed at the initial onset of digestive syndrome (**figure 11**).

### 4.1 The diversity of features detected and lipids annotated in estuarine waters

Over 220 features were identified in the meta-lipidome from the UV treatment experiments in 2020 and 2021 (**figure 6a, table S1**). This untargeted approach showed that there were more lipid compounds dissolved in the waters feeding the Mook Sea Farm oyster hatchery in 2020 than in 2021, and UV treatment increased the abundance of a subset of compounds (**figure 6a**). We putatively annotated 56 of these compounds as lipids using the LOBSTAHs database (**figure 6b,e**), which matches ms1 and adduct formation to an *in silico* database generated for the most abundant lipid subclasses in marine systems (Collins et al., 2016).

Lipids are typically considered to be hydrophobic. However, smaller lipids with oxygen are more water-soluble (polar), and these are precisely the type of compounds that we observed. FFA, oxylipins, oxidized monoacylglycerides (MAGs), and IP-DAGs were the dominant subclasses of annotated lipids. Glycolipids and phospholipids are precursors for oxylipin production in *Skeletonema* and *Thalssiosira* diatoms (Cutignano et al., 2006; d’Ippolito et al., 2004; Fontana et al., 2007). Very few intact lipids were annotated in the dissolved lipidome; however, the oxidized lyso membrane lipids (PG, SQDG, MGDG, or PA) and oxidized betaine-like lipids annotated in the lipidome may be oxylipin precursors or co-products of oxylipin production **figure 6b**).

### 4.2 Overall increase in dissolved lipid-like molecules in response to UV

In 2020, the natural waters of the Damariscotta Estuary were more lipid-rich and exhibited a stronger response to UV than in 2021 (**figure 6a-c,g-i**). Some of the compounds match oxylipins produced by phytoplankton macroalgae, and some are pharmaceuticals that are natural products from higher plants (**figure 6c,f** and **table 4**). Therefore, we believe that these compounds are from a mixture of marine and terrestrial sources, perhaps bits of detritus (fragmented cells, fecal material, decaying organisms) and small living cells that pass through the 1 µm filter. Future research efforts will focus on collecting eDNA while profiling oxylipins and other metabolites.

### 4.3 Oxidized lipids, anti-inflammatory metabolites, and antibiotics concentration increase with UV

We initially set out to investigate known marine oxylipins in the Damariscotta River Estuary, namely polyunsaturated aldehydes and non-volatile oxylipins (NVOs, sometimes referred to as LOFAs), which we linked to UV radiation. In fact, the difference between UV and non-UV was more significant for oxylipins as a class than for annotated lipids or the untargeted lipidome (**figure 6g-i**).

Our untargeted lipidomic approach opened the analytical window for unknown compounds that were also captured by solid-phase extraction. Of note are the ten most significant compounds for distinguishing between UV treatments across the two years (**figures 7b, 8b).** Five of these compounds are isomers putatively annotated as MAG 13:1 +4O based on the ms^1^ (m/z = 349.1865) matching the LOBSTAHs database with 2.5 ppm mass accuracy and ms^2^ matching predicted fragmentation of positional isomers of MAG 13:1 +4O (**figure S3**). The other five compounds were not annotated using the LOBSTAHS Database, which primarily contain lipids found in marine phytoplankton. We annotated four of these five compounds as two quinolone metabolites, a plasticizer, and a salicylic acid metabolite. A more detailed discussion of the bioactivity and potential sources of statistically significant compounds can be found in Section 4.3.3.

#### 4.3.1 Phytoplankton oxylipins with known allelopathic effects linked to UV

Within the oxylipidome, we observed that UV irradiation led to an increase in oxylipin abundance, especially in 2020, when there was a much higher absolute abundance of oxylipins (**figure 6i**) and a significantly different dissolved lipidome structure (**figure 6b-c** dendrograms). A series of C18 oxylipins was most abundant in 2020, whereas C16:4 oxylipins were more abundant in 2021 (**figures 6c, 9b, and 10**). C18 oxylipins have been well described in macroalgae and freshwater cyanobacteria (Andreou et al., 2009) giving us two potential sources for investigation. C16:4 oxylipins are specific to diatoms and have been shown to decrease copepod reproduction in the Adriatic Sea (Lauritano et al., 2016). Experiments with UV-oxidation of fatty acids suggest similar C18 and C16 oxylipins are amongst the causative agents of the oyster larval digestive syndrome described in the introduction (**figure 11**). We will also consider other sources of oxylipins in Sections 4.3.2 and 4.3.3.

##### 4.3.1.1 Oxidized lipids stunt growth and inhibit digestion

We conducted experiments with UV-oxidized decatriene and found that it produced the same larval digestive syndrome and histological response as was seen in February 2020 UV-treatment (**figure 11**). Inhibition by oxylipins is well documented in a number of species, and these compounds are often signaling molecules in reproductive pathways (*e.g.,* prostaglandins in the human birthing process, teratogenic effects on copepod nauplii (Caldwell, 2009), decreased egg hatching success (Miralto et al., 1999), and developmental abnormalities in sea urchins (Esposito et al., 2020). Similar to copepods, we did not observe any deleterious effects on the adult oysters in the Mook Sea Farm, as they were able to spawn without ill effects. However, the lingering effects on setting competency (**table 2**) and non-lethal shell deformities (**figure 3**) outlined in the introduction are further causes for concern. Delayed competency for setting increases production costs for oyster hatcheries and reduces the reliability of the standard processes.

#### 4.3.2 Highly oxidized monoacylglycerol lipid significantly correlated with UV treatment

At first glance, the MAG 13:1 +4O isomers linked to UV (**figures 6b, 7b**) may seem unlikely because small odd-chain fatty acids are rare in nature. Typically, the smallest monosaturated fatty acid microalgae biosynthesize is 16:1 although myristic acid can be a minor component of the membrane in some taxa (Jónasdóttir, 2019). Obscure references to C13:1 lipids in the literature include, C13:1 fatty acids identified in lipid extracts from cabbage (Peng, 1973), the lipopolysaccharides (LPS) of gram-negative bacteria that are associated with periodontal disease(Kumada et al., 1997), and in the LPS-like lipids of pathogenic gram-negative bacteria *Leptospira interrogans* which infect pets (Shimizu et al., 1987). Furthermore, (E-11) tridecaenoic acid C13:1 was found to inhibit desaturase activity in the production of sex pheromones by moths in a dose-dependent manner (Rodríguez et al., 2004). Trans-2-tridecanoic acid interacts with the sialic acid system to inhibit ulcers in rat guts (Mimura et al., 1983).

Alternatively, these isomers may represent the degradation products of IPLs that are converted into lyso fragments by losing one fatty acid and a MAG by losing the polar headgroup on the lipid. While this can be a biotic process through lipase activity, oxylipin biosynthesis, and membrane lipid catabolism (access to P in the phospholipid head groups), highly oxidized short-chain MAGs may be the abiotic byproducts of UV-oxidation. The large number of isomers supports UV degradation of lipids in the dissolved pool, because many different species of lipid would be degraded, leading to different oxidized isomers of MAG 13:1. This represents a new set of oxylipin analytes that should be studied with respect to their chemical ecology and impact on shellfish reproduction.

#### 4.3.3 Compounds annotated as microplastics, pharmaceuticals, and natural products released by UV sterilization

The MS-DIAL database was instrumental in assigning putative annotations to the unknown compounds from the initial pipeline, helping us putatively annotate the most distinctive features distinguishing UV from non-UV samples in the untargeted lipidome as 2-hydroxy quinolone, aspirin, 4-quinoline carboxylic acid, terephthalic acid, and an unknown compound that we believe is a halogenated organic compound (**figure 7b**; **table 4**). This range of compounds is representative of the general spread in compound classes annotated with MS-DIAL (**table s4**). Halogenated organic compounds, like the unknown, have been linked to phytoplankton in the Northeastern Atlantic and can volatilize from the ocean to the atmosphere, where they are greenhouse gases (Hepach et al., 2014; Shi & Wallace, 2018). Marine diatoms produce halogenated small fatty alkenes via the same enzymatic pathways used to produce oxylipins, whereas macroalgae produce cyclical C20 halogenated oxylipins (Kousaka et al., 2003; Wichard & Pohnert, 2006).

The increase in terephthalic acid, a plastic monomer found in water bottles and synthetic clothing, is likely due to the degradation of microplastics by UV. Natural waters are known to contain hundreds of dissolved pollutants, including pharmaceuticals, that enter the environment through municipal wastewater treatment streams (Joss & Ternes, 2007). In this dataset alone, we identified 15 compounds that were pharmaceuticals, such as gemfibrozil, a medication used for lipid regulation in human blood serum; 4-Hydroxyquinoline, an antibiotic and over-the-counter skin lightener; tetradecylsulfate, an anti-sclerosing agent used to treat varicose veins; and triamcinolone, a steroid used to treat rheumatic disorders and allergies (**table s4**).

However, the question remains, why does the concentration of anthropogenic compounds dissolved in water increase with UV exposure. Organic pollutants are known to adsorb onto microplastics because of their hydrophobicity and leech from microplastics as polymers degrade (Ding et al., 2021). The terephthalic acid signal corroborates the hypothesis that UV leeched these compounds from microplastics into the water (< 1 μm in this case). Alternatively, bioaccumulated pollutants may be released when the small cells (< 1 μm) remaining in the water are exposed to UV radiation.

A third potential source for some of the simpler pharmaceuticals, such as aspirin and salicylic acid, and potentially the two quinolone compounds, 2-hydroxy quinolone and 4-quinoline carboxylic acid, would be phyto-detritus (marine or terrestrial) or living cells smaller than 1 μm respond to UV pretreatment with their own anti-inflammatories. Both aspirin and quinolone antibiotics are natural products first extracted from terrestrial plants (Dias et al., 2012). Salicylic acid, which can be used as a chemical exfoliant in skincare, is a secondary plant metabolite of the same pathway that produces aspirin. There were several other natural products putatively identified with MS-DIAL, including 6-Gingerol, Cucumenol, rotenone, loganic acid, and onopordopicrin, which are potential pharmaceuticals, but also occur as natural products that could have been produced *in situ* (**table s4**).

The oxylipin chain lengths suggest production by a mixed community. Considering the bulk of oxylipin literature, the presence of 16:4 oxylipins and C20:5 fatty acids (isomers of the essential fatty acid EPA) suggest diatoms as sources in February 2021. The presence of C22 oxylipins may indicate diatoms such as *Lepidocylindrus* (Nanjappa et al., 2014), but dinoflagellates, haptophytes, and cryptophytes traditionally produce more C22 fatty acids than diatoms overall (Jónasdóttir, 2019). The 18:n oxylipins associated with June 2020 suggested bacteria or fungi as sources (Andreou et al., 2009; Martínez et al., 2010), although 18:n oxylipins were a minor component of the lipidome in experiments exploring grazing and viral infection of planktonic marine diatoms (Johnson et al., 2020; Ulloa et al., 2025).

### 4.4 Seasonal variability in dissolved lipidome

The different suites of oxylipins in summer 2020 and February 2021, as well as the higher concentration of dissolved organic lipids in summer compared to February, suggest seasonal variability that likely correlates with photosynthetic biomass, which is consistent with a time series study of NVOs in the Adriatic (Russo et al., 2020). In addition to natural seasonal variability, these differences in baseline dissolved lipidome between the two years could also be linked to differences in the phytoplankton bloom state or community structure (Ribalet et al., 2014; Russo et al., 2020; Ulloa et al., 2025). A more complete time series of dissolved organics in important aquaculture regions over multiple years would help constrain the environmental factors and provide information to mitigate these ‘oxylipin events’ in the future.

### 4.5 Oxylipin Events, emerging problem for aquaculture

Oxylipin poisoning of larvae may be an emergent problem. Over the last decade, domoic acid production by diatoms, which causes amnesic shellfish poisoning, has emerged as a threat to marine ecosystems and aquaculture along the East Coast. Prior to the 2010s, domoic acid events were found on the West Coast (Sterling et al., 2022). Si and N stress, increased temperature, exposure to oxidative stress, and exposure to UV have been shown to increase oxylipin production in marine cultures and mesocosms (Collins et al., 2016, 2018; Cózar et al., 2018; Ramsden et al., 2019; Ribalet et al., 2009; Ulloa et al., 2025). Therefore, we would expect that climate change, specifically the increased temperature of the global ocean and increased nutrient stress due to stratification, deoxygenation, and denitrification, would promote oxylipin production and lead to more events. Our depleted ozone is a “natural” analog to the high levels of UV used to sterilize incoming hatchery water. Oxylipins may suppress food webs in regions exposed to excess UV or other oxidizing agents. For example, Collins et al. (2016) observed an increase in oxidized lipids in the ocean surface during polar day in the Antarctic. Additionally, if oxylipin events are an emerging issue, shifts in the phytoplankton community or benthic macroalgae could be a cause.

Other hatcheries along the East Coast of the US experience periodic unexplained crashes in larval and seed production (Gray et al., 2022) and a NOAA grant is currently wrapping up that documented two years of larval issues at MSF and two other hatcheries in Virginia and New York (NOAA Award NA22NMF4270132; (Glover, 2024). Therefore, it is important for the aquaculture industry to understand the cause of lower larval yields at Mook Sea Farm, the link with UV sterilization, and develop mitigation strategies.

### 5.1 Conclusion

The UV treatment of natural waters from the Damariscotta River estuary led to an increase in the relative concentrations of several oxidized lipids and oxylipins. Exposing oyster embryos to a commercially available oxylipin precursor resulted in the same digestive syndrome observed in larvae grown in UV-treated seawater. This is another example of how oxylipins decrease the developmental success of metazoans. This study employed a chemoinformatic pipeline focused on marine lipids, and another focused on publicly available datasets. Using untargeted analysis, we observed that UV treatment also increased the concentration of several pollutants and natural products that were not originally the focus of our study.

## Supporting information

Supplemental Report 1

Supplemental Report 2

## Acknowledgements

Mook Sea Farm (MSF) received funding to investigate larval digestive syndrome in 2020 through the Maine Sea Grant Program Development Funds, Bigelow Laboratory Center for Venture Research on Seafood Solutions, and Aquaculture Research Institute Rapid Response Funds. For their invaluable help during 2020 and 2021 troubleshooting, MSF thanks the entire MSF staff, Heather Leslie, Damian Brady, Paul Rawson, Larry Mayer, Tim Miller, Debbie Bouchard, Gayle Zydlewski, Nichole Price, Pete Countway, Nicole Poulton, Laura Lubelczyk, Chris Aeppli, Steve Archer, Marta Gomez-Chiarri, Evelyn Takyi, Gary Wikfors, Dave Veilleux, and Hal Schreier. Funding support for BRE, JS, and MW from NOAA Award NA22NMF4270132.

