## Supplemental Information for "Oxylipins and Other Natural Products Produced by UV-sterilization Impact Commercial Oyster Larvae Production in a New England Estuary"

Supplementary Methods
Supplemental Figures
Supplementary Tables- Please see the Excel file.

Supplementary Report 1- MetaboAnalyst Report for the LOBSTAHS Annotated Lipidome
Supplementary Report 2- MetaboAnalyst Report for Oxylipidome

**Supplemental Methods**

1. Standard Larval and Seed Production at Mook Sea Farm

The Mook Sea Farm hatchery pumps water from the Damariscotta River Estuary, with an annual salinity range of ~26-31 ppt, averaging around 29 ppt. Under normal circumstances, the hatcheries spawn broodstock oysters every two weeks from early January through early June, producing up to 120 million seeds annually. The hatchery obtains selectively bred diploid (2n) and tetraploid (4n) broodstock oysters from the Rutgers Haskin Shellfish Research Laboratory and the Aquaculture Genetics & Breeding Technology Center (ABC) at the Virginia Institute of Marine Science throughout the hatchery season, conditioning broodstock in quarantined tanks on-site at MSF. Broodstock and larval oysters were fed microalgae produced in-house: *Pavlova* sp., *Isochrysis galbana*, and *Chaetoceros* spp. produced using traditional phototrophic methodology, and *Tetraselmis* sp. (MC:2 strain) produced using proprietary heterotrophic methodology.

Since the late 1990s, seawater coming into the hatchery has been treated using serial (5 µm followed by 1 µm) polyester filter bags with welded seams into a head tank, followed by an in-line 1 µm polyester filter bag, followed by an Aqua Ultraviolet Classic 80 W UV sterilization unit, with a water flow rate that gives a UV dosage of ~52 $\mu W/s/cm^2$. Throughout the season, MSF produces multiple genetic crosses of oysters, resulting in diploid (2n) and triploid (3n) larvae, as seed are sold to farms along the East Coast, with different farms preferring diploid or triploid seed and different geographic regions needing different disease-resistant crosses. Broodstock spawns under quarantined conditions following standard heat shock techniques (Helm and Bourne 2004).

Larvae are raised in seawater heated to 24 °C and fed daily starting on day 1 once they have developed a mouth, with algal species appropriate for larval size. Larvae were observed and photographed daily under a compound microscope (OMAX 40X-1600X M837FLR Trinocular compound microscope equipped with an OMAX A35140U 14MP camera) to monitor development. Water is changed and larvae are 'graded' (sorted by size) through a stack of increasingly smaller mesh-size screens every other day. Signs of competency for setting typically appear by day 14 and include the development of a single eyespot, visible through the shell, a foot (not always visible), and the behavior of 'mucus-rafting,' as the larvae secrete a mucus that draws them together. Once larvae show signs of competency in setting (metamorphosis to juveniles), they are moved to a separate setting system to promote metamorphosis. The seed oysters were removed from the setting system and maintained in various upwelling systems. Seed are maintained in the hatchery until ambient conditions (seawater temperature and chlorophyll) at receiving farms are sufficient to support seed growth, at which point, seed are shipped from the hatchery to receiving farms, typically starting in late April or early May, with shipments made throughout the summer.

As described in the introduction, during the hatchery season of 2020, larvae reared following standard hatchery production methods presented with a digestive syndrome noticed by day 2, resulting in the inability to grow. Daily larval observations and micrographs were collected as described above. Experiments to determine the cause of digestive syndrome were carried out in smaller static tanks (15 L to 100 L), including testing every aspect of standard hatchery production techniques. After it was determined that eliminating the use of UV sterilization remedied the symptoms of digestive syndrome, production resumed as described above, with the elimination of the UV irradiation, although water continued to pass through the UV sterilizer with the bulbs turned off.

2. Histology of larval oysters

Oyster larvae were prepared at the hatchery for histological evaluation by fixing 10,000-100,000 larvae in ~10 mL of Davidson's fixative and shipping overnight to the Roger Williams University Aquatic Diagnostic Laboratory. Larvae were embedded in agar and paraffin and step-sectioned.

2. Dissolved organic matter analysis using untargeted lipidomics

2.1 Extraction and UPLC-MS$^n$ analysis

At UC-Berkeley, the samples were thawed in a whirlpak bag before flushing with ~3 mL of LC-grade high-purity water. The dissolved organics were eluted from the SPE cartridges using ~2 mL of methanol. As an internal standard,10 µL of 50 ng/mL Equi-SPLASH was added to the glass centrifuge tube used as a collection vial for SPE elution.

Samples were analyzed using ultra-high-performance liquid chromatography (UPLC) on an Agilent Vanquish system paired with an Orbitrap ID-X Mass Spectrometer (Thermo Scientific, San Jose, CA) for high-resolution accurate mass mass spectrometry (HRAM-MS). A reverse phase method was used to separate the compounds by hydrophobicity on a Thermo Accucore C8 column (150 mm, 2.1 mm, 2.1 µm). The mobile phases were water for eluent A and 70:30 acetonitrile: isopropanol for eluent B. The additives acetic acid and ammonium acetate were added to both eluents A and B to help with adduct formation. The gradient was as follows: 55% eluent B isocratic hold for first minute followed by an increase to 99% eluent B over 17 min, and an isocratic hold for 10 min.

A data-dependent acquisition strategy was employed, whereby compounds greater than $5x10^4$ were fragmented to produce $ms^2$, which were reanalyzed in the orbitrap mass analyzer. Oxylipins and their precursor fatty acids were best analyzed in negative mode. Other data acquisition settings for the mass spectrometry include resolution = 120,000, scan range = 100-

1000 m/z, stepped collision energy = 25, 30, 35 %, spray voltage = 2500 V, vaporizer temperature = 350 C, and ion transfer tube temperature = 325 C.

2.2.2 Chemoinformatics Pipeline

Lipidomic features (distinct m/z and retention time) were putatively annotated as lipids based on their $ms^1$ and adduct hierarchy using the LOBSTAHs database (Collins et al., 2016). Peaks were manually verified, and $ms^2$ was checked for the loss of water and acetate, which are characteristic of oxylipins in El-MAVEN (Agrawal et al. 2019). Some peaks were further annotated to the functional and positional levels using a fragmentation library populated with *in silico* models of oxylipin fragmentation and $ms^2$ from reference standards run on a wide variety of mass spectrometers in MS-DIAL and GNPS (Tsugawa et al. 2020, Wang et al. 2016). Please see **figure S1** for our lipidomic workflow, and **table 3** for details of our data reduction through the annotation process.

2.2.3 Comparative Lipidomics

The peak intensity of each compound was normalized to the volume that was filtered and the recovery of the deuterated- lyso phosphatidylethanolamine (LysoPE) from the internal standard mix. Statistical analyses were performed using MetaboAnalyst 5.0, after mean normalization and log transformation of the data (Pang et al. 2022). We analyzed the untargeted lipidome, which included annotated compounds, as well as unannotated mass spectrometry features reported as a mass-to-charge ratio (m/z) and retention time (RT). We also analyzed the annotated lipidome (N= 56 compounds) and the oxylipins (N=19 compounds) as separate datasets.

The standard suite of statistics used to assess the change in the dissolved lipidome in response to UV exposure includes parametric testing, multivariate analysis, and machine learning approaches. Full reports for each data subset can be found in **Supplemental Reports 1-2**. The various types of statistical analyses agreed with one another, but some analyses teased apart trends more readily than others did. Here, we present hierarchical clustering of samples and features within the lipidome for the full untargeted lipidome, annotated lipidome, oxylipins, and free fatty acid precursors (i.e., the oxylipidome; Ward and Euclidean; **figure 6**). For the untargeted lipidome, we present the sPLS-DA analysis to demonstrate how well the treatments were separated and which features were the most important for distinguishing between UV and non-UV treatments (**figure 7(a)**). Random Forest analyses were used to demonstrate the diagnostic compounds with putative lipid annotations for the UV vs. non-UV treatments, as well as for 2020 vs. 2021 (**figure 8(a)**), whereas PLS-DA was used to demonstrate the difference between compounds annotated as oxylipins (**figure 9(a)**).

**Supplemental Figures**

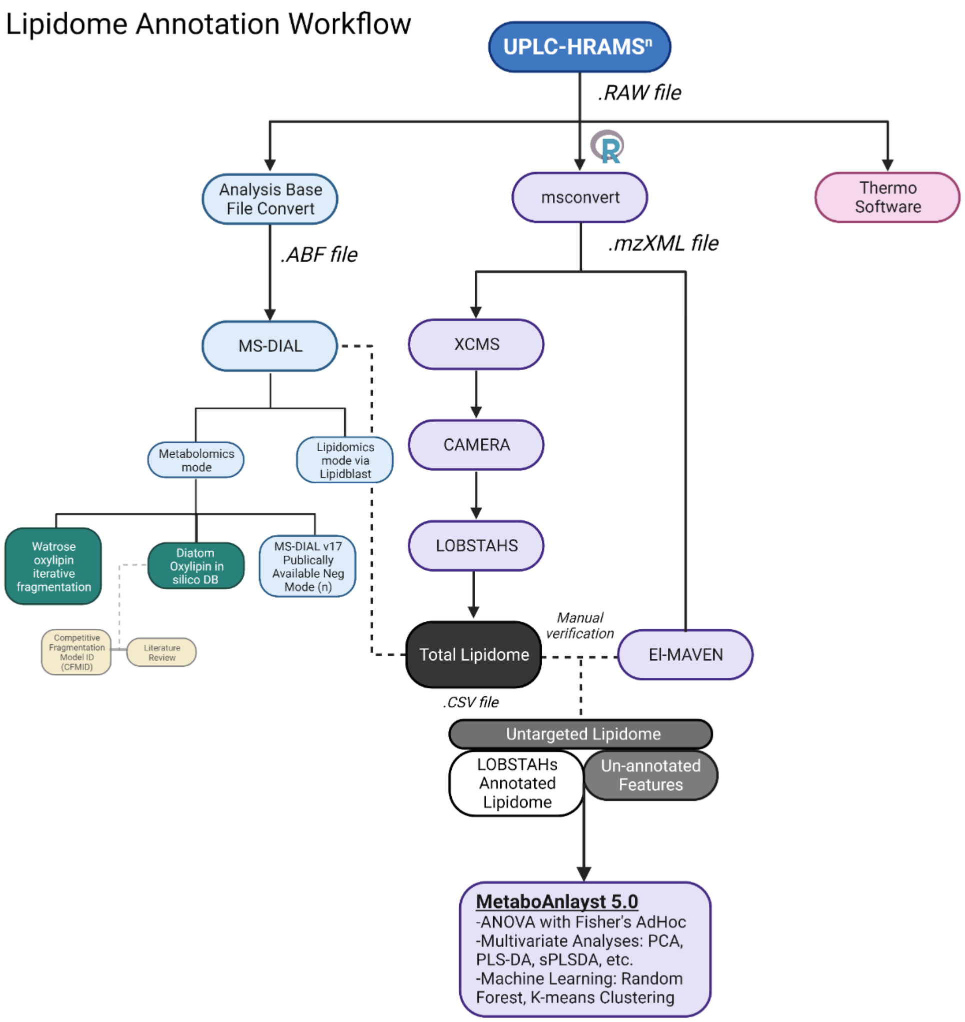


Supplemental Figure 1. Lipidomic workflow employed for the mass spectrometric analysis. Dissolved lipidome samples were analyzed on an Orbitrap ID-X instrument generating ultrahigh performance liquid chromatography- high resolution accurate mass spectrometry (UPLC-HRAMS$^n$) data files (i.e., .raw files) with data dependent secondary fragmentation (ms$^2$). These .raw files were analyzed based on their ms1 and adduct formation using LOBSTAHs ( ), an iterative in silico database containing oxylipins and fatty acids with 12-24 carbons, 0-6 double bond equivalents, and 0-4 oxygenations (purple pathway). Analysis in MS-DIAL (teal pathway) allowed us to compare our data to publicly available spectral libraries and boutique in silico databases presented in Watrose et al., (2018) and Edwards et al., (2024). The untargeted lipidome refers to the LOBSTAHS annotated lipidome plus the compounds pulled out with XCMS and CAMERA that did not match the database (i.e., un-annotated features). Statistical analyses were done in MetaboAnalyst for the untargered lipidome, the LOBSTAHS annotated lipidome, and the oxylipidome (the oxylipins only).

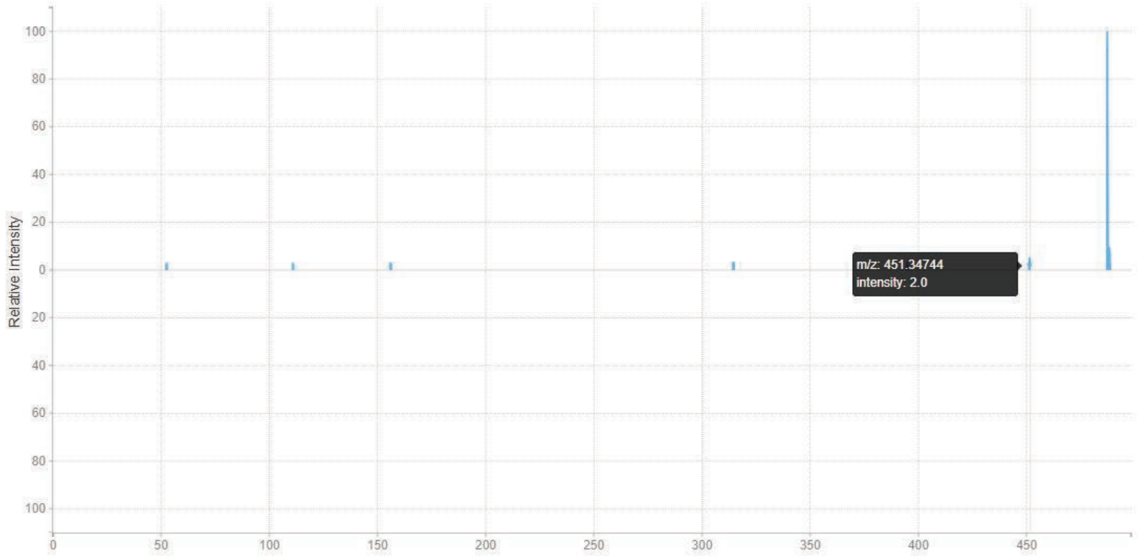


Supplemental Figure 2. Fragmentation for a compound originally annotated as DAG 26:6 +1O showing clear loss of a Chloride ion. The parent [M-H]- ion can be observed at 487.3060 Da in the ms2 spectra with the [M-Cl]- ion present at 451.34744 Da.

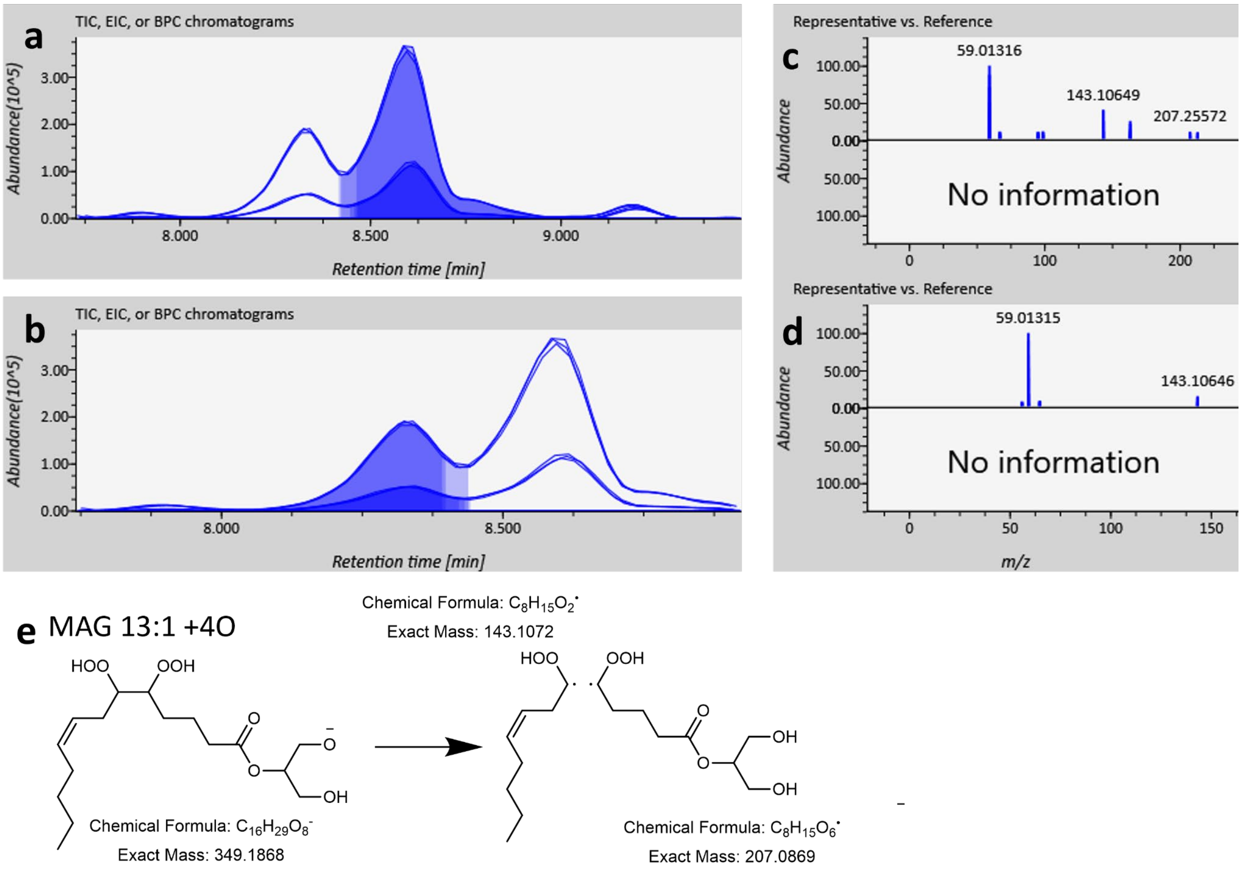


Supplemental Figure 3. Fragmentation of two MAG 13:1 +4O isomers at retention times of a) 8.9 and b) 8.4 with extracted ion chromatograms for the mass 349.1868 and ms2 fragmentation for each isomer separately (c-d). Proposed fragmentation for generating a 143.1072 fragment (m/z 143.10646 observed which is 5.17 ppm off; ms2 mass tolerance was set to 10 ppm) from a hypothetical MAG 13:1 +4O isomer, 5,6- dihydroperoxy trideca-8-enoic acid.

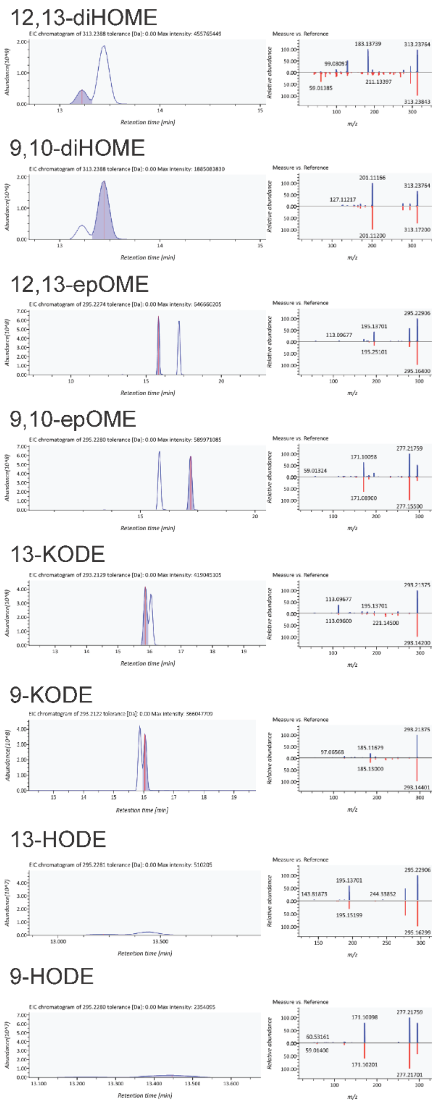


Supplemental Figure 4. Extracted ion chromatograms and ms2 fragmentation of Linoleic Acid Oxylipins MaxSpec® LC-MS Mixture from Caymen chemical. The authentic standard mixture contained 12,13 dihydroxy octadecaenoic acid (12,13-diHOME; NVO 18:1 +2O), 9,10-diHOME, 12,13-epoxy octadecaenoic acid (12,13-epOME; NVO 18:2 +1O), 9,10 epOME, 13-keto octadecadienoic acid (13-KODE; NVO 18:3 +1O), 9-KODE, 13- hydroxy octadecadienoic acid (13-HODE; NVO 18:2 +1O), and 9-HODE. The 13- isomers elutes before the 9- isomers. The dihydroxy oxylipins elute earliest in our reverse phase method. For regioisomers, epOME and HODE, the hydroxy acids elute before the epoxy acids. The ms2 fragmentation we observed is displayed on the top panel of the spectra with the reference standard from the boutique database built with CFM-ID.

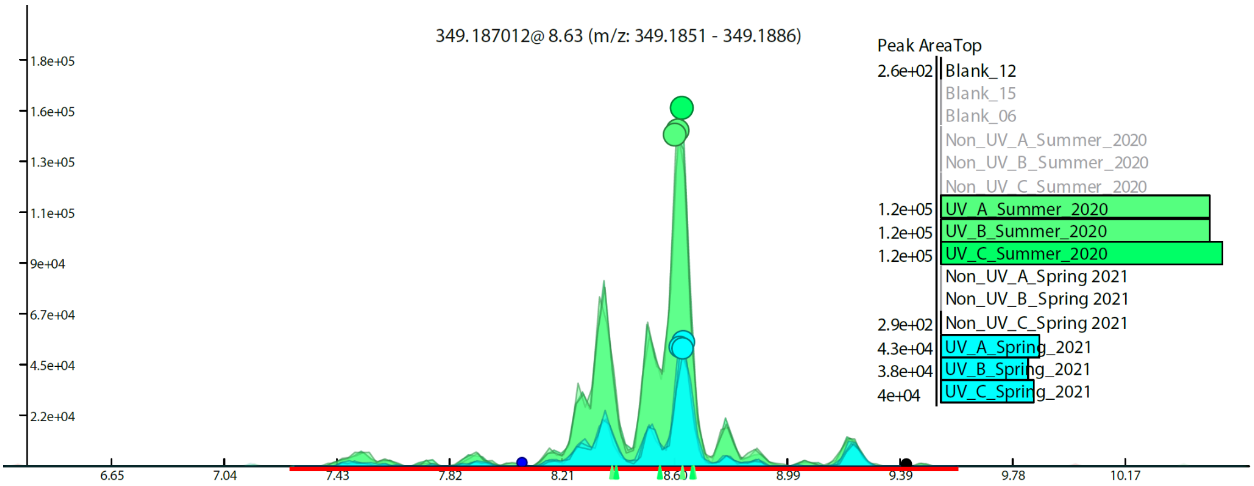


Supplemental Figure 5. Extracted ion chromatogram of the mass 349.187042 with the peak area in each sample integrated across all MAG 13:1 +4O isomers from ~7.43 to ~9.39 minutes in the bar chart. In both years, these isomers were more abundant in the UV treated samples than in the samples without UV treatment (i.e., filtration only). UV appeared to elicit a larger increase in MAG 13+4O isomers in Spring 2021 compared to Summer 2020, but it may be due to a seasonal difference.

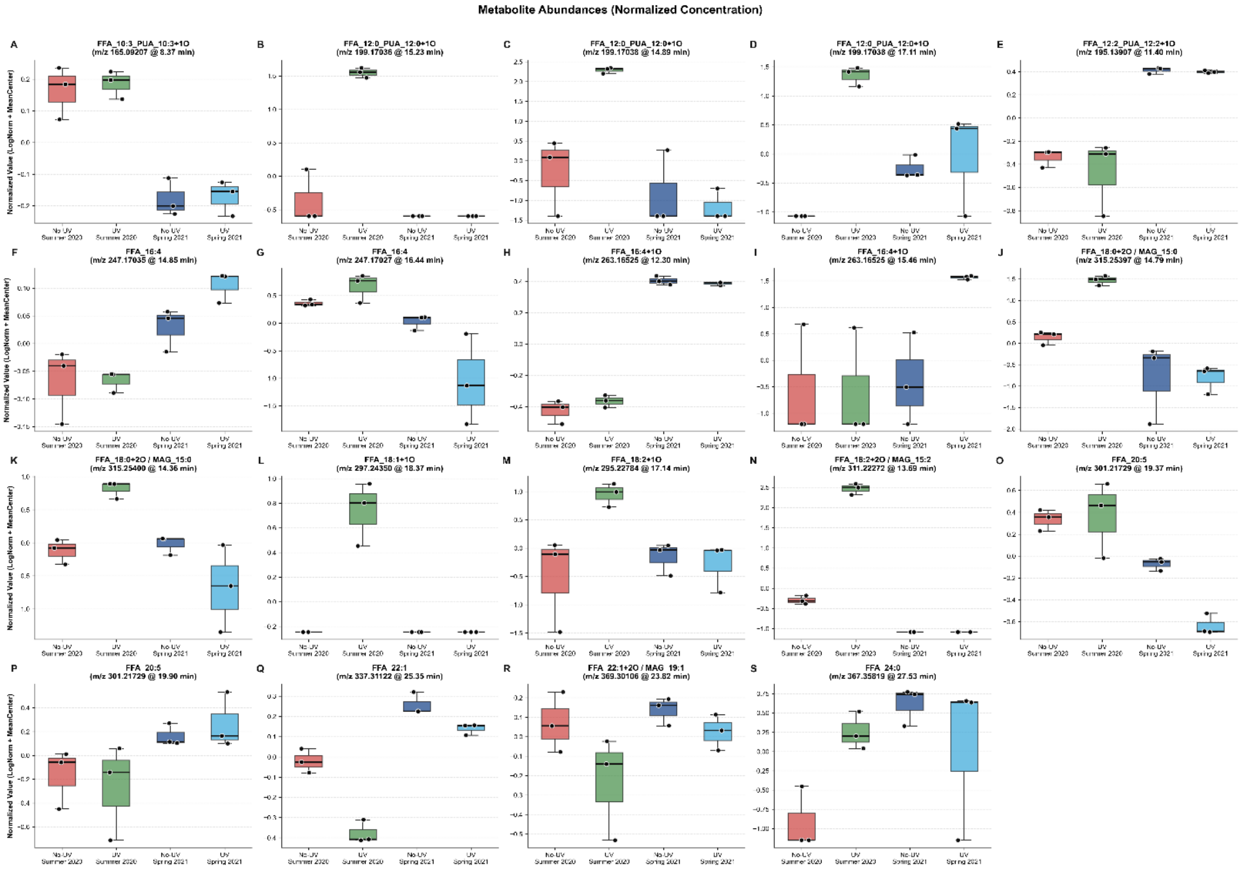


Supplemental Figure 6. Box-and-whisker plots of significant compounds within the oxylipidome. A-S) Average peak abundance (Original Conc.) and the average fold change from the mean (Normalized Conc.) for the 19 compounds that were significantly different between treatments. See posthoc test for significant relationships (Supplemental Table 4).
