## Supplemental Report 1 for "Oxylipins and Other Natural Products Produced by UV-sterilization Impact Commercial Oyster Larvae Production in a New England Estuary"

### Metabolomic Data Analysis with MetaboAnalyst 5.0

Name: guest7036401620298107543

January 23, 2023

#### 1 Data Processing and Normalization

##### 1.1 Reading and Processing the Raw Data

MetaboAnalyst accepts a variety of data types generated in metabolomic studies, including compound concentration data, binned NMR/MS spectra data, NMR/MS peak list data, as well as MS spectra (NetCDF, mzXML, mzDATA). Users need to specify the data types when uploading their data in order for MetaboAnalyst to select the correct algorithm to process them. Table 1 summarizes the result of the data processing steps.

###### 1.1.1 Reading Peak Intensity Table

The peak intensity table should be uploaded in comma separated values (.csv) format. Samples can be in rows or columns, with class labels immediately following the sample IDs.

Samples are in columns and features in rows. The uploaded file is in comma separated values (.csv) format. The uploaded data file contains 12 (samples) by 220 (peaks(mz/rt)) data matrix.

###### 1.1.2 Data Integrity Check

Before data analysis, a data integrity check is performed to make sure that all the necessary information has been collected. The class labels must be present and contain only two classes. If samples are paired, the class label must be from -n/2 to -1 for one group, and 1 to n/2 for the other group (n is the sample number and must be an even number). Class labels with same absolute value are assumed to be pairs. Compound concentration or peak intensity values should all be non-negative numbers. By default, all missing values, zeros and negative values will be replaced by the half of the minimum positive value found within the data (see next section)

###### 1.1.3 Missing value imputations

Too many zeroes or missing values will cause difficulties for downstream analysis. MetaboAnalyst offers several different methods for this purpose. The default method replaces all the missing and zero values with a small values (the half of the minimum positive values in the original data) assuming to be the detection limit. The assumption of this approach is that most missing values are caused by low abundance metabolites (i.e.below the detection limit). In addition, since zero values may cause problem for data normalization (i.e. log), they are also replaced with this small value. User can also specify other methods, such as replace by mean/median, or use K-Nearest Neighbours (KNN), Probabilistic PCA (PPCA), Bayesian PCA (BPCA) method, Singular Value Decomposition (SVD) method to impute the missing values [1]. Please choose the one that is the most appropriate for your data.

[1] Stacklies W, Redestig H, Scholz M, Walther D, Selbig J. *pcaMethods: a bioconductor package, providing PCA methods for incomplete data.*, Bioinformatics 2007 23(9):1164-1167

Zero or missing values were replaced by 1/5 of the min positive value for each variable.

##### 1.1.4 Data Filtering

The purpose of the data filtering is to identify and remove variables that are unlikely to be of use when modeling the data. No phenotype information are used in the filtering process, so the result can be used with any downstream analysis. This step can usually improves the results. Data filter is strongly recommended for datasets with large number of variables (> 250) datasets contain much noise (i.e.chemometrics data). Filtering can usually improve your results[2].

*For data with number of variables < 250, this step will reduce 5% of variables; For variable number between 250 and 500, 10% of variables will be removed; For variable number bwteen 500 and 1000, 25% of variables will be removed; And 40% of variabled will be removed for data with over 1000 variables. The None option is only for less than 5000 features. Over that, if you choose None, the IQR filter will still be applied. In addition, the maximum allowed number of variables is* ***10000***

No filtering was applied

Table 1: Summary of data processing results

| | Features (positive) | Missing/Zero | Features (processed) |
|---|---|---|---|
| Non_UV_A_Summer_2020 | 185 | 35 | 220 |
| Non_UV_B_Summer_2020 | 182 | 38 | 220 |
| Non_UV_C_Summer_2020 | 185 | 35 | 220 |
| Recirc_A_Summer_2020 | 211 | 9 | 220 |
| Recirc_B_Summer_2020 | 210 | 10 | 220 |
| Recirc_C_Summer_2020 | 211 | 9 | 220 |
| Non_UV_A_Spring_2021 | 190 | 30 | 220 |
| Non_UV_B_Spring_2021 | 188 | 32 | 220 |
| Non_UV_C_Spring_2021 | 187 | 33 | 220 |
| Recirc_A_Spring_2021 | 188 | 32 | 220 |
| Recirc_B_Spring_2021 | 197 | 23 | 220 |
| Recirc_C_Spring_2021 | 196 | 24 | 220 |

[2] Hackstadt AJ, Hess AM.*Filtering for increased power for microarray data analysis*, BMC Bioinformatics. 2009; 10: 11.

#### 1.2 Data Normalization

The data is stored as a table with one sample per row and one variable (bin/peak/metabolite) per column. The normalization procedures implemented below are grouped into four categories. Sample specific normalization allows users to manually adjust concentrations based on biological inputs (i.e. volume, mass); row-wise normalization allows general-purpose adjustment for differences among samples; data transformation and scaling are two different approaches to make features more comparable. You can use one or combine both to achieve better results.

The normalization consists of the following options:

1. Row-wise procedures:
   - Sample specific normalization (i.e. normalize by dry weight, volume)
   - Normalization by the sum
   - Normalization by the sample median
   - Normalization by a reference sample (probabilistic quotient normalization)[3]
   - Normalization by a pooled or average sample from a particular group
   - Normalization by a reference feature (i.e. creatinine, internal control)
   - Quantile normalization
2. Data transformation :
   - Log transformation (base 10)
   - Square root transformation
   - Cube root transformation
3. Data scaling:
   - Mean centering (mean-centered only)
   - Auto scaling (mean-centered and divided by standard deviation of each variable)
   - Pareto scaling (mean-centered and divided by the square root of standard deviation of each variable)
   - Range scaling (mean-centered and divided by the value range of each variable)

Figure 1 shows the effects before and after normalization.

[3]Dieterle F, Ross A, Schlotterbeck G, Senn H. *Probabilistic quotient normalization as robust method to account for dilution of complex biological mixtures. Application in 1H NMR metabonomics*, 2006, Anal Chem 78 (13);4281 - 4290

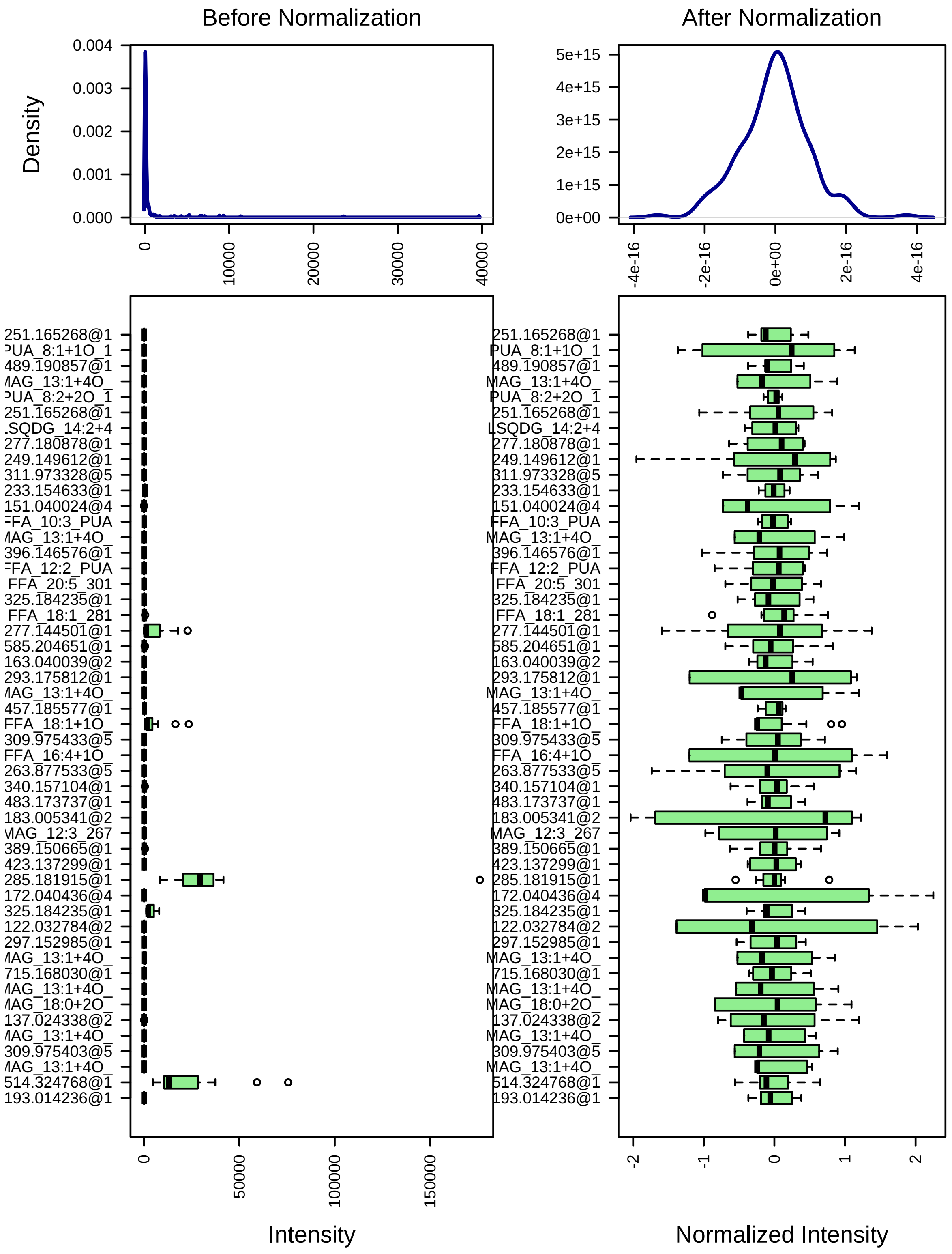


Figure 1: Box plots and kernel density plots before and after normalization. The boxplots show at most 50 features due to space limit. The density plots are based on all samples. Selected methods : Row-wise normalization: N/A; Data transformation: Log10 Normalization; Data scaling: Mean Centering.

#### 2 Statistical and Machine Learning Data Analysis

MetaboAnalyst offers a variety of methods commonly used in metabolomic data analyses. They include:

1. Univariate analysis methods:
   - Fold Change Analysis
   - T-tests
   - Volcano Plot
   - One-way ANOVA and post-hoc analysis
   - Correlation analysis
2. Multivariate analysis methods:
   - Principal Component Analysis (PCA)
   - Partial Least Squares - Discriminant Analysis (PLS-DA)
3. Robust Feature Selection Methods in microarray studies
   - Significance Analysis of Microarray (SAM)
   - Empirical Bayesian Analysis of Microarray (EBAM)
4. Clustering Analysis
   - Hierarchical Clustering
     - Dendrogram
     - Heatmap
   - Partitional Clustering
     - K-means Clustering
     - Self-Organizing Map (SOM)
5. Supervised Classification and Feature Selection methods
   - Random Forest
   - Support Vector Machine (SVM)

`Please note: some advanced methods are available only for two-group sample analyais.`

#### 2.1 One-way ANOVA

Univariate analysis methods are the most common methods used for exploratory data analysis. For multi-group analysis, MetaboAnalyst provides one-way Analysis of Variance (ANOVA). As ANOVA only tells whether the overall comparison is significant or not, it is usually followed by post-hoc analyses in order to identify which two levels are different. MetaboAnalyst provides two most commonly used methods for this purpose - Fisher's least significant difference method (Fisher's LSD) and Tukey's Honestly Significant Difference (Tukey's HSD). The univariate analyses provide a preliminary overview about features that are potentially significant in discriminating the conditions under study.

Figure 2 shows the important features identified by ANOVA analysis. Table 2 shows the details of these features. The `post-hoc Sig. Comparison` column shows the comparisons between different levels that are significant given the p value threshold.

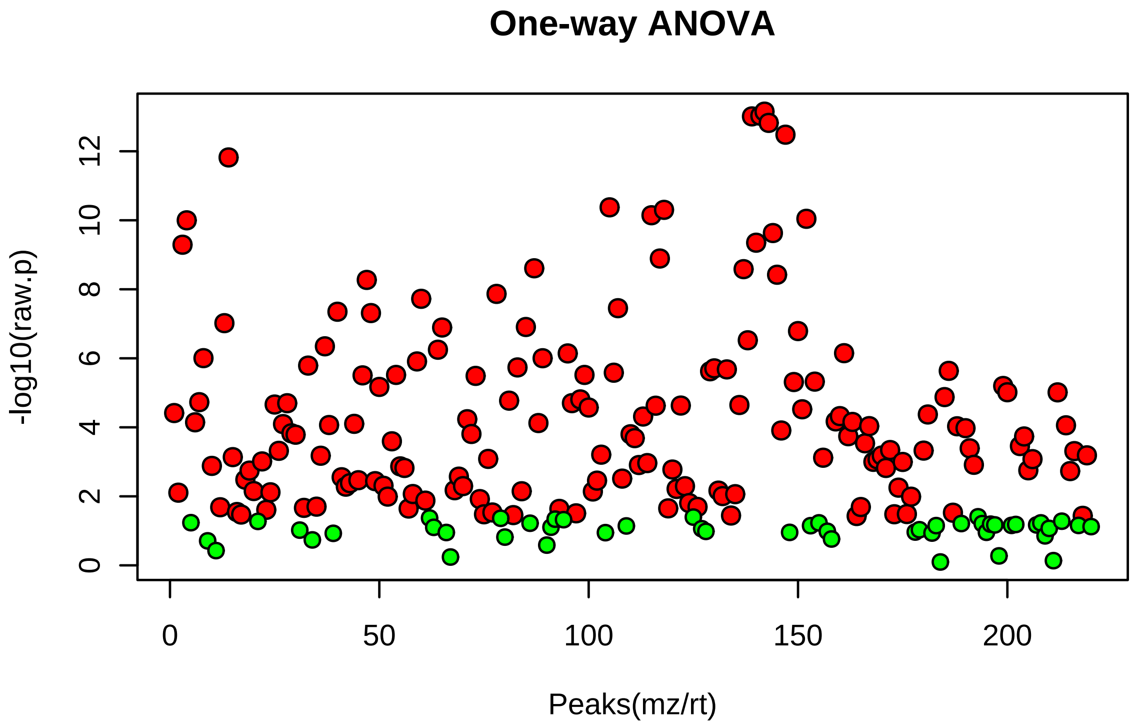


Figure 2: Important features selected by ANOVA plot with p value threshold 0.05.

Table 2: Top 50 features identified by One-way ANOVA and post-hoc analysis

| | Peaks(mz/rt) | f.value | p.value | -log10(p) | FDR | Fisher's LSD |
|---|---|---|---|---|---|---|
| 1 | MAG_13:1+4O_349.186859@8.604212 | 6466.000 | 7.1064e-14 | 13.1480 | 7.1574e-12 | b_UV_Summer_2020 - a_No |
| 2 | MAG_13:1+4O_349.186859@8.494180 | 6039.100 | 9.3375e-14 | 13.0300 | 7.1574e-12 | b_UV_Summer_2020 - a_No |
| 3 | MAG_13:1+4O_349.186798@8.338817 | 5972.600 | 9.7601e-14 | 13.0110 | 7.1574e-12 | b_UV_Summer_2020 - a_No |
| 4 | MAG_13:1+4O_349.186859@8.759778 | 5358.200 | 1.5064e-13 | 12.8220 | 8.2851e-12 | b_UV_Summer_2020 - a_No |
| 5 | MAG_13:1+4O_349.187012@9.212515 | 4400.800 | 3.3091e-13 | 12.4800 | 1.4560e-11 | b_UV_Summer_2020 - a_No |
| 6 | 144.045395@3.384538 | 3010.300 | 1.5096e-12 | 11.8210 | 5.5351e-11 | b_UV_Summer_2020 - a_No |
| 7 | 309.975403@4.722495 | 1307.700 | 4.2173e-11 | 10.3750 | 1.3254e-09 | a_No_UV_Summer_2020 - b |
| 8 | 311.973389@4.722494 | 1253.700 | 4.9898e-11 | 10.3020 | 1.3722e-09 | a_No_UV_Summer_2020 - b |
| 9 | FFA_18:2+2O_MAG_15:2_311.222717@13.685581 | 1145.600 | 7.1523e-11 | 10.1460 | 1.7483e-09 | b_UV_Summer_2020 - a_No |
| 10 | 359.147766@13.452975 | 1079.600 | 9.0623e-11 | 10.0430 | 1.9937e-09 | b_UV_Summer_2020 - a_No |
| 11 | 135.045059@2.652001 | 1052.700 | 1.0022e-10 | 9.9990 | 2.0045e-09 | b_UV_Summer_2020 - a_No |
| 12 | MAG_13:1+4O_349.186890@7.490250 | 850.590 | 2.3449e-10 | 9.6299 | 4.2989e-09 | b_UV_Summer_2020 - a_No |
| 13 | MAG_13:1+4O_349.186859@8.261336 | 723.560 | 4.4674e-10 | 9.3499 | 7.5601e-09 | b_UV_Summer_2020 - a_No |
| 14 | 135.045059@2.248155 | 699.540 | 5.1103e-10 | 9.2916 | 8.0305e-09 | b_UV_Summer_2020 - a_No |
| 15 | 311.973328@5.058717 | 555.130 | 1.2831e-09 | 8.8918 | 1.8818e-08 | a_No_UV_Summer_2020 - b |
| 16 | 277.144501@13.447971 | 471.160 | 2.4634e-09 | 8.6085 | 3.3778e-08 | b_UV_Summer_2020 - a_No |
| 17 | MAG_13:1+4O_349.186768@8.866138 | 464.350 | 2.6101e-09 | 8.5833 | 3.3778e-08 | b_UV_Summer_2020 - a_No |
| 18 | MAG_13:1+4O_349.186890@7.893439 | 422.920 | 3.7839e-09 | 8.4221 | 4.6248e-08 | b_UV_Summer_2020 - a_No |
| 19 | 179.034958@1.231828 | 387.610 | 5.3495e-09 | 8.2717 | 6.1942e-08 | b_UV_Summer_2020 - a_No |
| 20 | FFA_16:4+1O_263.165253@12.302199 | 306.330 | 1.3604e-08 | 7.8663 | 1.4964e-07 | c_No_UV_Spring_2021 - a_N |
| 21 | 221.154572@16.206219 | 281.980 | 1.8886e-08 | 7.7239 | 1.9785e-07 | b_UV_Summer_2020 - a_No |
| 22 | 309.975433@5.064293 | 240.710 | 3.5314e-08 | 7.4521 | 3.5314e-07 | a_No_UV_Summer_2020 - b |
| 23 | 172.040436@1.330356 | 226.750 | 4.4712e-08 | 7.3496 | 4.2768e-07 | b_UV_Summer_2020 - a_No |
| 24 | 179.107758@12.427021 | 221.710 | 4.8868e-08 | 7.3110 | 4.4796e-07 | b_UV_Summer_2020 - a_No |
| 25 | 144.045395@1.253267 | 186.880 | 9.5871e-08 | 7.0183 | 8.4367e-07 | b_UV_Summer_2020 - a_No |
| 26 | MAG_12:3_267.160156@8.557397 | 175.150 | 1.2374e-07 | 6.9075 | 1.0442e-06 | a_No_UV_Summer_2020 - c |
| 27 | MAG_10:3_239.128845@11.397508 | 173.590 | 1.2815e-07 | 6.8923 | 1.0442e-06 | c_No_UV_Spring_2021 - a_N |
| 28 | 353.826019@8.917586 | 163.410 | 1.6252e-07 | 6.7891 | 1.2770e-06 | a_No_UV_Summer_2020 - b |
| 29 | MAG_13:1+4O_349.186798@8.175578 | 139.760 | 3.0011e-07 | 6.5227 | 2.2767e-06 | b_UV_Summer_2020 - a_No |
| 30 | 165.019302@1.136820 | 126.070 | 4.4933e-07 | 6.3474 | 3.2951e-06 | b_UV_Summer_2020 - a_No |
| 31 | 233.154633@14.778790 | 118.930 | 5.6422e-07 | 6.2486 | 4.0041e-06 | b_UV_Summer_2020 - a_No |
| 32 | 379.156189@13.636354 | 112.110 | 7.1033e-07 | 6.1485 | 4.8118e-06 | a_No_UV_Summer_2020 - c |
| 33 | 293.175812@13.194001 | 111.650 | 7.2177e-07 | 6.1416 | 4.8118e-06 | c_No_UV_Spring_2021 - a_N |
| 34 | 138.019623@4.274860 | 102.950 | 9.9003e-07 | 6.0044 | 6.2542e-06 | b_UV_Summer_2020 - a_No |
| 35 | 278.147858@13.457795 | 102.820 | 9.9498e-07 | 6.0022 | 6.2542e-06 | b_UV_Summer_2020 - a_No |
| 36 | 219.175308@18.905254 | 97.310 | 1.2322e-06 | 5.9093 | 7.5304e-06 | a_No_UV_Summer_2020 - c |
| 37 | 163.040039@2.298887 | 90.595 | 1.6260e-06 | 5.7889 | 9.6683e-06 | b_UV_Summer_2020 - a_No |
| 38 | 265.147919@13.251337 | 87.761 | 1.8391e-06 | 5.7354 | 1.0647e-05 | a_No_UV_Summer_2020 - c |
| 39 | FFA_22:1_337.311218@25.352736 | 86.357 | 1.9574e-06 | 5.7083 | 1.1042e-05 | a_No_UV_Summer_2020 - b |
| 40 | 347.151031@13.249694 | 84.825 | 2.0977e-06 | 5.6782 | 1.1538e-05 | a_No_UV_Summer_2020 - c |
| 41 | LPG_10:1+2O_LPA_13:1+4O_429.154175@13.253326 | 82.726 | 2.3110e-06 | 5.6362 | 1.2401e-05 | a_No_UV_Summer_2020 - c |
| 42 | 330.810394@5.037743 | 82.127 | 2.3767e-06 | 5.6240 | 1.2450e-05 | a_No_UV_Summer_2020 - c |
| 43 | 309.975403@5.861267 | 80.127 | 2.6141e-06 | 5.5827 | 1.3374e-05 | a_No_UV_Summer_2020 - b |
| 44 | FFA_12:0_PUA_12:0+1O_199.170364@15.234773 | 77.113 | 3.0306e-06 | 5.5185 | 1.4825e-05 | b_UV_Summer_2020 - a_No |
| 45 | 297.152985@13.633081 | 77.101 | 3.0324e-06 | 5.5182 | 1.4825e-05 | a_No_UV_Summer_2020 - c |
| 46 | 176.035339@1.518626 | 76.505 | 3.1244e-06 | 5.5052 | 1.4943e-05 | b_UV_Summer_2020 - a_No |
| 47 | 251.165268@10.677092 | 75.829 | 3.2330e-06 | 5.4904 | 1.5133e-05 | c_No_UV_Spring_2021 - a_N |
| 48 | 362.122284@13.441428 | 68.645 | 4.7394e-06 | 5.3243 | 2.1722e-05 | b_UV_Summer_2020 - a_No |
| 49 | 350.125488@13.249694 | 68.153 | 4.8722e-06 | 5.3123 | 2.1875e-05 | a_No_UV_Summer_2020 - c |
| 50 | 505.140656@13.254817 | 63.627 | 6.3385e-06 | 5.1980 | 2.7889e-05 | a_No_UV_Summer_2020 - c |

#### 2.2 Sparse Partial Least Squares - Discriminant Analysis (sPLS-DA)

The sparse PLS-DA (sPLS-DA) algorithm can be used to effectively reduce the number of variables (metabolites) in high-dimensional metabolomics data to produce robust and easy-to-interpret models. Users can control the sparseness of the model by controlling the number of components in the model and the number of variables in each component. For more information, please refer to Cao et al. 2011 (PMC3133555).

Figure 3 shows the overview of scores plots; Figure 4 shows the 2-D scores plot between selected components; Figure 5 shows the loading plot of the top ranked features; Figure 6 shows the 3-D scores plot between selected components; Figure 7 shows the performance of the sPLS-DA model evaluated using cross-validations;

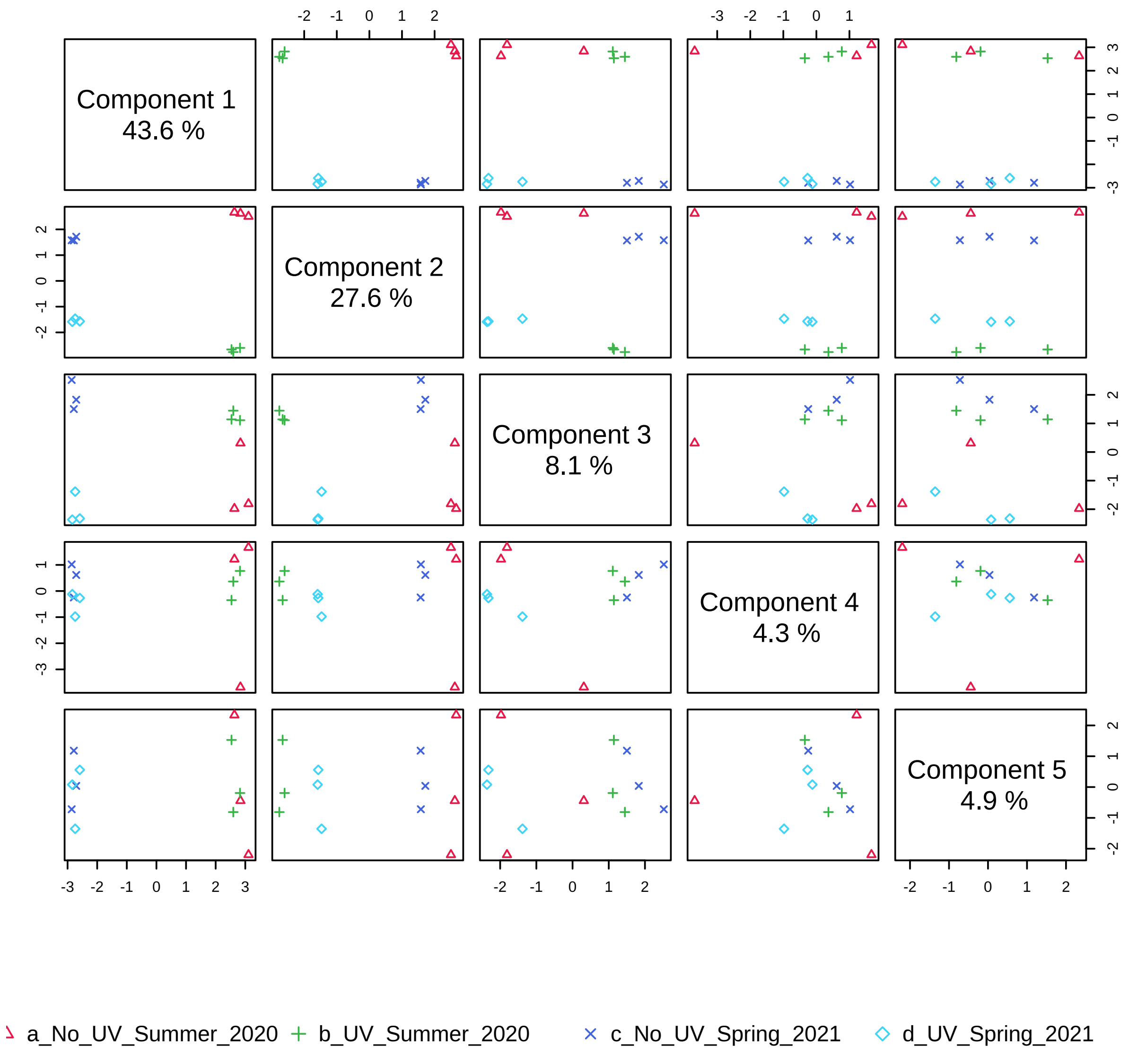


Figure 3: Pairwise scores plots between the selected components. The explained variance of each component is shown in the corresponding diagonal cell.

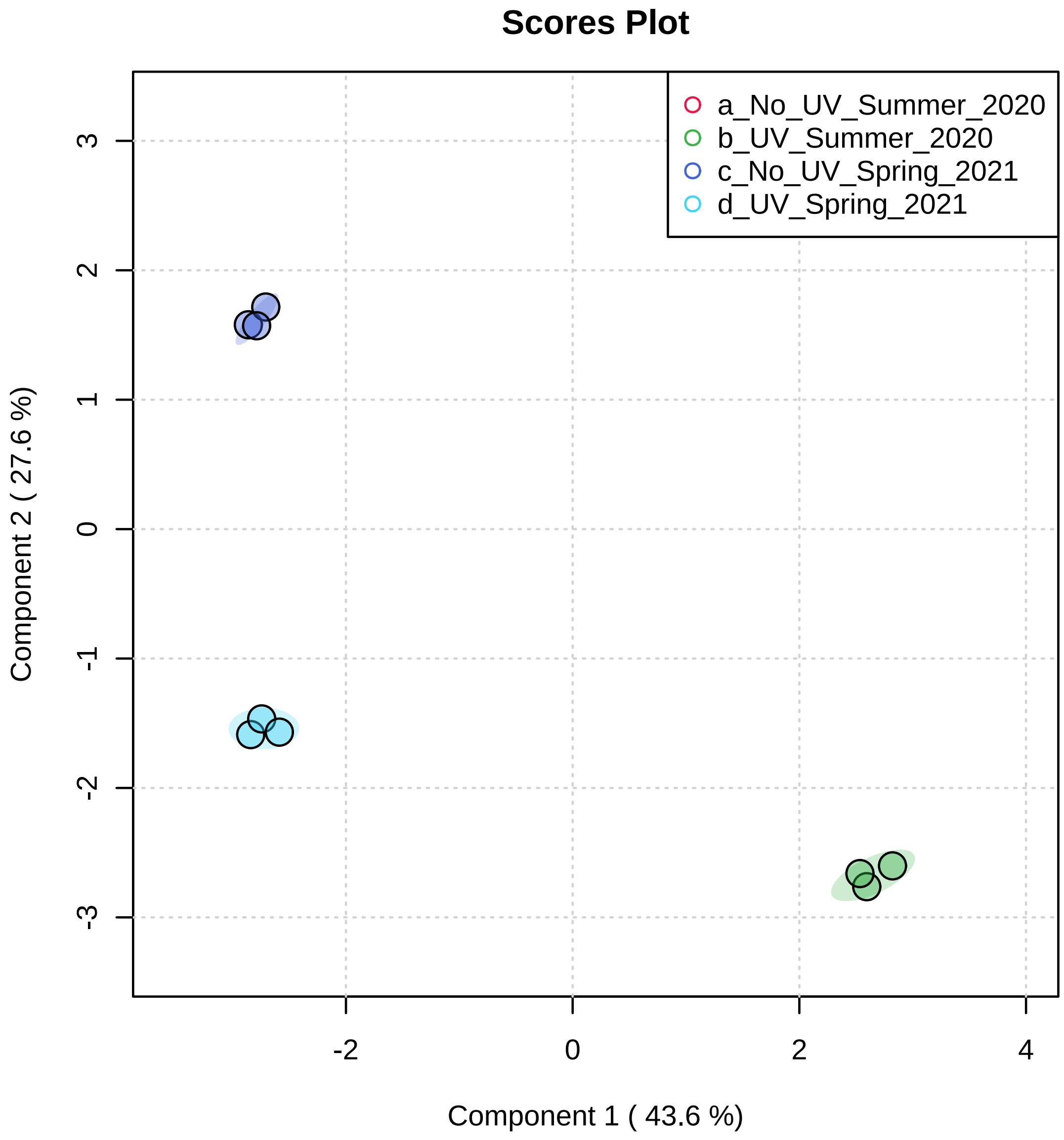


Figure 4: Scores plot between the selected PCs. The explained variances are shown in brackets.

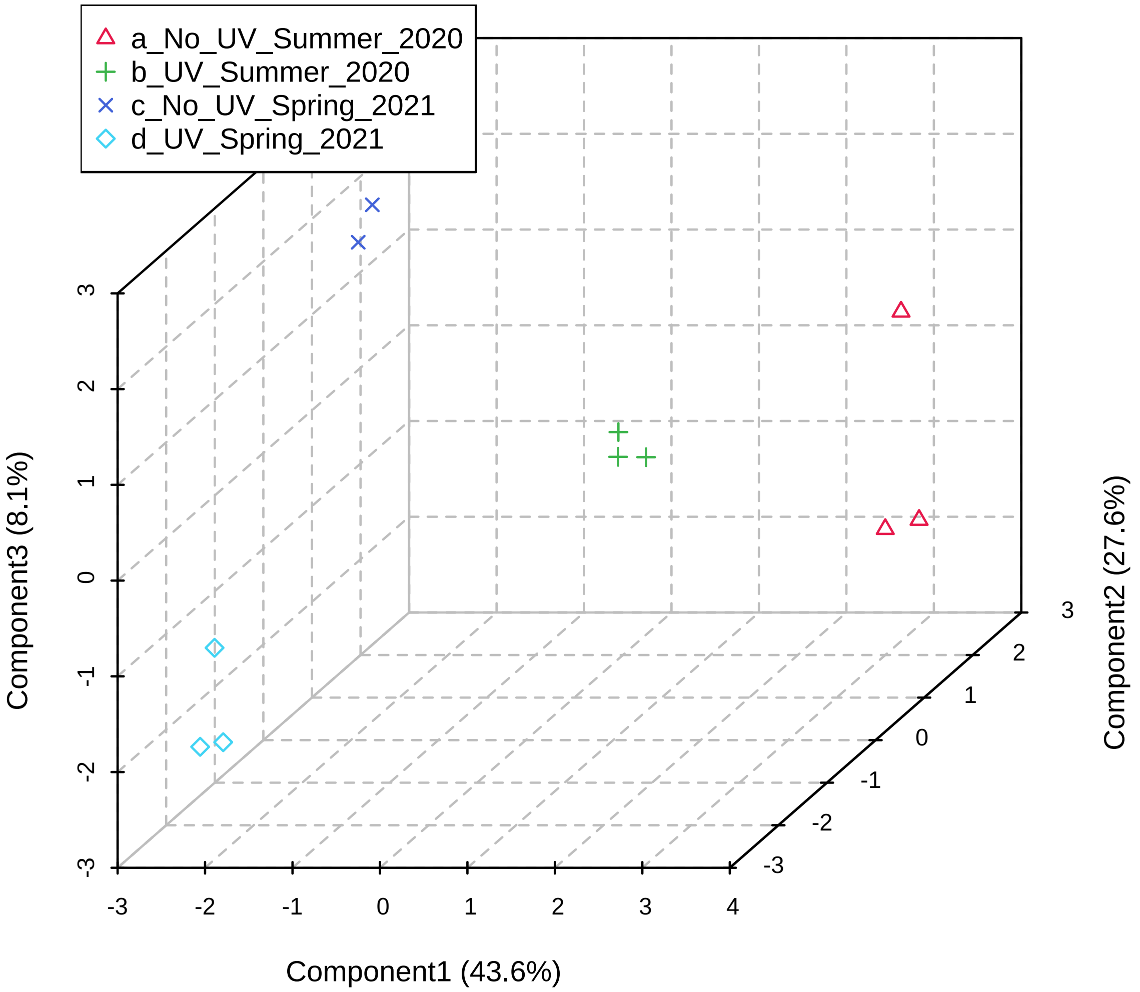


Figure 5: 3D scores plot between the selected PCs. The explained variances are shown in brackets.

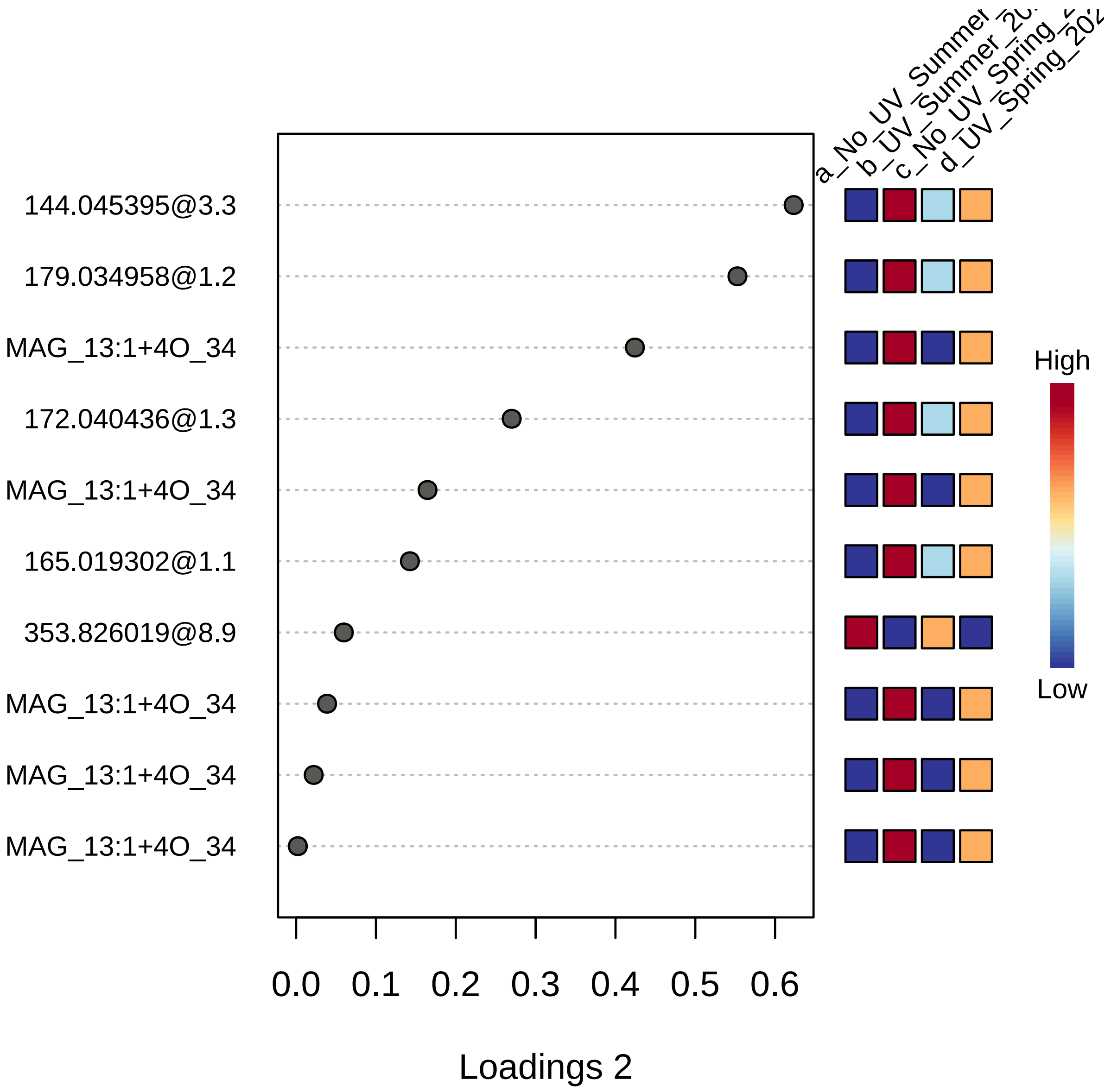


Figure 6: Plot showing the variables selected by the sPLS-DA model for a given component. The variables are ranked by the absolute values of their loadings.

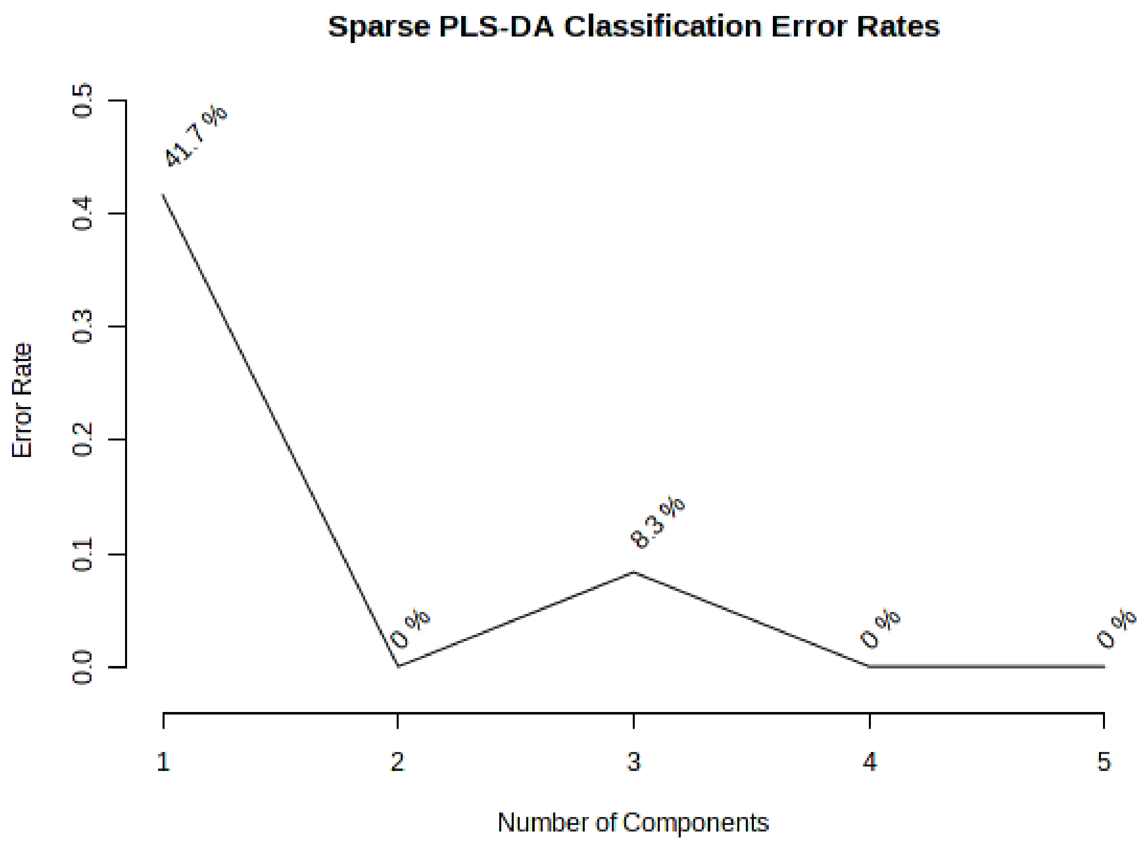


Figure 7: Plot of the performance of the sPLS-DA model evaluated using cross validations (CV) with increasing numbers of components created using the specified number of the variables. The error rate is on the y-axis and the number of components is on the x-axis.

##### 2.3 Hierarchical Clustering

In (agglomerative) hierarchical cluster analysis, each sample begins as a separate cluster and the algorithm proceeds to combine them until all samples belong to one cluster. Two parameters need to be considered when performing hierarchical clustering. The first one is similarity measure - Euclidean distance, Pearson's correlation, Spearman's rank correlation. The other parameter is clustering algorithms, including average linkage (clustering uses the centroids of the observations), complete linkage (clustering uses the farthest pair of observations between the two groups), single linkage (clustering uses the closest pair of observations) and Ward's linkage (clustering to minimize the sum of squares of any two clusters). Heatmap is often presented as a visual aid in addition to the dendrogram.

Hierachical clustering is performed with the `hclust` function in package `stat`. Figure 8 shows the clustering result in the form of a dendrogram. Figure 9 shows the clustering result in the form of a heatmap.

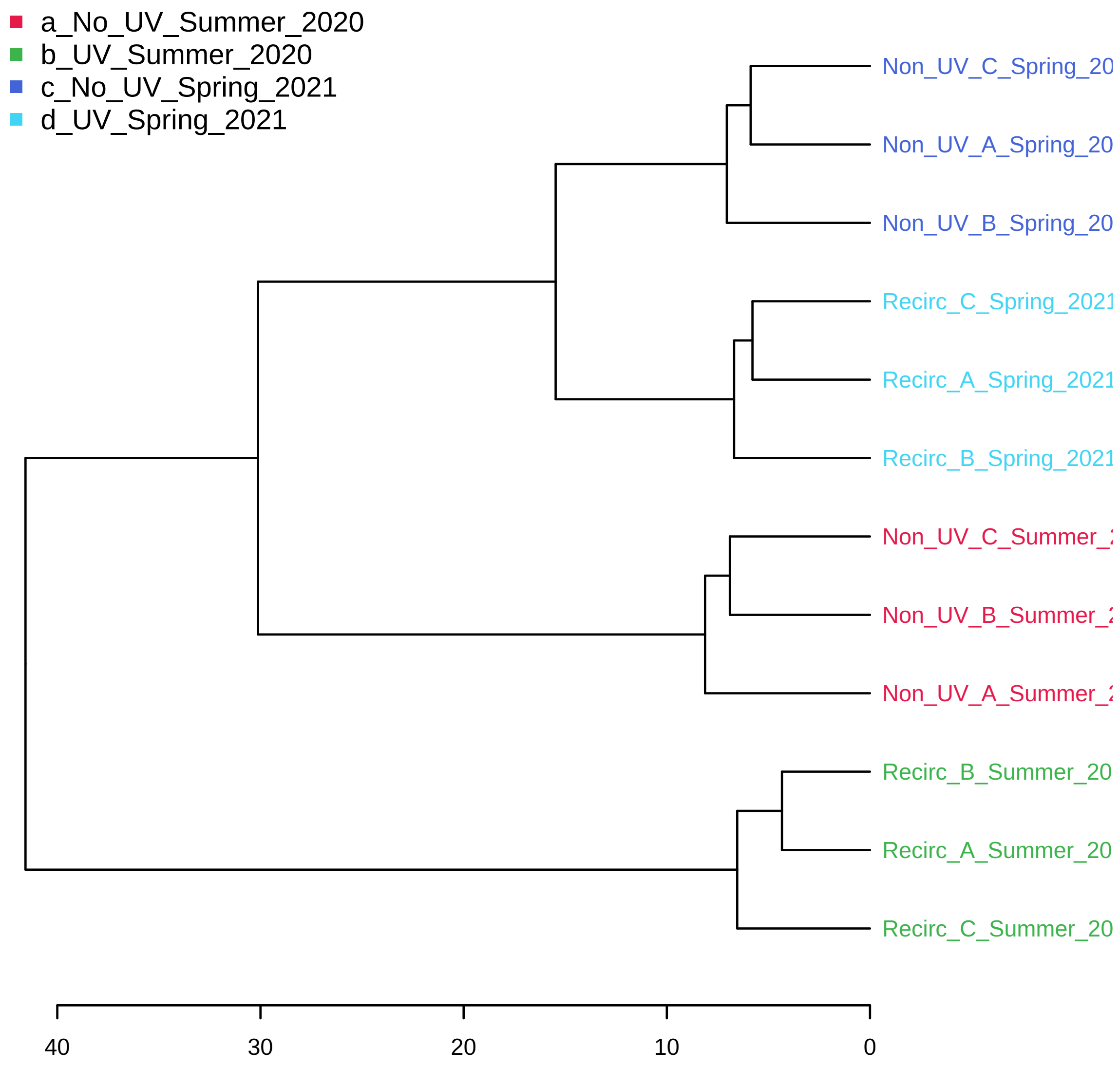


Figure 8: Clustering result shown as dendrogram (distance measure using `euclidean`, and clustering algorithm using `ward.D`).

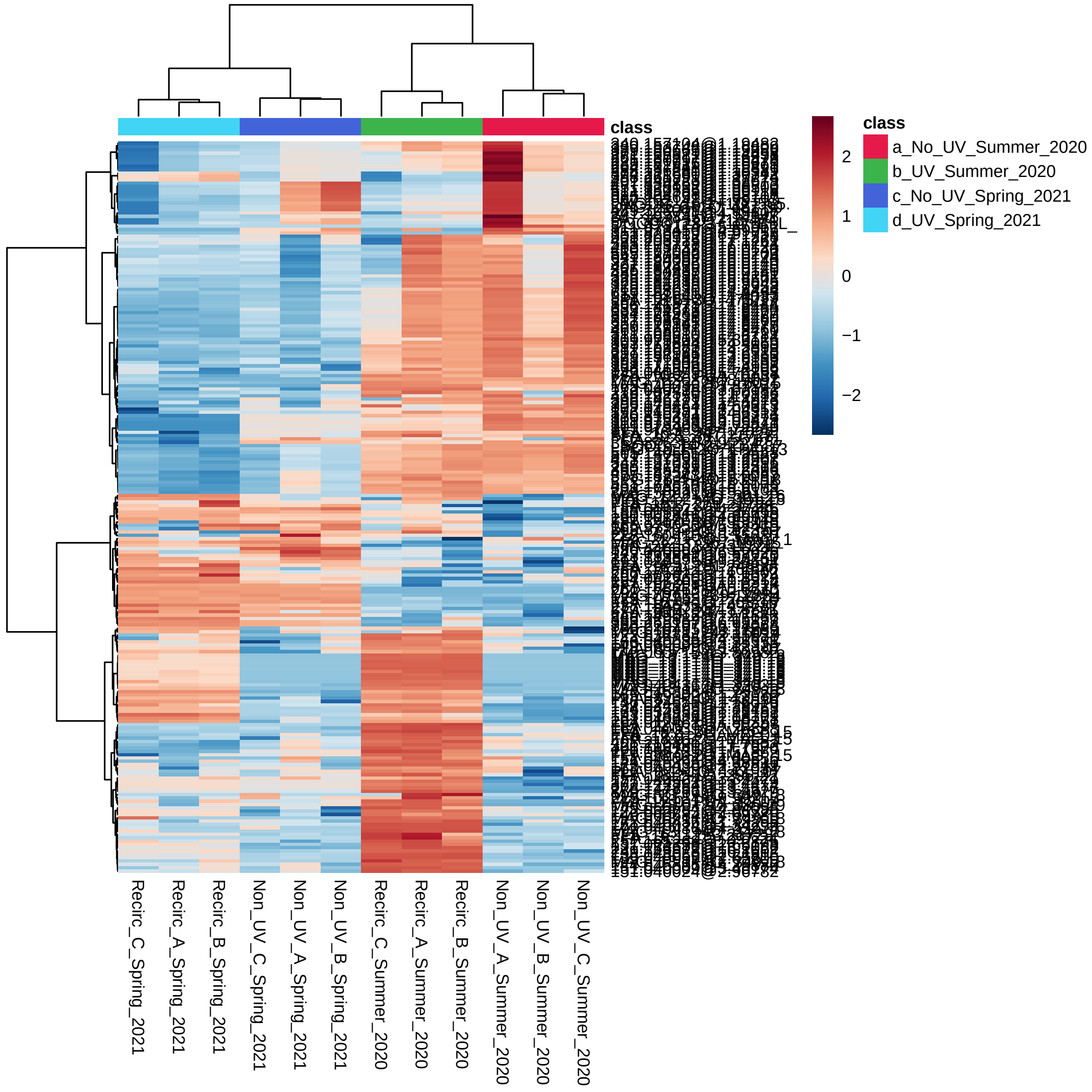


Figure 9: Clustering result shown as heatmap (distance measure using `euclidean`, and clustering algorithm using `ward.D`).

#### 2.4 K-means Clustering

K-means clustering is a nonhierarchical clustering technique. It begins by creating k random clusters (k is supplied by user). The program then calculates the mean of each cluster. If an observation is closer to the centroid of another cluster then the observation is made a member of that cluster. This process is repeated until none of the observations are reassigned to a different cluster.

K-means analysis is performed using the `kmeans` function in the package `stat`. Figure 10 shows clustering the results. Table 3 shows the members in each cluster from K-means analysis.

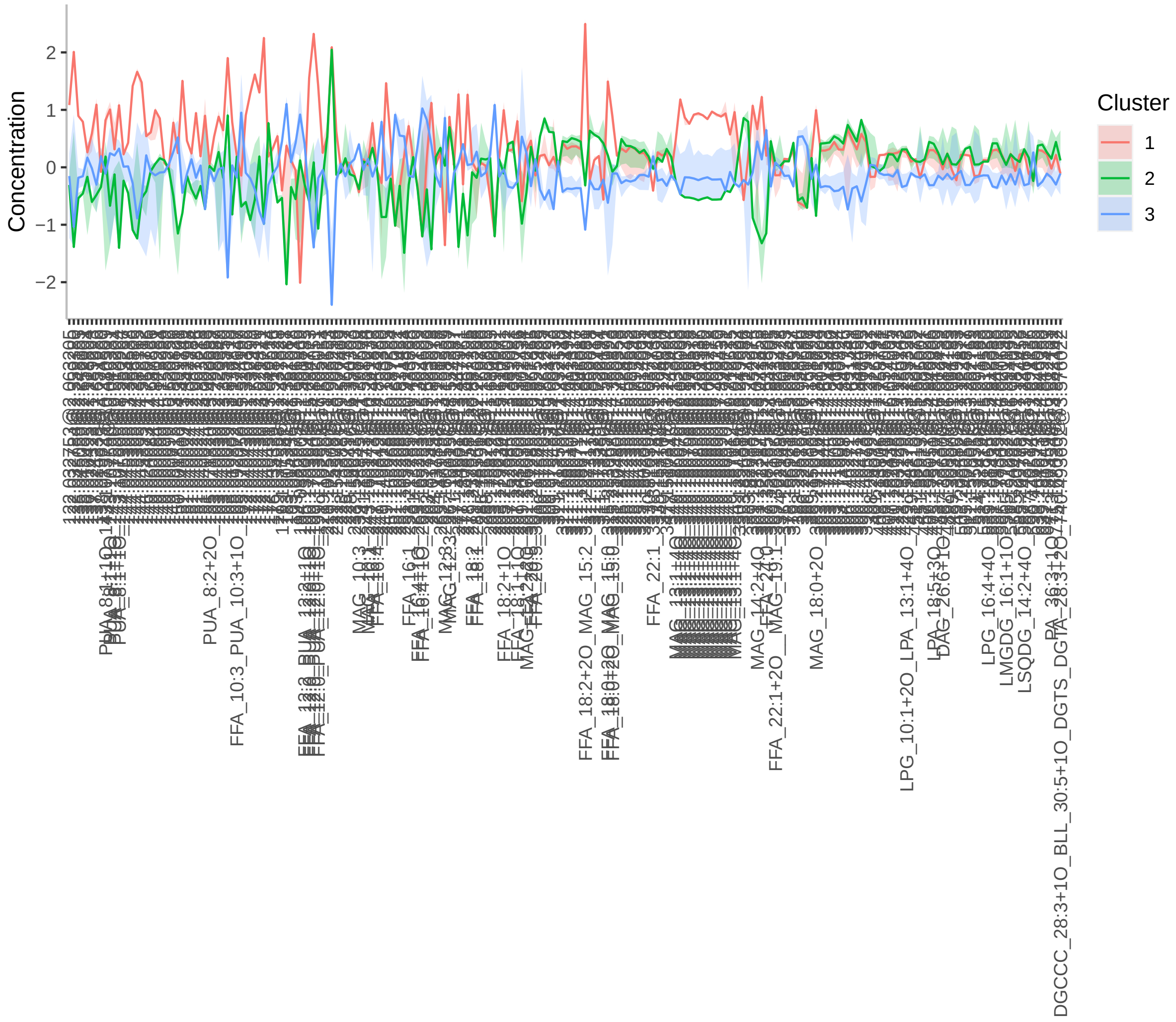


Figure 10: K-means cluster analysis. The x-axes are variable indices and y-axes are relative intensities. The blue lines represent median intensities of corresponding clusters

Table 3: Clustering result using K-means

| | Samples in each cluster |
|---|---|
| Cluster( 1 ) | Recirc_A_Summer_2020 Recirc_B_Summer_2020 Recirc_C_Summer_2020 |
| Cluster( 2 ) | Non_UV_A_Summer_2020 Non_UV_B_Summer_2020 Non_UV_C_Summer_2020 |
| Cluster( 3 ) | Non_UV_A_Spring_2021 Non_UV_B_Spring_2021 Non_UV_C_Spring_2021 Recirc_A_Spring_2021 Recirc_B_Spring_2021 Recirc_C_Spring_2021 |

#### 2.5 Random Forest (RF)

Random Forest is a supervised learning algorithm suitable for high dimensional data analysis. It uses an ensemble of classification trees, each of which is grown by random feature selection from a bootstrap sample at each branch. Class prediction is based on the majority vote of the ensemble. RF also provides other useful information such as OOB (out-of-bag) error, variable importance measure, and outlier measures. During tree construction, about one-third of the instances are left out of the bootstrap sample. This OOB data is then used as test sample to obtain an unbiased estimate of the classification error (OOB error). Variable importance is evaluated by measuring the increase of the OOB error when it is permuted. The outlier measures are based on the proximities during tree construction.

RF analysis is performed using the `randomForest` package[4]. Table 4 shows the confusion matrix of random forest. Figure 11 shows the cumulative error rates of random forest analysis for given parameters. Figure 12 shows the important features ranked by random forest. Figure 13 shows the outlier measures of all samples for the given parameters. The OOB error is 0

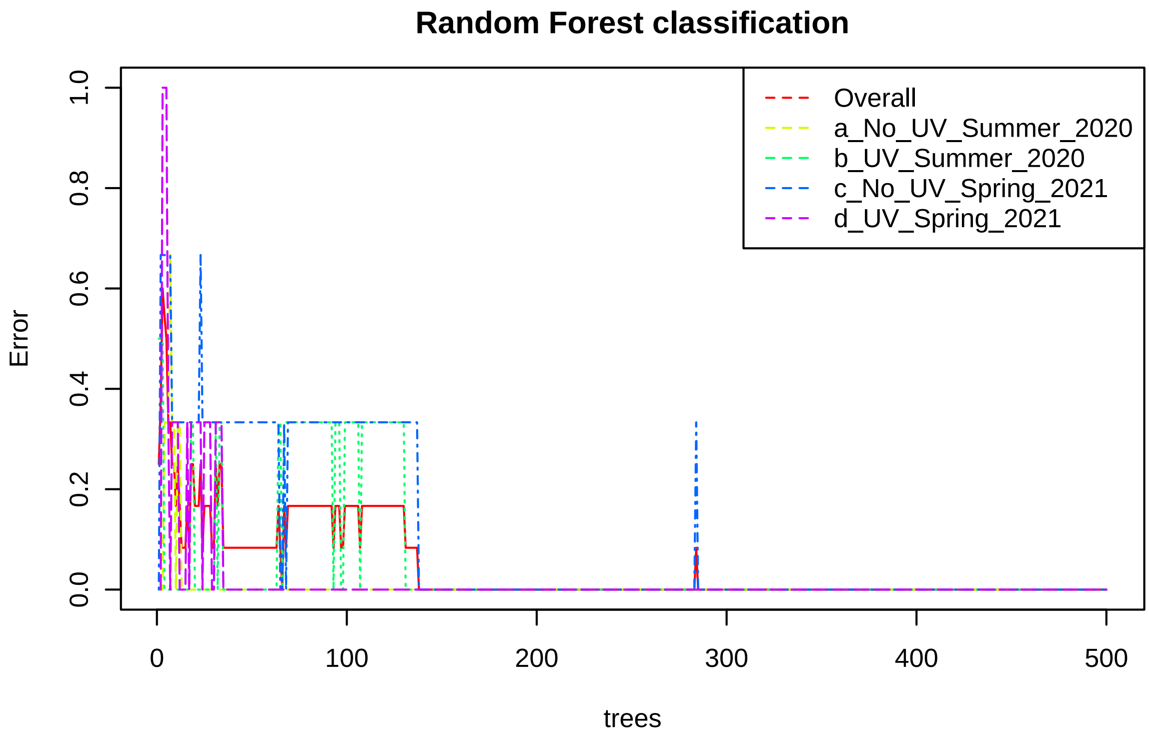


Figure 11: Cumulative error rates by Random Forest classification. The overall error rate is shown as the black line; the red and green lines represent the error rates for each class.

| | a_No_UV_Summer_2020 | b_UV_Summer_2020 | c_No_UV_Spring_2021 | d_UV_Spring_2021 | class.err[illegible] |
|---|---|---|---|---|---|
| a_No_UV_Summer_2020 | 3.00 | 0.00 | 0.00 | 0.00 | 0.[illegible] |
| b_UV_Summer_2020 | 0.00 | 3.00 | 0.00 | 0.00 | 0.[illegible] |
| c_No_UV_Spring_2021 | 0.00 | 0.00 | 3.00 | 0.00 | 0.[illegible] |
| d_UV_Spring_2021 | 0.00 | 0.00 | 0.00 | 3.00 | 0.[illegible] |

Table 4: Random Forest Classification Performance

[4]Andy Liaw and Matthew Wiener. *Classification and Regression by randomForest*, 2002, R News

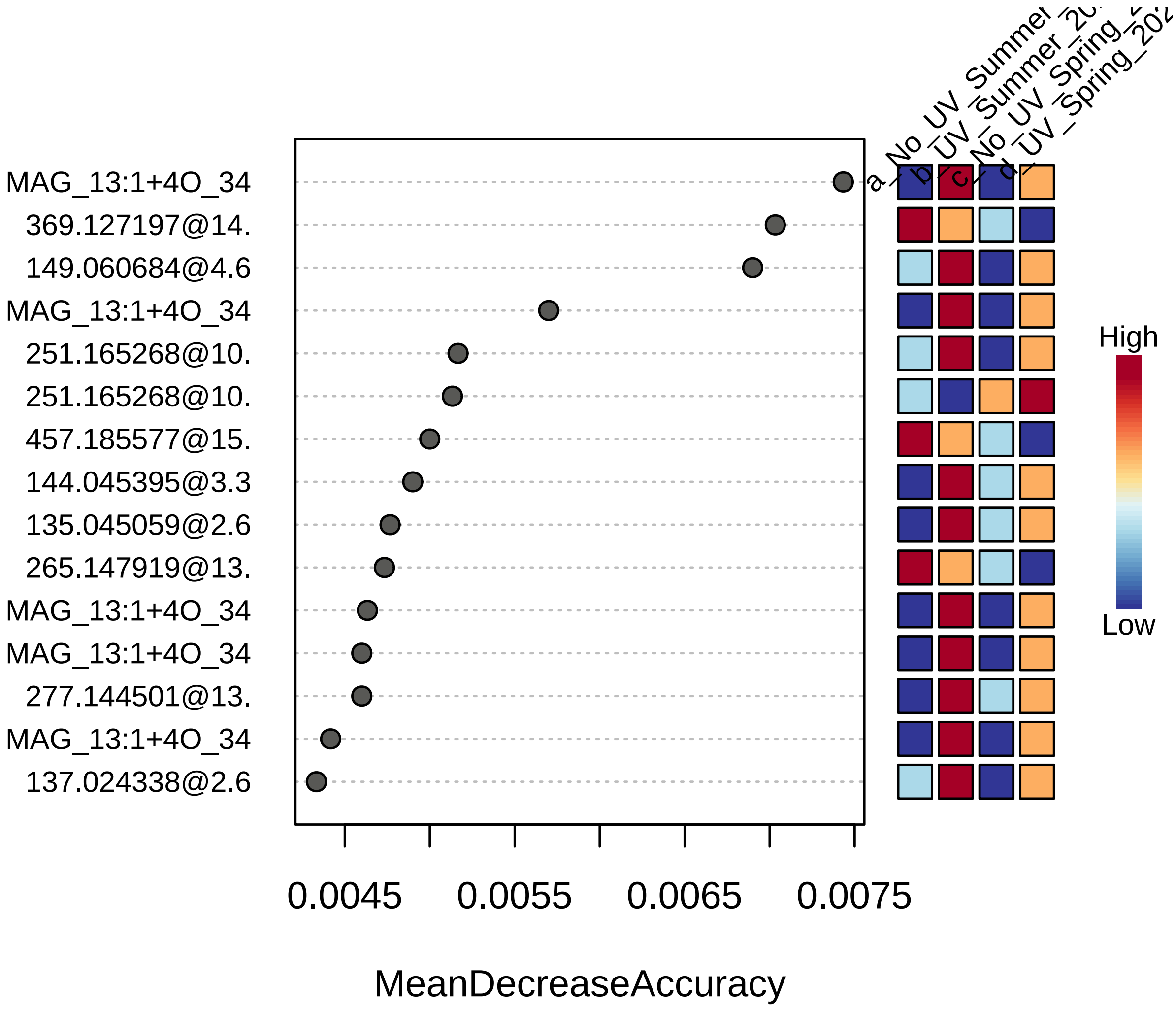


Figure 12: Significant features identified by Random Forest. The features are ranked by the mean decrease in classification accuracy when they are permuted.

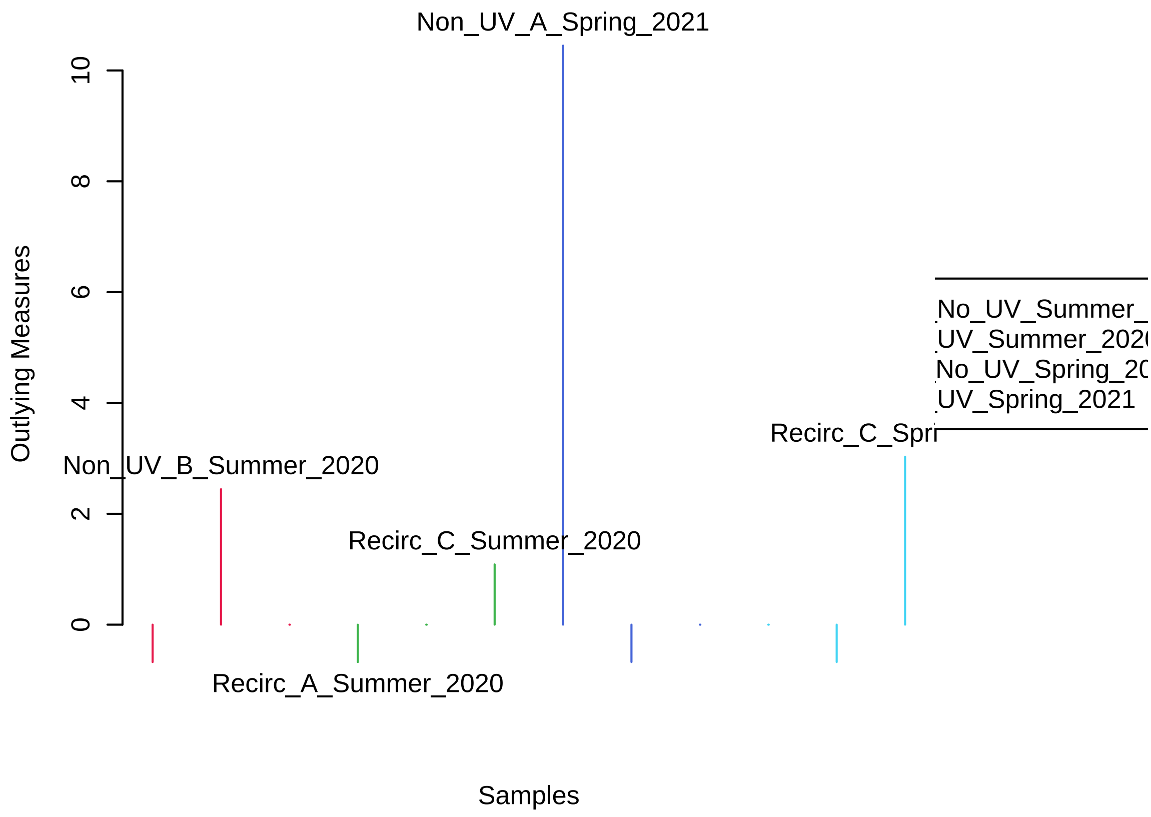


Figure 13: Potential outliers identified by Random Forest. Only the top five are labeled.

### 3 Appendix: R Command History

```
 [1] "mSet<-InitDataObjects(\"pktable\", \"stat\", FALSE)"
 [2] "mSet<-Read.TextData(mSet, \"Replacing_with_your_file_path\", \"colu\", \"disc\");"
 [3] "mSet<-SanityCheckData(mSet)"
 [4] "mSet<-ReplaceMin(mSet);"
 [5] "mSet<-SanityCheckData(mSet)"
 [6] "mSet<-FilterVariable(mSet, \"none\", \"F\", 25)"
 [7] "mSet<-PreparePrenormData(mSet)"
 [8] "mSet<-Normalization(mSet, \"NULL\", \"LogNorm\", \"MeanCenter\", ratio=FALSE, ratioNum=20)"
 [9] "mSet<-PlotNormSummary(mSet, \"norm_0_\", \"png\", 72, width=NA)"
[10] "mSet<-PlotSampleNormSummary(mSet, \"snorm_0_\", \"png\", 72, width=NA)"
[11] "mSet<-ANOVA.Anal(mSet, F, 0.05, \"fisher\", FALSE)"
[12] "mSet<-PlotANOVA(mSet, \"aov_0_\", \"png\", 72, width=NA)"
[13] "mSet<-UpdateLoadingCmpd(mSet, \"MAG_13:1+40_349.186859@8.604212\")"
[14] "mSet<-PlotCmpdSummary(mSet, \"MAG_13:1+40_349.186859@8.604212\",\"NA\", 0, \"png\", 72, width=
[15] "mSet<-UpdateLoadingCmpd(mSet, \"MAG_13:1+40_349.186859@8.494180\")"
[16] "mSet<-PlotCmpdSummary(mSet, \"MAG_13:1+40_349.186859@8.494180\",\"NA\", 1, \"png\", 72, width=
[17] "mSet<-UpdateLoadingCmpd(mSet, \"MAG_13:1+40_349.186859@8.604212\")"
[18] "mSet<-PlotCmpdSummary(mSet, \"MAG_13:1+40_349.186859@8.604212\",\"NA\", 2, \"png\", 72, width=
[19] "mSet<-UpdateLoadingCmpd(mSet, \"MAG_13:1+40_349.186798@8.338817\")"
[20] "mSet<-PlotCmpdSummary(mSet, \"MAG_13:1+40_349.186798@8.338817\",\"NA\", 3, \"png\", 72, width=
[21] "mSet<-UpdateLoadingCmpd(mSet, \"MAG_13:1+40_349.186859@8.759778\")"
[22] "mSet<-PlotCmpdSummary(mSet, \"MAG_13:1+40_349.186859@8.759778\",\"NA\", 4, \"png\", 72, width=
[23] "mSet<-SPLSR.Anal(mSet, 5, 10, \"same\", \"Mfold\")"
[24] "mSet<-PlotSPLSPairSummary(mSet, \"spls_pair_0_\", \"png\", 72, width=NA, 5)"
[25] "mSet<-PlotSPLS2DScore(mSet, \"spls_score2d_0_\", \"png\", 72, width=NA, 1,2,0.95,0,0)"
[26] "mSet<-PlotSPLS3DScoreImg(mSet, \"spls_score3d_0_\", \"png\", 72, width=NA, 1,2,3, 40)"
[27] "mSet<-PlotSPLSLoading(mSet, \"spls_loading_0_\", \"png\", 72, width=NA, 1,\"overview\");"
[28] "mSet<-PlotSPLSDA.Classification(mSet, \"spls_cv_0_\", \"png\", 72, width=NA)"
[29] "mSet<-PlotSPLS3DLoading(mSet, \"spls_loading3d_0_\", \"json\", 1,2,3)"
[30] "mSet<-PlotSPLSLoading(mSet, \"spls_loading_1_\", \"png\", 72, width=NA, 2,\"overview\");"
[31] "mSet<-PlotHCTree(mSet, \"tree_0_\", \"png\", 72, width=NA, \"euclidean\", \"ward.D\")"
[32] "mSet<-PlotHeatMap(mSet, \"heatmap_0_\", \"png\", 72, width=NA, \"norm\", \"row\", \"euclidean\
[33] "mSet<-RF.Anal(mSet, 500,7,1)"
[34] "mSet<-PlotRF.Classify(mSet, \"rf_cls_0_\", \"png\", 72, width=NA)"
[35] "mSet<-PlotRF.VIP(mSet, \"rf_imp_0_\", \"png\", 72, width=NA)"
[36] "mSet<-PlotRF.Outlier(mSet, \"rf_outlier_0_\", \"png\", 72, width=NA)"
[37] "mSet<-ComputeDSPC(mSet)"
[38] "mSet<-CreateGraph(mSet)"
[39] "mSet<-ComputeDSPC(mSet)"
[40] "mSet<-CreateGraph(mSet)"
[41] "FilterNetByCor(1.0E-6, 0.0, -1.0, 0.0, 0.0, 1.0)"
[42] "FilterNetByCor(1.0E-6, 0.0, -1.0, -1.0, 0.0, 0.0)"
[43] "FilterNetByCor(1.0E-6, 0.0, -1.0, 0.0, 0.0, 1.0)"
[44] "FilterNetByCor(0.0, 0.0, -1.0, -1.0, 0.0, 0.0)"
[45] "mSet<-Kmeans.Anal(mSet, 3)"
[46] "mSet<-PlotKmeans(mSet, \"km_0_\", \"png\", 72, width=NA, \"default\", \"F\")"
[47] "mSet<-PlotClustPCA(mSet, \"km_pca_0_\", \"png\", 72, width=NA, \"default\", \"km\", \"F\")"
[48] "mSet<-SaveTransformedData(mSet)"
[49] "mSet<-PreparePDFReport(mSet, \"guest7036401620298107543\")\n"
```

---

The report was generated on Mon Jan 23 18:08:06 2023 with R version 4.2.2 (2022-10-31), OS system: Linux, version: -Ubuntu SMP Tue Nov 22 19:54:14 UTC 2022 .
