## Supplemental Report 2 for "Oxylipins and Other Natural Products Produced by UV-sterilization Impact Commercial Oyster Larvae Production in a New England Estuary"

### Metabolomic Data Analysis with MetaboAnalyst 5.0

Name: guest4008043714584141497

April 28, 2022

### 1 Data Processing and Normalization

#### 1.1 Reading and Processing the Raw Data

MetaboAnalyst accepts a variety of data types generated in metabolomic studies, including compound concentration data, binned NMR/MS spectra data, NMR/MS peak list data, as well as MS spectra (NetCDF, mzXML, mzDATA). Users need to specify the data types when uploading their data in order for MetaboAnalyst to select the correct algorithm to process them. Table 1 summarizes the result of the data processing steps.

##### 1.1.1 Reading Concentration Data

The concentration data should be uploaded in comma separated values (.csv) format. Samples can be in rows or columns, with class labels immediately following the sample IDs.

Samples are in columns and features in rows. The uploaded file is in comma separated values (.csv) format. The uploaded data file contains 12 (samples) by 19 (compounds) data matrix.

No data filtering was performed.

Table 1: Summary of data processing results

| | Features (positive) | Missing/Zero | Features (processed) |
|---|---|---|---|
| Non_UV_A_Summer_2020 | 17 | 2 | 19 |
| Non_UV_B_Summer_2020 | 17 | 2 | 19 |
| Non_UV_C_Summer_2020 | 13 | 6 | 19 |
| Recirc_A_Summer_2020 | 19 | 0 | 19 |
| Recirc_B_Summer_2020 | 19 | 0 | 19 |
| Recirc_C_Summer_2020 | 19 | 0 | 19 |
| Non_UV_A_Spring_2021 | 13 | 6 | 19 |
| Non_UV_B_Spring_2021 | 13 | 6 | 19 |
| Non_UV_C_Spring_2021 | 10 | 9 | 19 |
| Recirc_A_Spring_2021 | 8 | 11 | 19 |
| Recirc_B_Spring_2021 | 11 | 8 | 19 |
| Recirc_C_Spring_2021 | 9 | 10 | 19 |

[2] Hackstadt AJ, Hess AM. *Filtering for increased power for microarray data analysis*, BMC Bioinformatics. 2009; 10: 11.

#### 1.2 Data Normalization

The data is stored as a table with one sample per row and one variable (bin/peak/metabolite) per column. The normalization procedures implemented below are grouped into four categories. Sample specific normalization allows users to manually adjust concentrations based on biological inputs (i.e. volume, mass); row-wise normalization allows general-purpose adjustment for differences among samples; data transformation and scaling are two different approaches to make features more comparable. You can use one or combine both to achieve better results.

The normalization consists of the following options:

1. Row-wise procedures:
   - Sample specific normalization (i.e. normalize by dry weight, volume)
   - Normalization by the sum
   - Normalization by the sample median
   - Normalization by a reference sample (probabilistic quotient normalization)[3]
   - Normalization by a pooled or average sample from a particular group
   - Normalization by a reference feature (i.e. creatinine, internal control)
   - Quantile normalization
2. Data transformation :
   - Log transformation (base 10)
   - Square root transformation
   - Cube root transformation
3. Data scaling:
   - Mean centering (mean-centered only)
   - Auto scaling (mean-centered and divided by standard deviation of each variable)
   - Pareto scaling (mean-centered and divided by the square root of standard deviation of each variable)
   - Range scaling (mean-centered and divided by the value range of each variable)

Figure 1 shows the effects before and after normalization.

[3]Dieterle F, Ross A, Schlotterbeck G, Senn H. *Probabilistic quotient normalization as robust method to account for dilution of complex biological mixtures. Application in 1H NMR metabonomics*, 2006, Anal Chem 78 (13);4281 - 4290

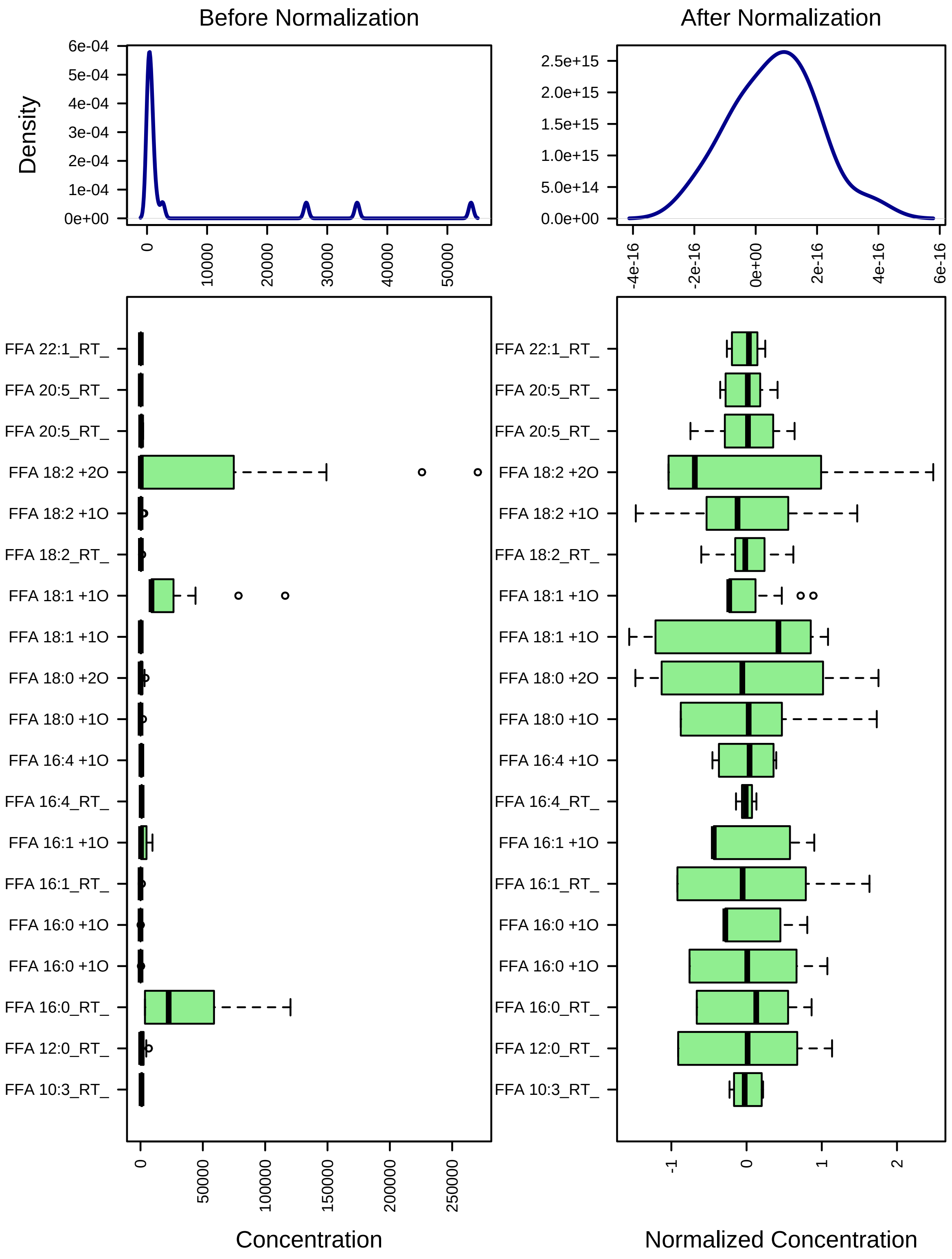


Figure 1: Box plots and kernel density plots before and after normalization. The boxplots show at most 50 features due to space limit. The density plots are based on all samples. Selected methods : Row-wise normalization: N/A; Data transformation: Log10 Normalization; Data scaling: Mean Centering.

#### 2 Statistical and Machine Learning Data Analysis

MetaboAnalyst offers a variety of methods commonly used in metabolomic data analyses. They include:

1. Univariate analysis methods:
   - Fold Change Analysis
   - T-tests
   - Volcano Plot
   - One-way ANOVA and post-hoc analysis
   - Correlation analysis
2. Multivariate analysis methods:
   - Principal Component Analysis (PCA)
   - Partial Least Squares - Discriminant Analysis (PLS-DA)
3. Robust Feature Selection Methods in microarray studies
   - Significance Analysis of Microarray (SAM)
   - Empirical Bayesian Analysis of Microarray (EBAM)
4. Clustering Analysis
   - Hierarchical Clustering
     - Dendrogram
     - Heatmap
   - Partitional Clustering
     - K-means Clustering
     - Self-Organizing Map (SOM)
5. Supervised Classification and Feature Selection methods
   - Random Forest
   - Support Vector Machine (SVM)

`Please note: some advanced methods are available only for two-group sample analyais.`

#### 2.1 One-way ANOVA

Univariate analysis methods are the most common methods used for exploratory data analysis. For multi-group analysis, MetaboAnalyst provides one-way Analysis of Variance (ANOVA). As ANOVA only tells whether the overall comparison is significant or not, it is usually followed by post-hoc analyses in order to identify which two levels are different. MetaboAnalyst provides two most commonly used methods for this purpose - Fisher's least significant difference method (Fisher's LSD) and Tukey's Honestly Significant Difference (Tukey's HSD). The univariate analyses provide a preliminary overview about features that are potentially significant in discriminating the conditions under study.

Figure 2 shows the important features identified by ANOVA analysis. Table 2 shows the details of these features. The `post-hoc Sig. Comparison` column shows the comparisons between different levels that are significant given the p value threshold.

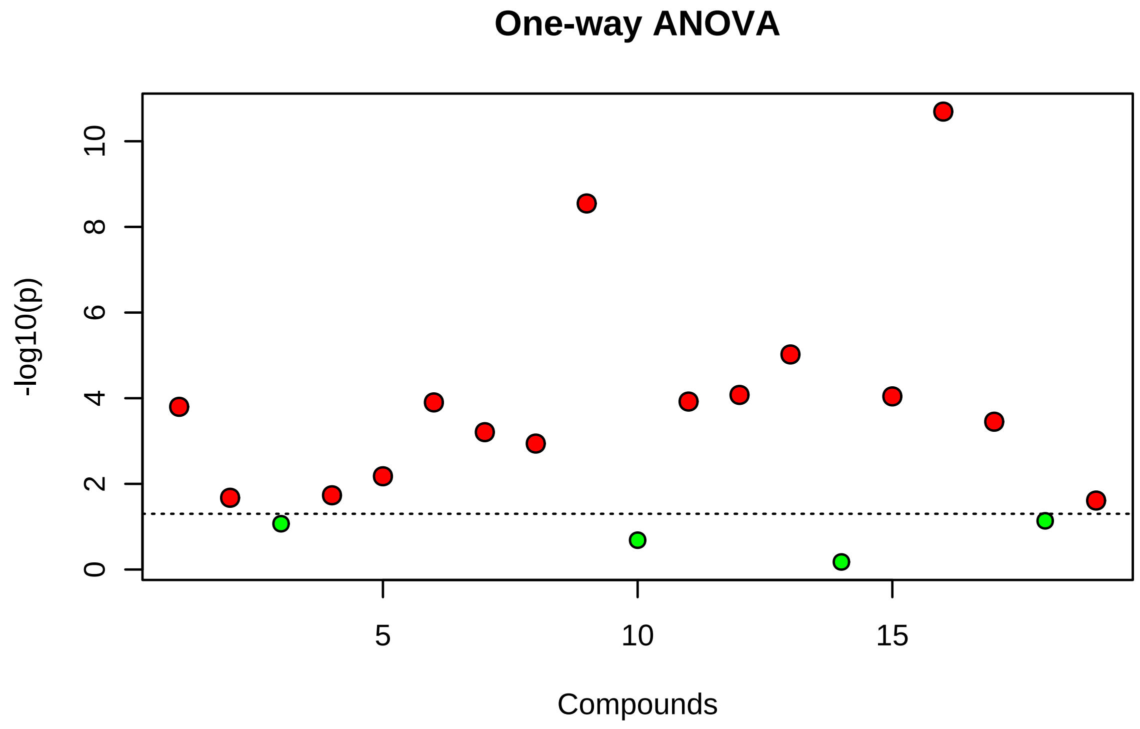


Figure 2: Important features selected by ANOVA plot with p value threshold 0.05.

Table 2: Important features identified by One-way ANOVA and post-hoc analysis

| | Compounds | f.value | p.value | -log10(p) | FDR | Fisher's LSD |
|---|---|---|---|---|---|---|
| 1 | FFA 18:2 +2O_RT_13.76_mz_311.2228 | 1570.30 | 0.00 | 10.69 | 0.00 | 2020_UV - 2020_nonUV; 2020_nonUV - 2021_nonUV; 2020_n |
| 2 | FFA 16:4 +1O_RT_12.31_mz_263.1652 | 454.73 | 0.00 | 8.55 | 0.00 | 2020_UV - 2020_nonUV; 2021_nonUV - 2020_nonUV; 2021_U |
| 3 | FFA 18:1 +1O_RT_18.39_mz_297.2435 | 57.20 | 0.00 | 5.02 | 0.00 | 2020_UV - 2020_nonUV; 2020_UV - 2021_nonUV; 2020_UV - |
| 4 | FFA 18:1 +1O_RT_16.58_mz_297.2435 | 31.96 | 0.00 | 4.08 | 0.00 | 2020_nonUV - 2021_UV; 2020_UV - 2021_nonUV; 2020_UV - |
| 5 | FFA 18:2 +1O_RT_17.18_mz_295.2279 | 31.28 | 0.00 | 4.04 | 0.00 | 2020_UV - 2020_nonUV; 2020_nonUV - 2021_UV; 2020_UV - |
| 6 | FFA 18:0 +2O_RT_14.78_mz_315.254 | 28.98 | 0.00 | 3.92 | 0.00 | 2020_UV - 2020_nonUV; 2020_nonUV - 2021_nonUV; 2020_n |
| 7 | FFA 16:1_RT_27.81_mz_253.2173 | 28.62 | 0.00 | 3.90 | 0.00 | 2020_UV - 2020_nonUV; 2020_nonUV - 2021_nonUV; 2020_n |
| 8 | FFA 10:3_RT_8.33_mz_165.0921 | 26.80 | 0.00 | 3.80 | 0.00 | 2020_nonUV - 2021_nonUV; 2020_nonUV - 2021_UV; 2020_U |
| 9 | FFA 20:5_RT_19.4_mz_301.2173 | 21.40 | 0.00 | 3.45 | 0.00 | 2020_nonUV - 2021_nonUV; 2020_nonUV - 2021_UV; 2020_U |
| 10 | FFA 16:1 +1O_RT_15.53_mz_269.2122 | 18.21 | 0.00 | 3.21 | 0.00 | 2020_UV - 2020_nonUV; 2020_nonUV - 2021_nonUV; 2020_n |
| 11 | FFA 16:4_RT_14.85_mz_247.1703 | 15.20 | 0.00 | 2.94 | 0.00 | 2021_nonUV - 2020_nonUV; 2021_UV - 2020_nonUV; 2021_n |
| 12 | FFA 16:0 +1O_RT_16.72_mz_271.2278 | 8.73 | 0.01 | 2.18 | 0.01 | 2020_UV - 2020_nonUV; 2020_UV - 2021_nonUV; 2020_UV - |
| 13 | FFA 16:0 +1O_RT_17.8_mz_271.2278 | 6.08 | 0.02 | 1.73 | 0.03 | 2020_UV - 2020_nonUV; 2020_UV - 2021_nonUV; 2020_UV - |
| 14 | FFA 12:0_RT_17.14_mz_199.1704 | 5.78 | 0.02 | 1.68 | 0.03 | 2020_UV - 2020_nonUV; 2020_UV - 2021_nonUV; 2020_UV - |
| 15 | FFA 22:1_RT_25.35_mz_337.3112 | 5.46 | 0.02 | 1.61 | 0.03 | 2021_nonUV - 2020_nonUV; 2021_UV - 2020_nonUV; 2021_n |

#### 2.2 Correlation Analysis

Correlation analysis can be used to visualize the overall correlations between different features It can also be used to identify which features are correlated with a feature of interest. Correlation analysis can also be used to identify if certain features show particular patterns under different conditions. Users first need to define a pattern in the form of a series of hyphenated numbers. For example, in a time-series study with four time points, a pattern of of `1-2-3-4` is used to search compounds with increasing the concentration as time changes; while a pattern of `3-2-1-3` can be used to search compounds that decrease at first, then bounce back to the original level.

Figure 3 shows the overall correlation heatmap.

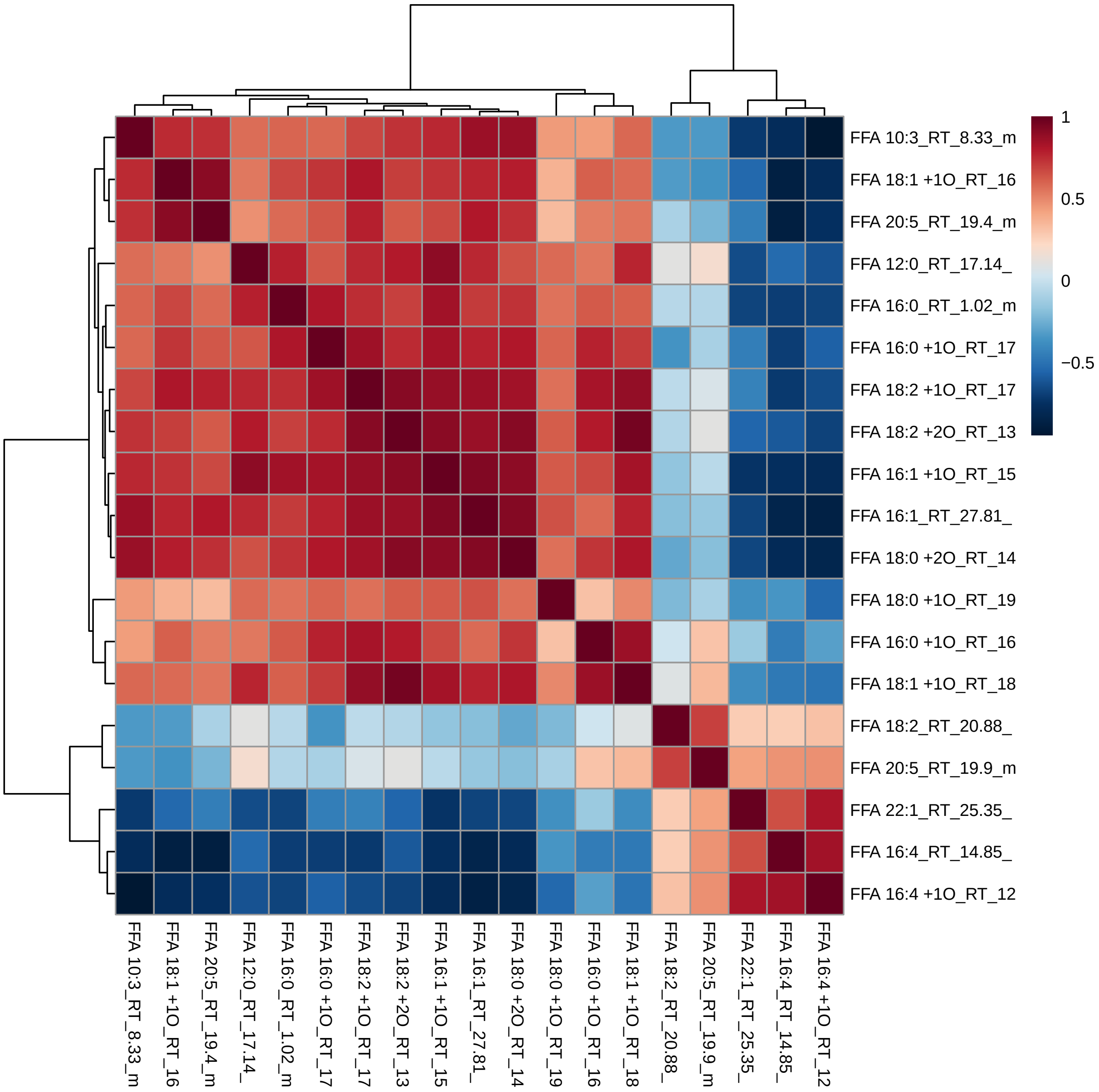


Figure 3: Correlation Heatmaps

#### 2.3 Principal Component Analysis (PCA)

PCA is an unsupervised method aiming to find the directions that best explain the variance in a data set (X) without referring to class labels (Y). The data are summarized into much fewer variables called *scores* which are weighted average of the original variables. The weighting profiles are called *loadings*. The PCA analysis is performed using the `prcomp` package. The calculation is based on singular value decomposition.

The Rscript `chemometrics.R` is required. Figure 4 is pairwise score plots providing an overview of the various seperation patterns among the most significant PCs; Figure 5 is the scree plot showing the variances explained by the selected PCs; Figure 6 shows the 2-D scores plot between selected PCs; Figure 7 shows the 3-D scores plot between selected PCs; Figure 8 shows the loadings plot between the selected PCs; Figure 9 shows the biplot between the selected PCs.

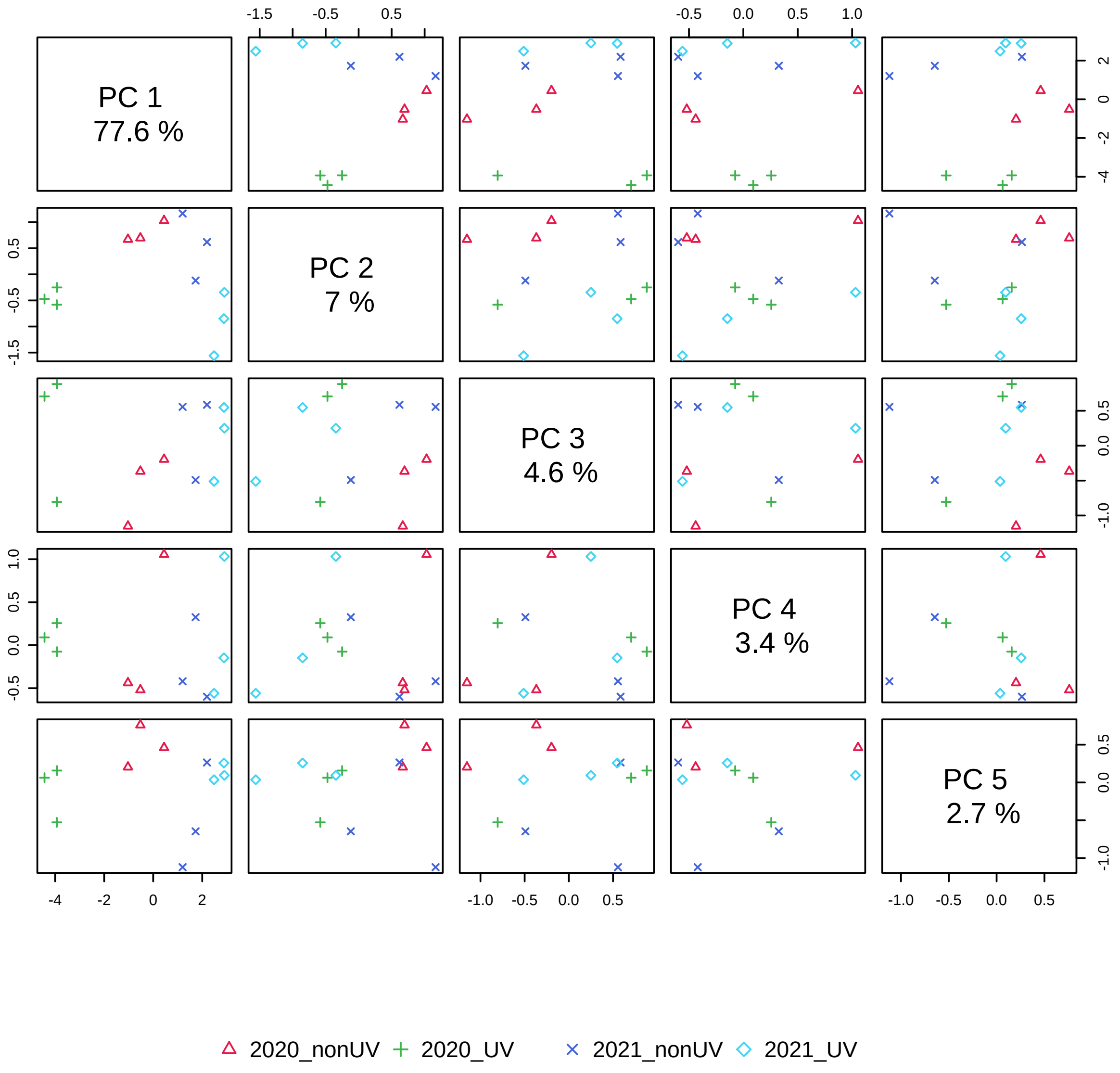


Figure 4: Pairwise score plots between the selected PCs. The explained variance of each PC is shown in the corresponding diagonal cell.

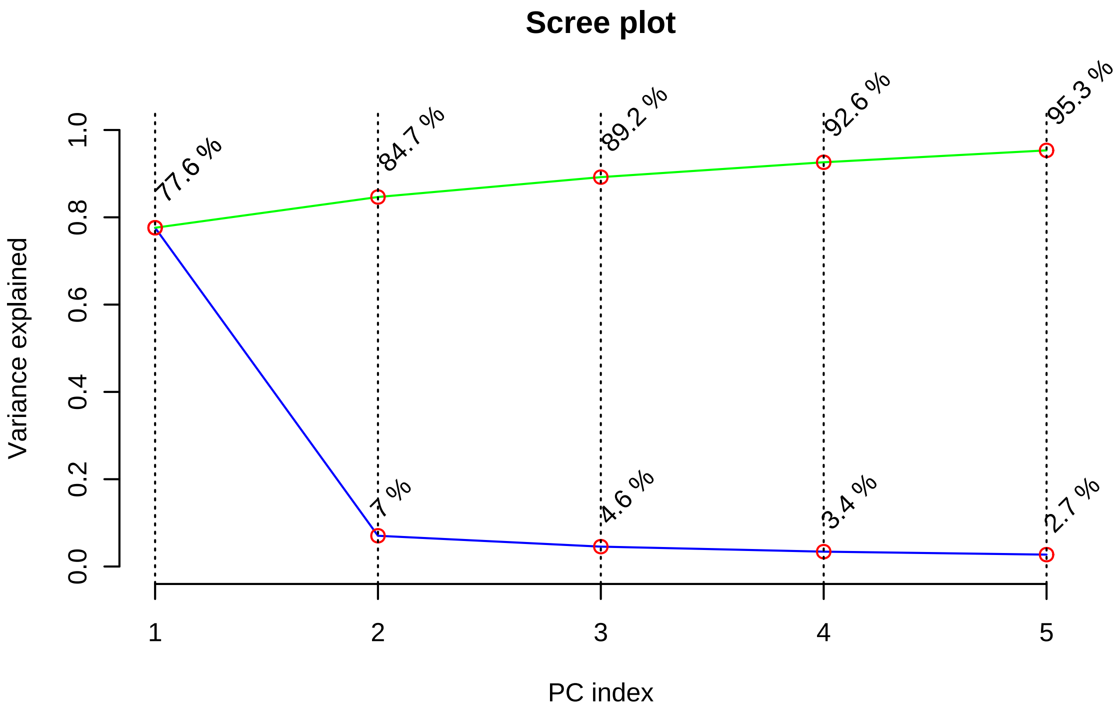


Figure 5: Scree plot shows the variance explained by PCs. The green line on top shows the accumulated variance explained; the blue line underneath shows the variance explained by individual PC.

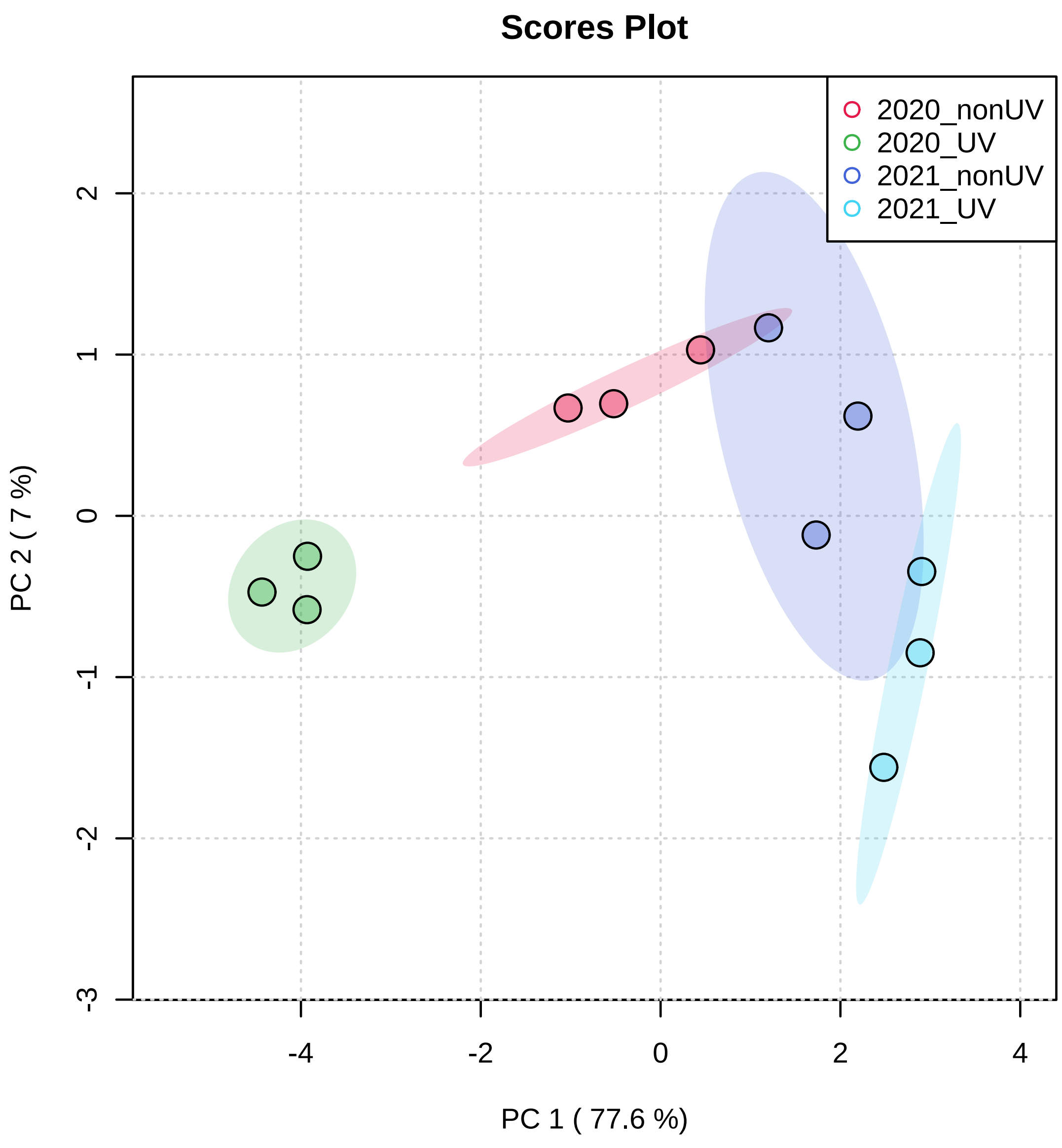


Figure 6: Scores plot between the selected PCs. The explained variances are shown in brackets.

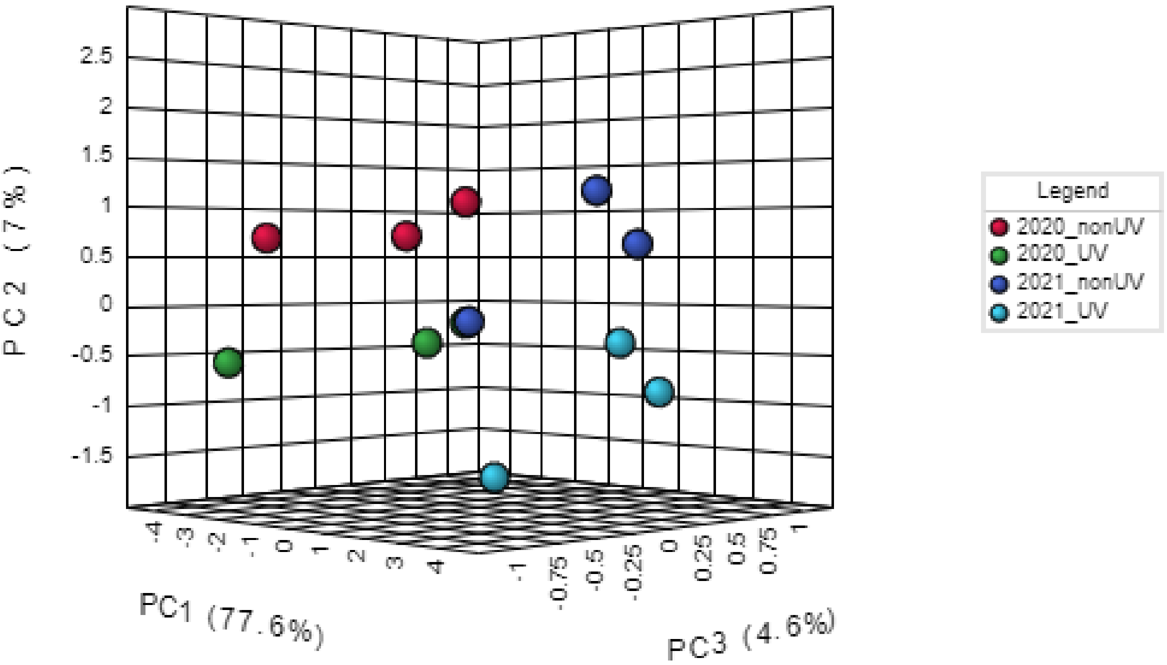


Figure 7: 3D score plot between the selected PCs. The explained variances are shown in brackets.

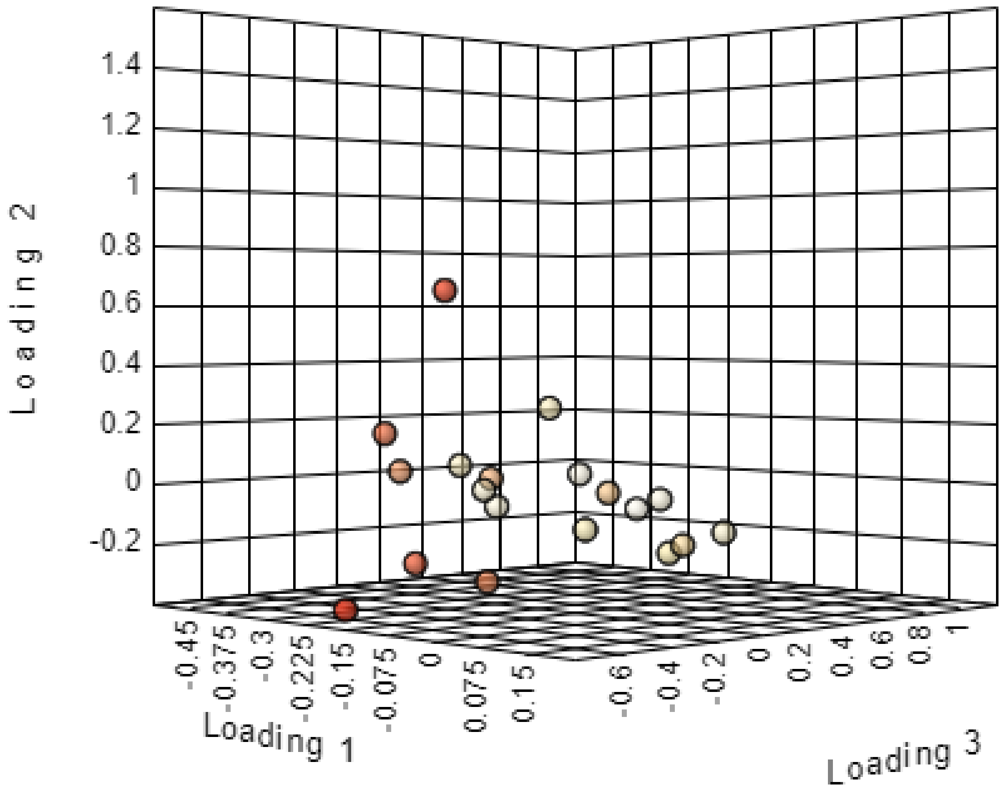


Figure 8: Loadings plot for the selected PCs.

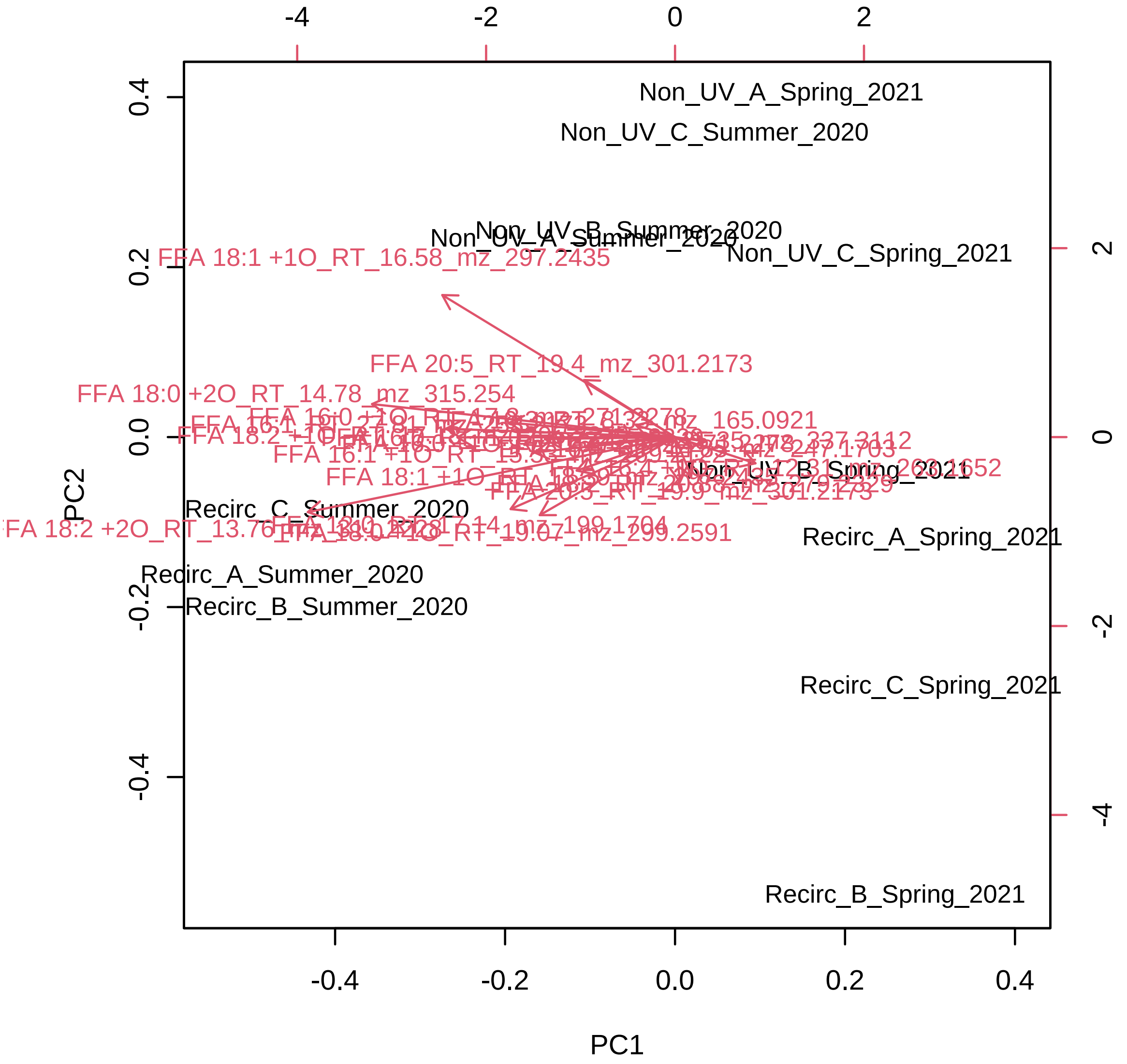


Figure 9: PCA biplot between the selected PCs. Note, you may want to test different centering and scaling normalization methods for the biplot to be displayed properly.

#### 2.4 Partial Least Squares - Discriminant Analysis (PLS-DA)

PLS is a supervised method that uses multivariate regression techniques to extract via linear combination of original variables (X) the information that can predict the class membership (Y). The PLS regression is performed using the `plsr` function provided by R `pls` package[4]. The classification and cross-validation are performed using the corresponding wrapper function offered by the `caret` package[5].

To assess the significance of class discrimination, a permutation test was performed. In each permutation, a PLS-DA model was built between the data (X) and the permuted class labels (Y) using the optimal number of components determined by cross validation for the model based on the original class assignment. MetaboAnalyst supports two types of test statistics for measuring the class discrimination. The first one is based on prediction accuracy during training. The second one is separation distance based on the ratio of the between group sum of the squares and the within group sum of squares (B/W-ratio). If the observed test statistic is part of the distribution based on the permuted class assignments, the class discrimination cannot be considered significant from a statistical point of view.[6].

There are two variable importance measures in PLS-DA. The first, Variable Importance in Projection (VIP) is a weighted sum of squares of the PLS loadings taking into account the amount of explained Y-variation in each dimension. Please note, VIP scores are calculated for each components. When more than components are used to calculate the feature importance, the average of the VIP scores are used. The other importance measure is based on the weighted sum of PLS-regression. The weights are a function of the reduction of the sums of squares across the number of PLS components. Please note, for multiple-group (more than two) analysis, the same number of predictors will be built for each group. Therefore, the coefficient of each feature will be different depending on which group you want to predict. The average of the feature coefficients are used to indicate the overall coefficient-based importance.

Figure 10 shows the overview of scores plots; Figure 11 shows the 2-D scores plot between selected components; Figure 12 shows the 3-D scores plot between selected components; Figure 13 shows the loading plot between the selected components;Figure 14 shows the classification performance with different number of components; Figure 15 shows the results of permutation test for model validation; Figure 16 shows important features identified by PLS-DA.

[4]Ron Wehrens and Bjorn-Helge Mevik.*pls: Partial Least Squares Regression (PLSR) and Principal Component Regression (PCR)*, 2007, R package version 2.1-0

[5]Max Kuhn. Contributions from Jed Wing and Steve Weston and Andre Williams.*caret: Classification and Regression Training*, 2008, R package version 3.45

[6]Bijlsma et al.*Large-Scale Human Metabolomics Studies: A Strategy for Data (Pre-) Processing and Validation*, Anal Chem. 2006, 78 567 - 574

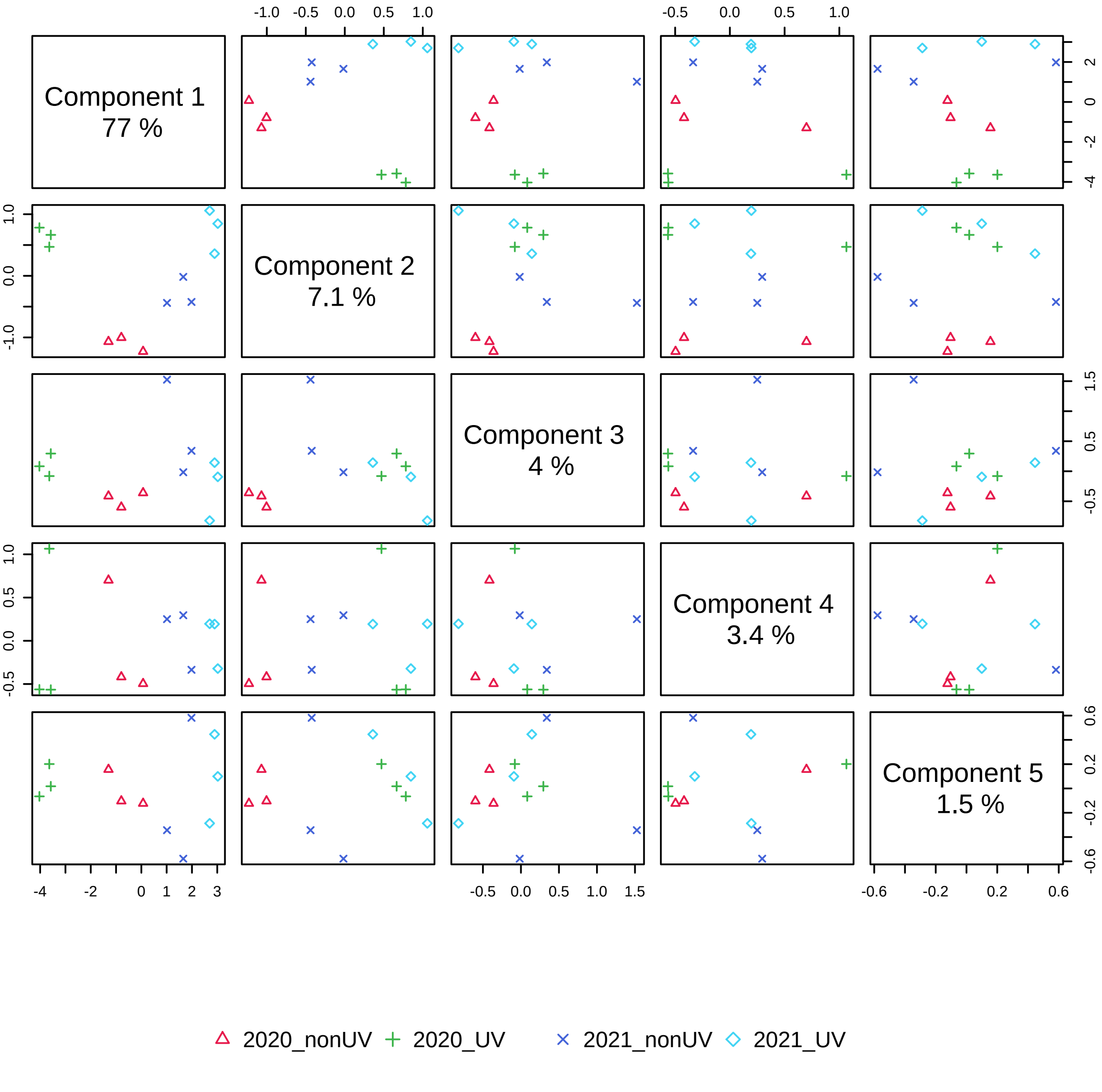


Figure 10: Pairwise scores plots between the selected components. The explained variance of each component is shown in the corresponding diagonal cell.

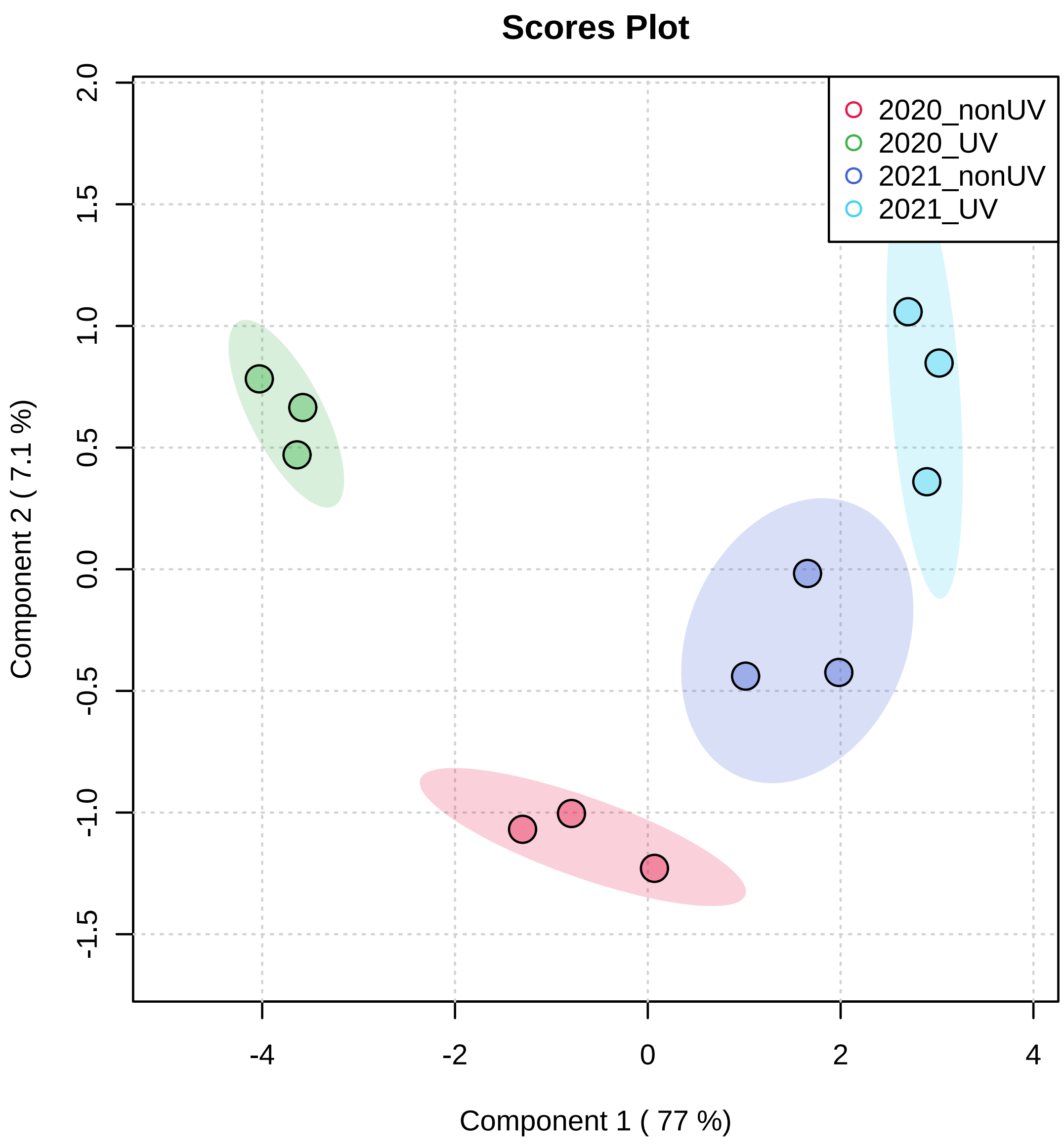


Figure 11: Scores plot between the selected PCs. The explained variances are shown in brackets.

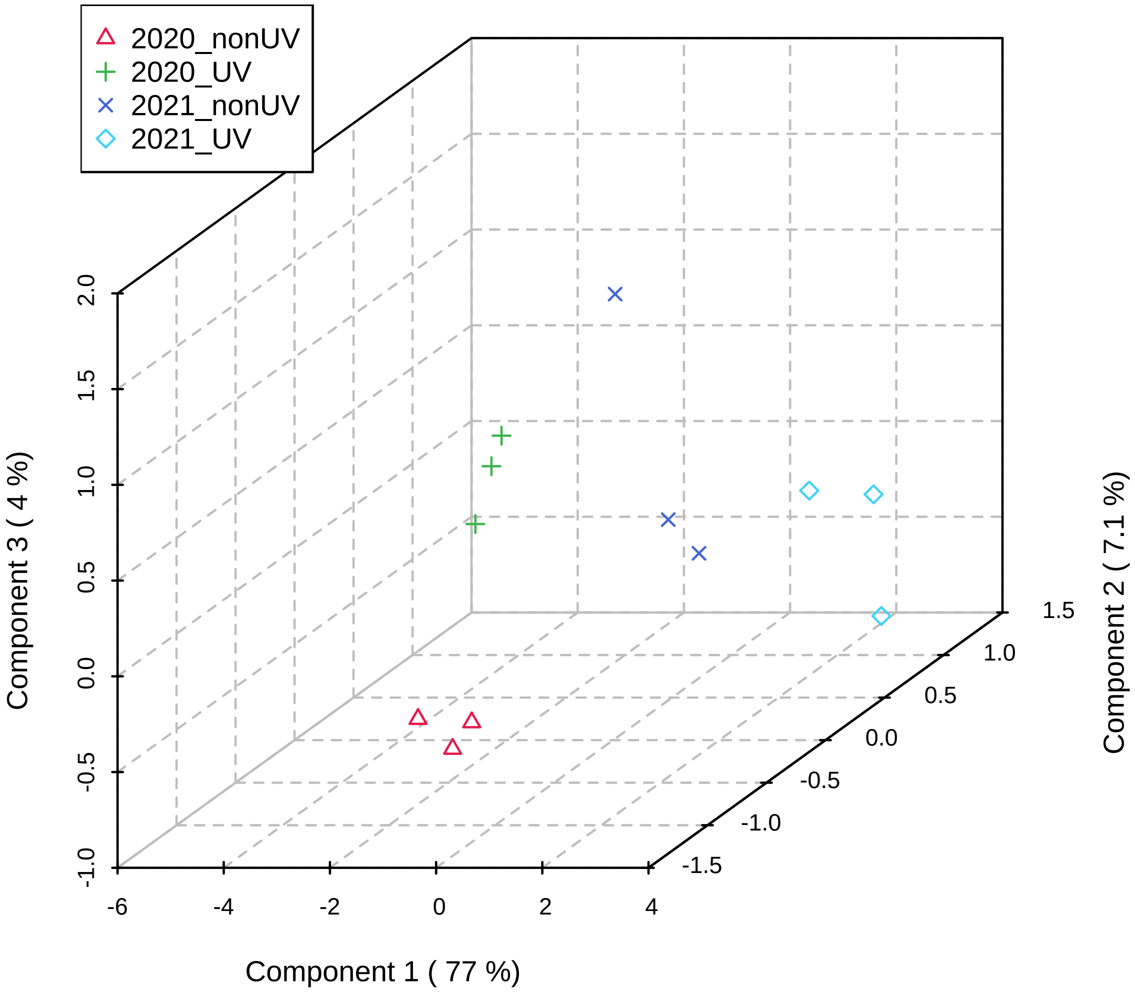


Figure 12: 3D scores plot between the selected PCs. The explained variances are shown in brackets.

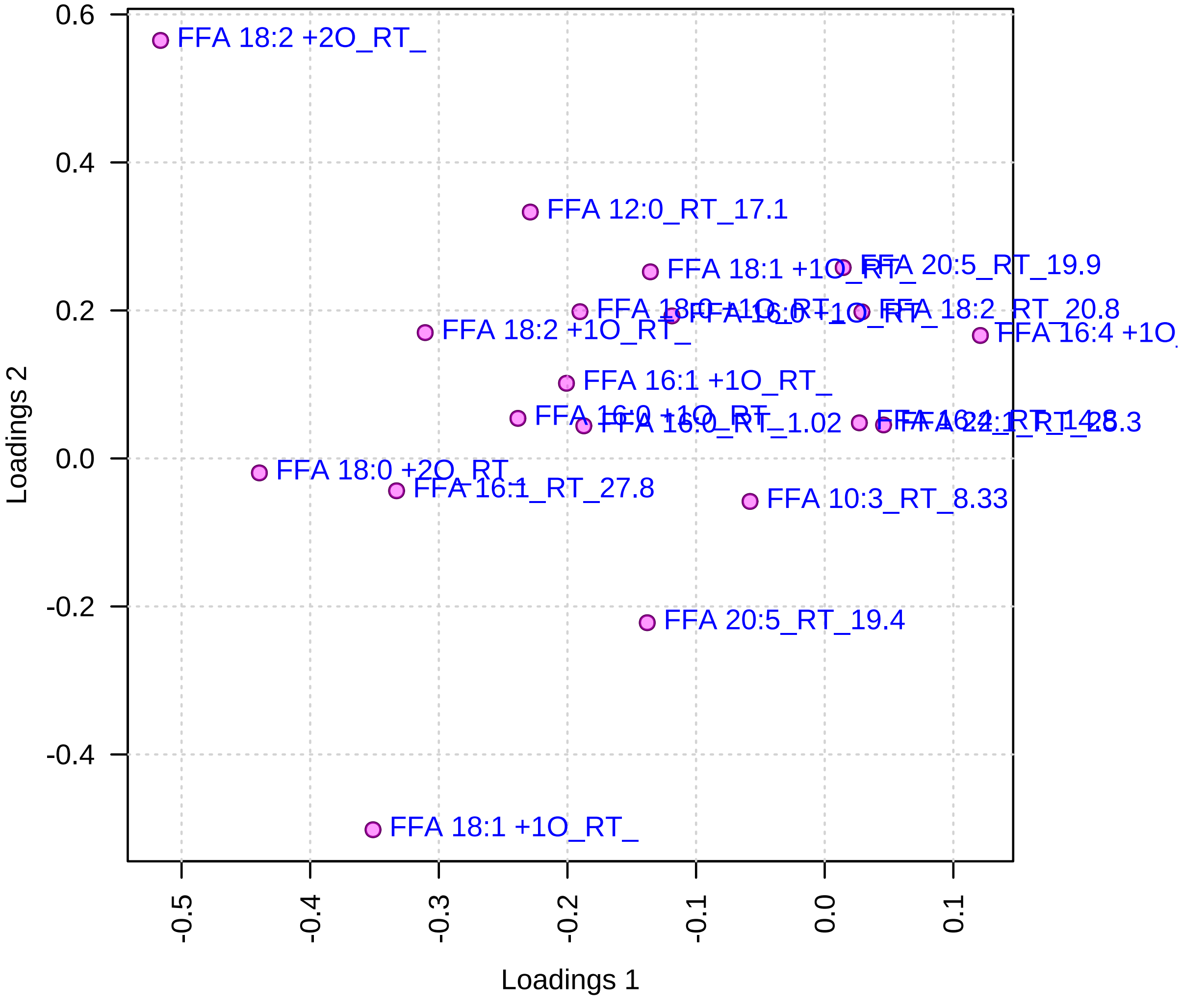


Figure 13: Loadings plot between the selected PCs.

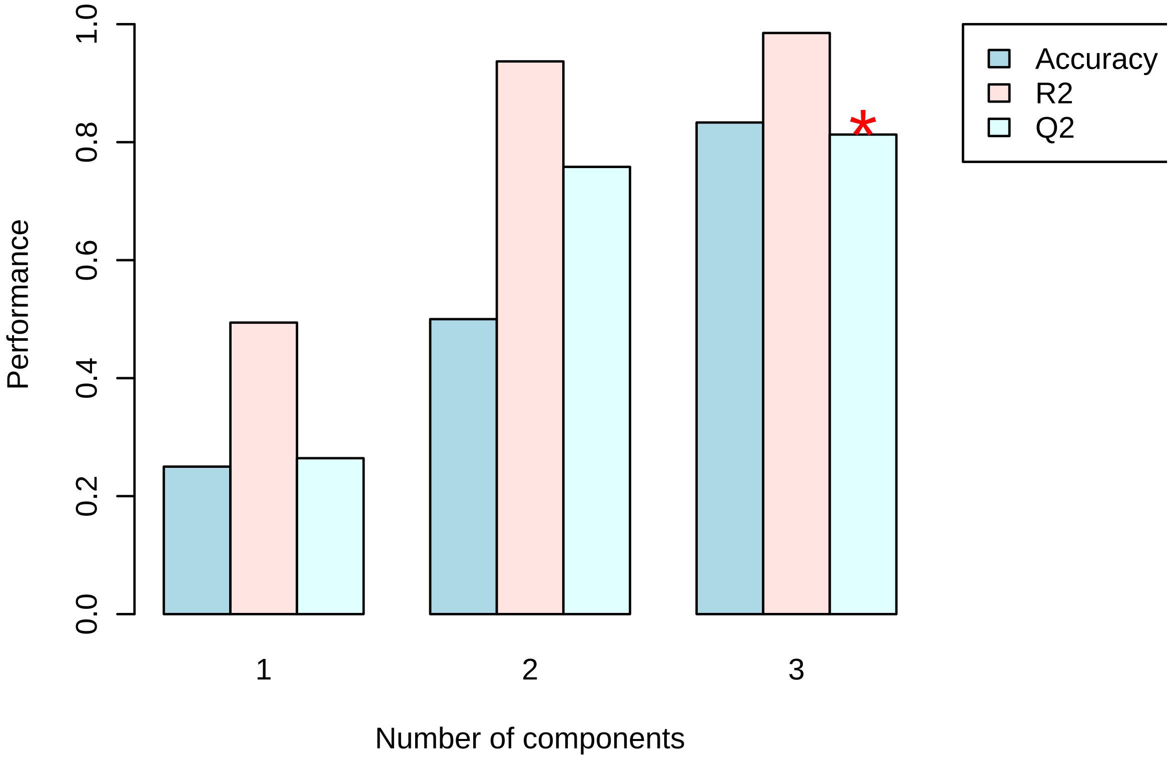


Figure 14: PLS-DA classification using different number of components. The red star indicates the best classifier.

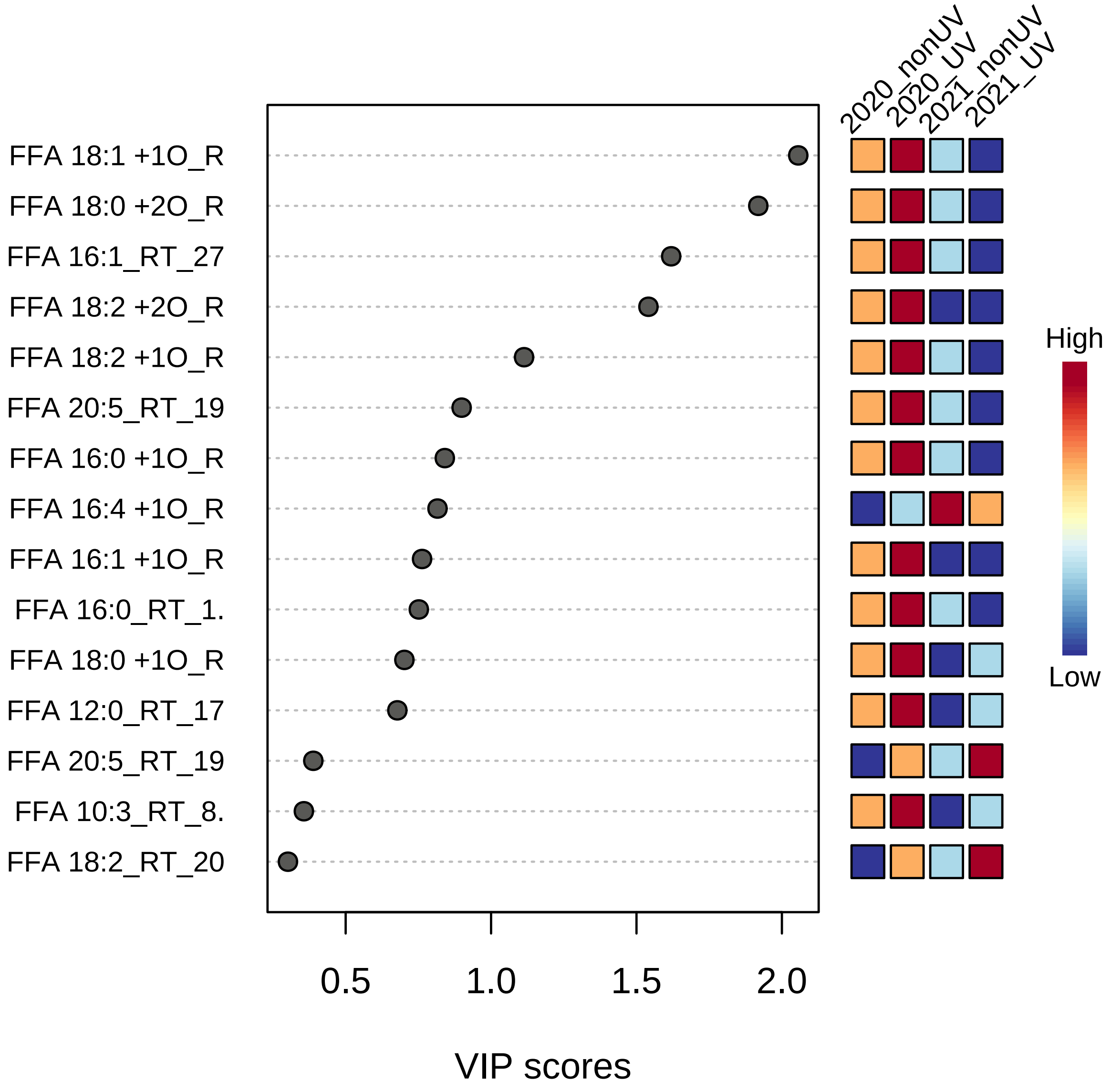


Figure 15: Important features identified by PLS-DA. The colored boxes on the right indicate the relative concentrations of the corresponding metabolite in each group under study.

#### 2.5 Hierarchical Clustering

In (agglomerative) hierarchical cluster analysis, each sample begins as a separate cluster and the algorithm proceeds to combine them until all samples belong to one cluster. Two parameters need to be considered when performing hierarchical clustering. The first one is similarity measure - Euclidean distance, Pearson's correlation, Spearman's rank correlation. The other parameter is clustering algorithms, including average linkage (clustering uses the centroids of the observations), complete linkage (clustering uses the farthest pair of observations between the two groups), single linkage (clustering uses the closest pair of observations) and Ward's linkage (clustering to minimize the sum of squares of any two clusters). Heatmap is often presented as a visual aid in addition to the dendrogram.

Hierachical clustering is performed with the `hclust` function in package `stat`. Figure 17 shows the clustering result in the form of a dendrogram. Figure 18 shows the clustering result in the form of a heatmap.

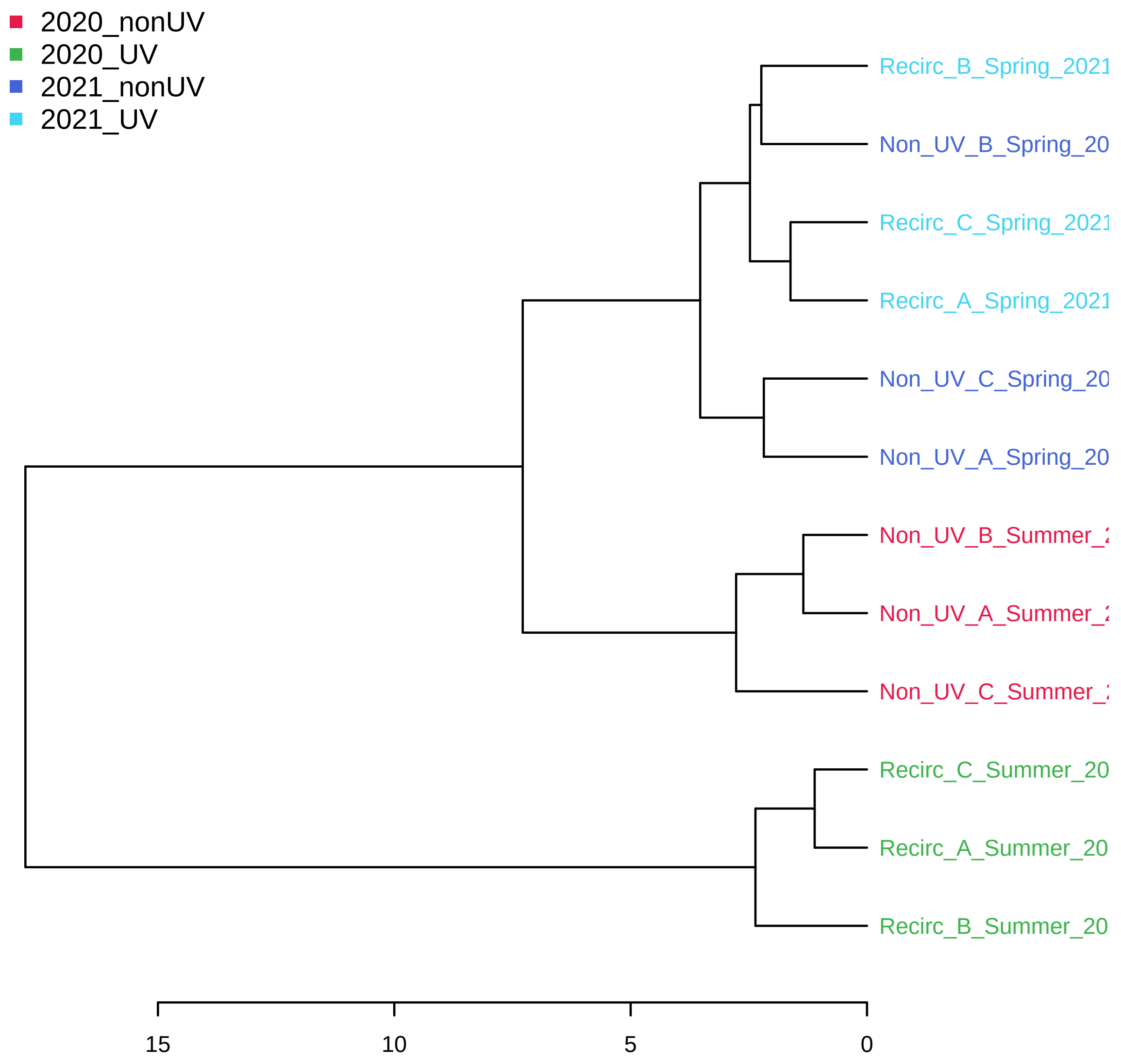


Figure 16: Clustering result shown as dendrogram (distance measure using `euclidean`, and clustering algorithm using `ward.D`).

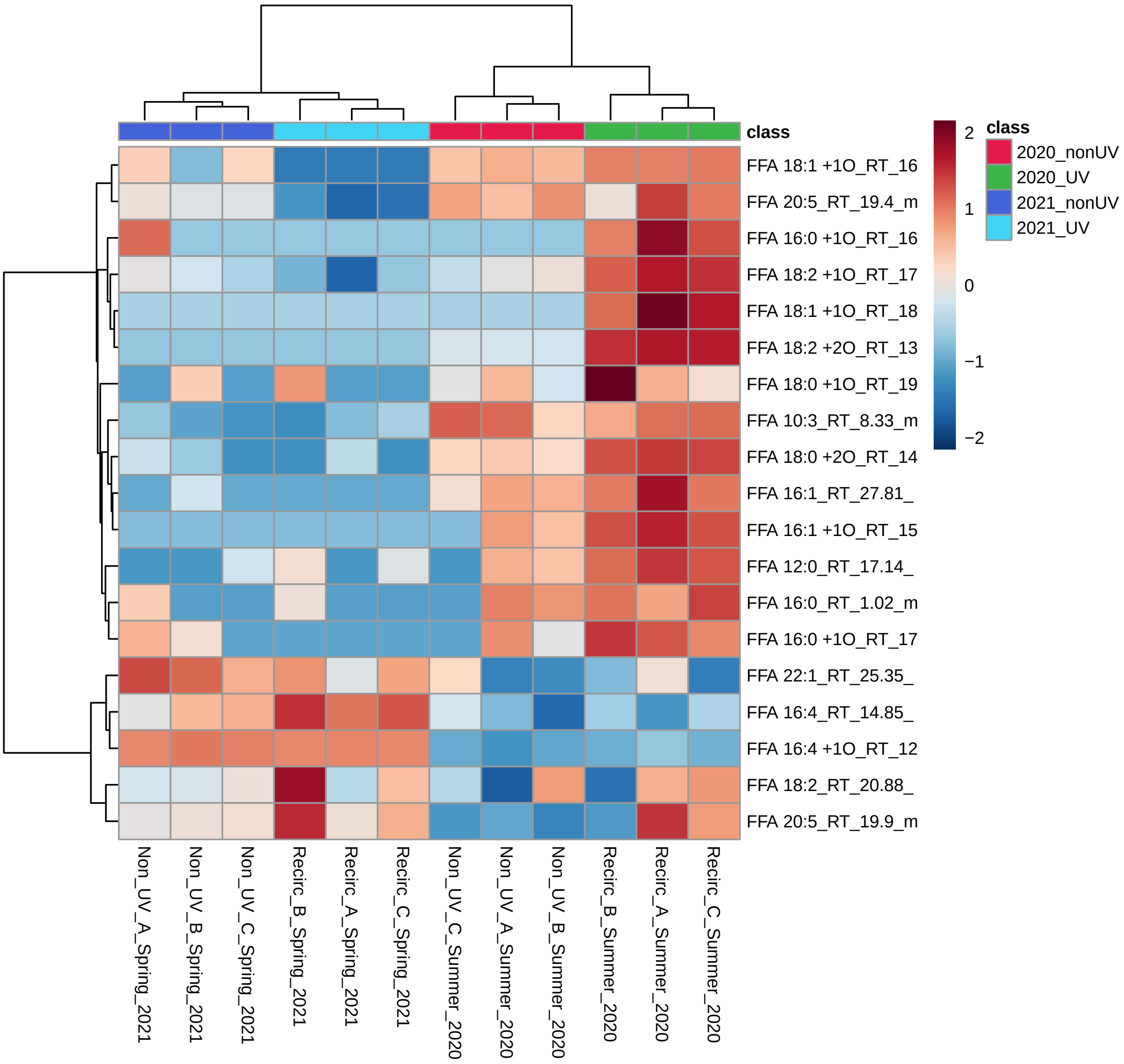


Figure 17: Clustering result shown as heatmap (distance measure using `euclidean`, and clustering algorithm using `ward.D`).

#### 2.6 K-means Clustering

K-means clustering is a nonhierarchical clustering technique. It begins by creating k random clusters (k is supplied by user). The program then calculates the mean of each cluster. If an observation is closer to the centroid of another cluster then the observation is made a member of that cluster. This process is repeated until none of the observations are reassigned to a different cluster.

K-means analysis is performed using the `kmeans` function in the package `stat`. Figure 19 shows clustering the results. Table 3 shows the members in each cluster from K-means analysis.

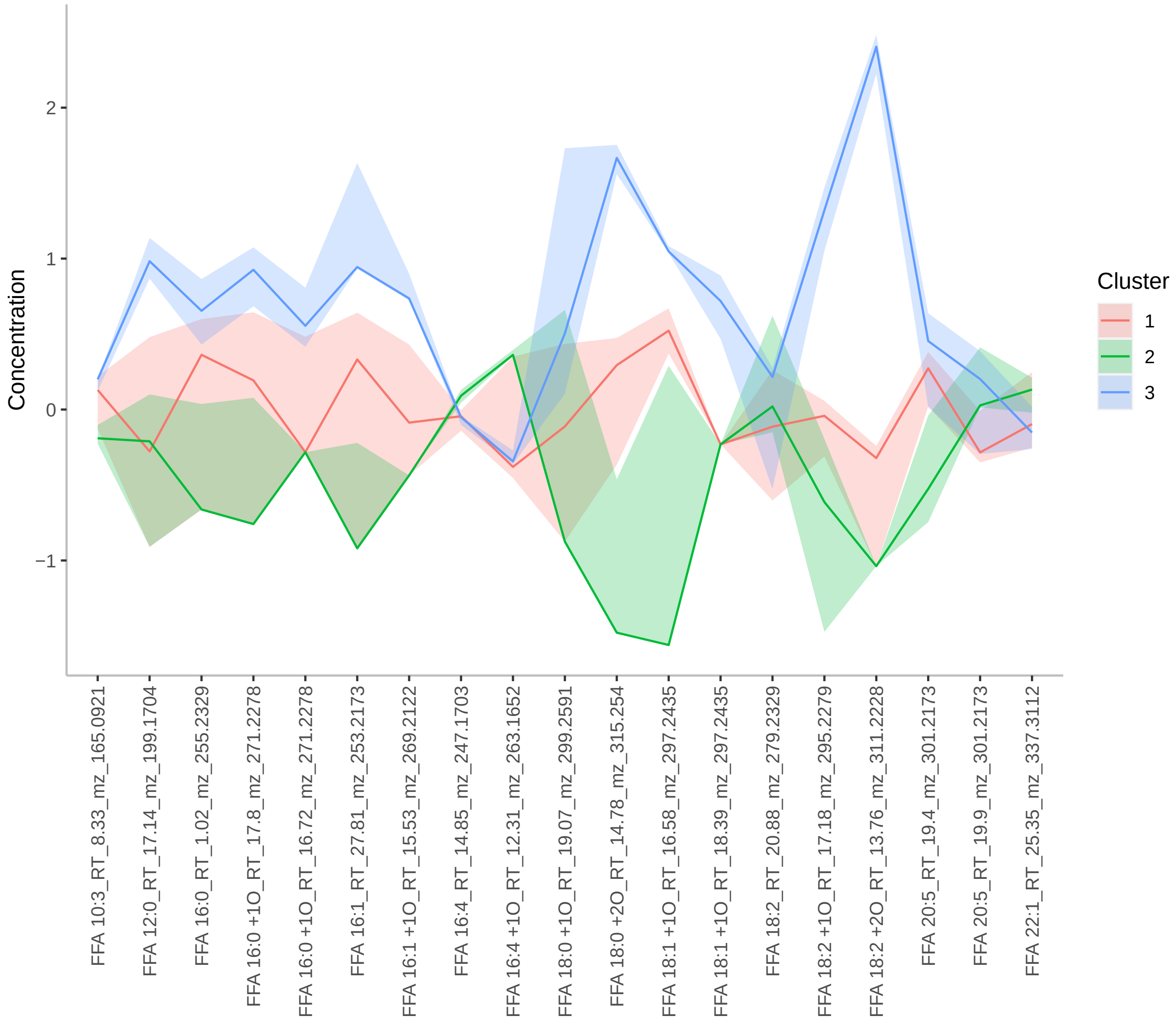


Figure 18: K-means cluster analysis. The x-axes are variable indices and y-axes are relative intensities. The blue lines represent median intensities of corresponding clusters

Table 3: Clustering result using K-means

| | Samples in each cluster |
|---|---|
| Cluster( 1 ) | Non_UV_A_Summer_2020 Non_UV_B_Summer_2020 Non_UV_C_Summer_2020 Non_UV_A_Spring_2021 |
| Cluster( 2 ) | Non_UV_B_Spring_2021 Non_UV_C_Spring_2021 Recirc_A_Spring_2021 Recirc_B_Spring_2021 Recirc_C_Spring_2021 |
| Cluster( 3 ) | Recirc_A_Summer_2020 Recirc_B_Summer_2020 Recirc_C_Summer_2020 |

#### 2.7 Random Forest (RF)

Random Forest is a supervised learning algorithm suitable for high dimensional data analysis. It uses an ensemble of classification trees, each of which is grown by random feature selection from a bootstrap sample at each branch. Class prediction is based on the majority vote of the ensemble. RF also provides other useful information such as OOB (out-of-bag) error, variable importance measure, and outlier measures. During tree construction, about one-third of the instances are left out of the bootstrap sample. This OOB data is then used as test sample to obtain an unbiased estimate of the classification error (OOB error). Variable importance is evaluated by measuring the increase of the OOB error when it is permuted. The outlier measures are based on the proximities during tree construction.

RF analysis is performed using the `randomForest` package[7]. Table 4 shows the confusion matrix of random forest. Figure 20 shows the cumulative error rates of random forest analysis for given parameters. Figure 21 shows the important features ranked by random forest. Figure 22 shows the outlier measures of all samples for the given parameters. The OOB error is 0.167

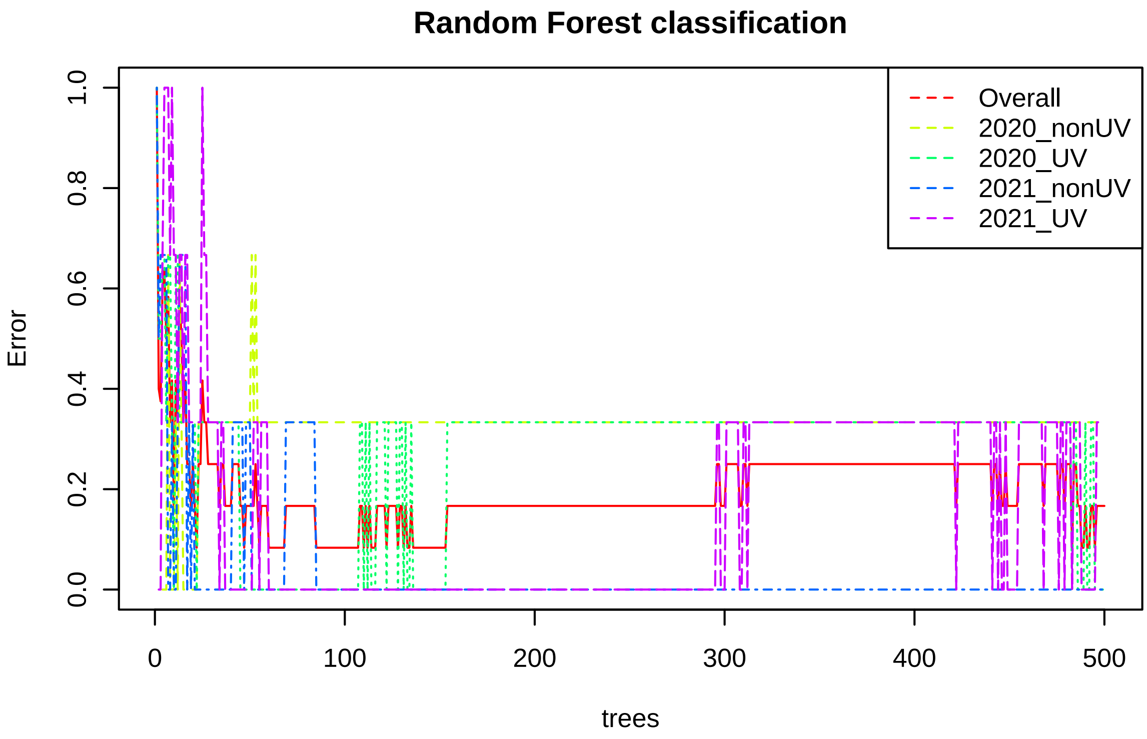


Figure 19: Cumulative error rates by Random Forest classification. The overall error rate is shown as the black line; the red and green lines represent the error rates for each class.

| | 2020_nonUV | 2020_UV | 2021_nonUV | 2021_UV | class.error |
|---|---|---|---|---|---|
| 2020_nonUV | 2.00 | 0.00 | 1.00 | 0.00 | 0.33 |
| 2020_UV | 0.00 | 3.00 | 0.00 | 0.00 | 0.00 |
| 2021_nonUV | 0.00 | 0.00 | 3.00 | 0.00 | 0.00 |
| 2021_UV | 0.00 | 0.00 | 1.00 | 2.00 | 0.33 |

Table 4: Random Forest Classification Performance

[7] Andy Liaw and Matthew Wiener. *Classification and Regression by randomForest*, 2002, R News

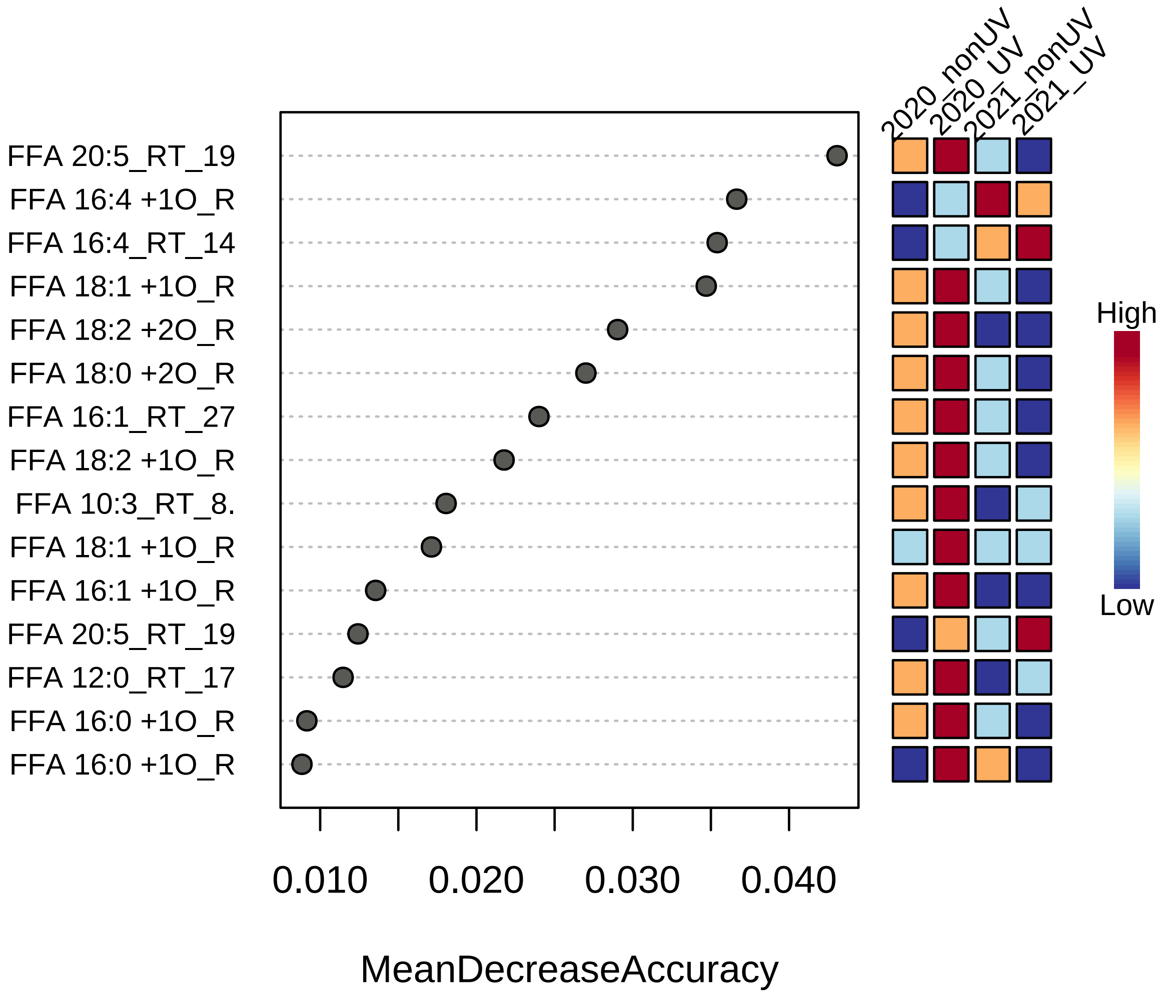


Figure 20: Significant features identified by Random Forest. The features are ranked by the mean decrease in classification accuracy when they are permuted.

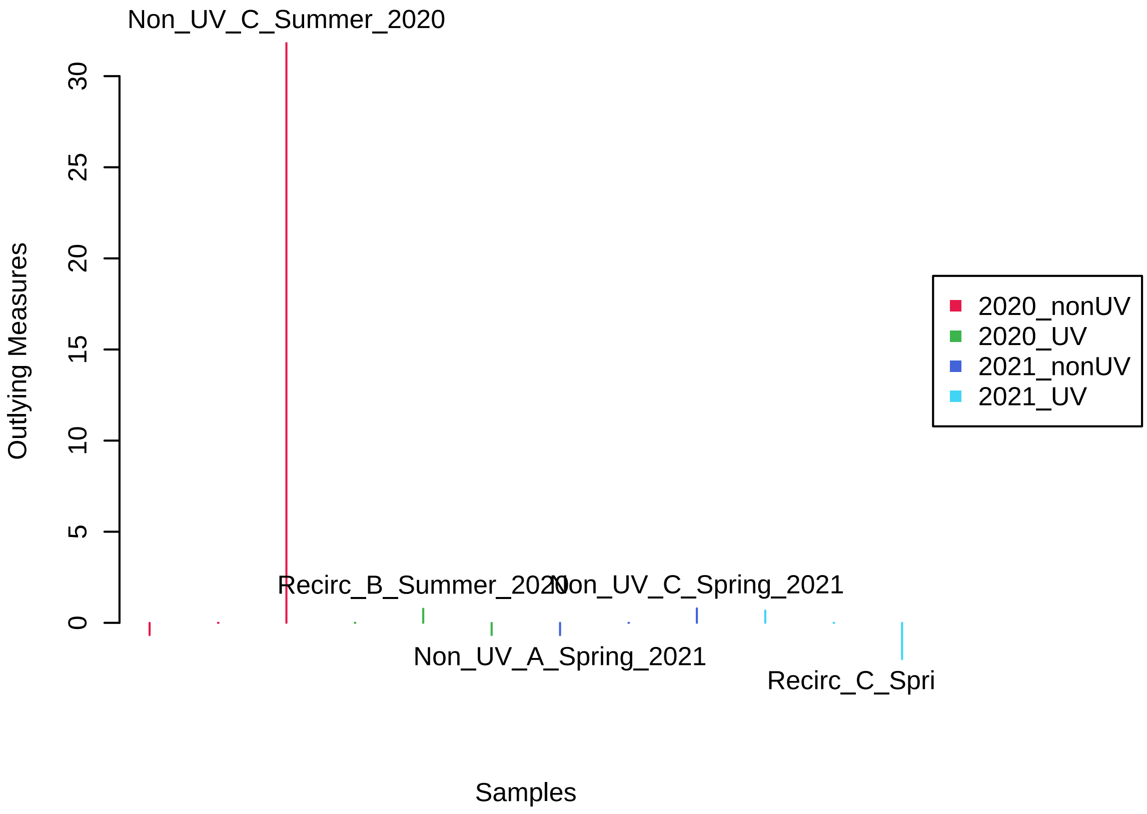


Figure 21: Potential outliers identified by Random Forest. Only the top five are labeled.

#### 3 Appendix: R Command History

```
 [1] "mSet<-InitDataObjects(\"conc\", \"stat\", FALSE)"
 [2] "mSet<-Read.TextData(mSet, \"Replacing_with_your_file_path\", \"colu\", \"disc\");"
 [3] "mSet<-SanityCheckData(mSet)"
 [4] "mSet<-ReplaceMin(mSet);"
 [5] "mSet<-PreparePrenormData(mSet)"
 [6] "mSet<-Normalization(mSet, \"NULL\", \"LogNorm\", \"MeanCenter\", ratio=FALSE, ratioNum=20)"
 [7] "mSet<-PlotNormSummary(mSet, \"norm_0_\", \"png\", 72, width=NA)"
 [8] "mSet<-PlotSampleNormSummary(mSet, \"snorm_0_\", \"png\", 72, width=NA)"
 [9] "mSet<-ANOVA.Anal(mSet, F, 0.05, \"fisher\", FALSE)"
[10] "mSet<-PlotANOVA(mSet, \"aov_0_\", \"png\", 72, width=NA)"
[11] "mSet<-UpdateLoadingCmpd(mSet, \"FFA 16:4 +1O_RT_12.31_mz_263.1652\")"
[12] "mSet<-PlotCmpdSummary(mSet, \"FFA 16:4 +1O_RT_12.31_mz_263.1652\",\"NA\", 0, \"png\", 72, widtl
[13] "mSet<-UpdateLoadingCmpd(mSet, \"FFA 18:2 +2O_RT_13.76_mz_311.2228\")"
[14] "mSet<-PlotCmpdSummary(mSet, \"FFA 18:2 +2O_RT_13.76_mz_311.2228\",\"NA\", 1, \"png\", 72, widtl
[15] "mSet<-UpdateLoadingCmpd(mSet, \"FFA 18:1 +1O_RT_18.39_mz_297.2435\")"
[16] "mSet<-PlotCmpdSummary(mSet, \"FFA 18:1 +1O_RT_18.39_mz_297.2435\",\"NA\", 2, \"png\", 72, widtl
[17] "mSet<-UpdateLoadingCmpd(mSet, \"FFA 10:3_RT_8.33_mz_165.0921\")"
[18] "mSet<-PlotCmpdSummary(mSet, \"FFA 10:3_RT_8.33_mz_165.0921\",\"NA\", 3, \"png\", 72, width=NA)
[19] "mSet<-UpdateLoadingCmpd(mSet, \"FFA 12:0_RT_17.14_mz_199.1704\")"
[20] "mSet<-PlotCmpdSummary(mSet, \"FFA 12:0_RT_17.14_mz_199.1704\",\"NA\", 4, \"png\", 72, width=NA
[21] "mSet<-UpdateLoadingCmpd(mSet, \"FFA 16:0_RT_1.02_mz_255.2329\")"
[22] "mSet<-PlotCmpdSummary(mSet, \"FFA 16:0_RT_1.02_mz_255.2329\",\"NA\", 5, \"png\", 72, width=NA)
[23] "mSet<-UpdateLoadingCmpd(mSet, \"FFA 16:0 +1O_RT_17.8_mz_271.2278\")"
[24] "mSet<-PlotCmpdSummary(mSet, \"FFA 16:0 +1O_RT_17.8_mz_271.2278\",\"NA\", 6, \"png\", 72, width
[25] "mSet<-UpdateLoadingCmpd(mSet, \"FFA 16:1_RT_27.81_mz_253.2173\")"
[26] "mSet<-PlotCmpdSummary(mSet, \"FFA 16:1_RT_27.81_mz_253.2173\",\"NA\", 7, \"png\", 72, width=NA
[27] "mSet<-UpdateLoadingCmpd(mSet, \"FFA 16:1_RT_27.81_mz_253.2173\")"
[28] "mSet<-PlotCmpdSummary(mSet, \"FFA 16:1_RT_27.81_mz_253.2173\",\"NA\", 8, \"png\", 72, width=NA
[29] "mSet<-UpdateLoadingCmpd(mSet, \"FFA 16:1 +1O_RT_15.53_mz_269.2122\")"
[30] "mSet<-PlotCmpdSummary(mSet, \"FFA 16:1 +1O_RT_15.53_mz_269.2122\",\"NA\", 9, \"png\", 72, widtl
[31] "mSet<-UpdateLoadingCmpd(mSet, \"FFA 16:1 +1O_RT_15.53_mz_269.2122\")"
[32] "mSet<-PlotCmpdSummary(mSet, \"FFA 16:1 +1O_RT_15.53_mz_269.2122\",\"NA\", 10, \"png\", 72, wid
[33] "mSet<-UpdateLoadingCmpd(mSet, \"FFA 16:4_RT_14.85_mz_247.1703\")"
[34] "mSet<-PlotCmpdSummary(mSet, \"FFA 16:4_RT_14.85_mz_247.1703\",\"NA\", 11, \"png\", 72, width=N
[35] "mSet<-UpdateLoadingCmpd(mSet, \"FFA 16:4 +1O_RT_12.31_mz_263.1652\")"
[36] "mSet<-PlotCmpdSummary(mSet, \"FFA 16:4 +1O_RT_12.31_mz_263.1652\",\"NA\", 12, \"png\", 72, wid
[37] "mSet<-UpdateLoadingCmpd(mSet, \"FFA 16:4_RT_14.85_mz_247.1703\")"
[38] "mSet<-PlotCmpdSummary(mSet, \"FFA 16:4_RT_14.85_mz_247.1703\",\"NA\", 13, \"png\", 72, width=N
[39] "mSet<-UpdateLoadingCmpd(mSet, \"FFA 18:0 +2O_RT_14.78_mz_315.254\")"
[40] "mSet<-PlotCmpdSummary(mSet, \"FFA 18:0 +2O_RT_14.78_mz_315.254\",\"NA\", 14, \"png\", 72, widtl
[41] "mSet<-UpdateLoadingCmpd(mSet, \"FFA 18:1 +1O_RT_16.58_mz_297.2435\")"
[42] "mSet<-PlotCmpdSummary(mSet, \"FFA 18:1 +1O_RT_16.58_mz_297.2435\",\"NA\", 15, \"png\", 72, wid
[43] "mSet<-UpdateLoadingCmpd(mSet, \"FFA 18:0 +2O_RT_14.78_mz_315.254\")"
[44] "mSet<-PlotCmpdSummary(mSet, \"FFA 18:0 +2O_RT_14.78_mz_315.254\",\"NA\", 16, \"png\", 72, widtl
[45] "mSet<-UpdateLoadingCmpd(mSet, \"FFA 18:1 +1O_RT_18.39_mz_297.2435\")"
[46] "mSet<-PlotCmpdSummary(mSet, \"FFA 18:1 +1O_RT_18.39_mz_297.2435\",\"NA\", 17, \"png\", 72, wid
[47] "mSet<-UpdateLoadingCmpd(mSet, \"FFA 18:1 +1O_RT_18.39_mz_297.2435\")"
[48] "mSet<-PlotCmpdSummary(mSet, \"FFA 18:1 +1O_RT_18.39_mz_297.2435\",\"NA\", 18, \"png\", 72, wid
[49] "mSet<-UpdateLoadingCmpd(mSet, \"FFA 18:2 +1O_RT_17.18_mz_295.2279\")"
[50] "mSet<-PlotCmpdSummary(mSet, \"FFA 18:2 +1O_RT_17.18_mz_295.2279\",\"NA\", 19, \"png\", 72, wid
[51] "mSet<-UpdateLoadingCmpd(mSet, \"FFA 18:2 +2O_RT_13.76_mz_311.2228\")"
[52] "mSet<-PlotCmpdSummary(mSet, \"FFA 18:2 +2O_RT_13.76_mz_311.2228\",\"NA\", 20, \"png\", 72, wid
[53] "mSet<-UpdateLoadingCmpd(mSet, \"FFA 18:2 +1O_RT_17.18_mz_295.2279\")"
[54] "mSet<-PlotCmpdSummary(mSet, \"FFA 18:2 +1O_RT_17.18_mz_295.2279\",\"NA\", 21, \"png\", 72, wid
[55] "mSet<-UpdateLoadingCmpd(mSet, \"FFA 20:5_RT_19.4_mz_301.2173\")"
[56] "mSet<-PlotCmpdSummary(mSet, \"FFA 20:5_RT_19.4_mz_301.2173\",\"NA\", 22, \"png\", 72, width=NA
```

```
[57] "mSet<-UpdateLoadingCmpd(mSet, \"FFA 22:1_RT_25.35_mz_337.3112\")"
[58] "mSet<-PlotCmpdSummary(mSet, \"FFA 22:1_RT_25.35_mz_337.3112\",\"NA\", 23, \"png\", 72, width=NA
[59] "mSet<-PlotCorrHeatMap(mSet, \"corr_0_\", \"png\", 72, width=NA, \"col\", \"pearson\", \"bwm\",
[60] "mSet<-ComputeDSPC(mSet)"
[61] "mSet<-CreateGraph(mSet)"
[62] "mSet<-ComputeDSPC(mSet)"
[63] "mSet<-CreateGraph(mSet)"
[64] "FilterNetByCor(0.0854, 0.0, -1.0, 0.0, 0.0, 1.0)"
[65] "mSet<-FilterBipartiNet(mSet, \"all\", 0.0, 1.0)"
[66] "mSet<-PCA.Anal(mSet)"
[67] "mSet<-PlotPCAPairSummary(mSet, \"pca_pair_0_\", \"png\", 72, width=NA, 5)"
[68] "mSet<-PlotPCAScree(mSet, \"pca_scree_0_\", \"png\", 72, width=NA, 5)"
[69] "mSet<-PlotPCA2DScore(mSet, \"pca_score2d_0_\", \"png\", 72, width=NA, 1,2,0.95,0,0)"
[70] "mSet<-PlotPCALoading(mSet, \"pca_loading_0_\", \"png\", 72, width=NA, 1,2);"
[71] "mSet<-PlotPCABiplot(mSet, \"pca_biplot_0_\", \"png\", 72, width=NA, 1,2)"
[72] "mSet<-PlotPCA3DLoading(mSet, \"pca_loading3d_0_\", \"json\", 1,2,3)"
[73] "mSet<-PLSR.Anal(mSet, reg=TRUE)"
[74] "mSet<-PlotPLSPairSummary(mSet, \"pls_pair_0_\", \"png\", 72, width=NA, 5)"
[75] "mSet<-PlotPLS2DScore(mSet, \"pls_score2d_0_\", \"png\", 72, width=NA, 1,2,0.95,0,0)"
[76] "mSet<-PlotPLS3DScoreImg(mSet, \"pls_score3d_0_\", \"png\", 72, width=NA, 1,2,3, 40)"
[77] "mSet<-PlotPLSLoading(mSet, \"pls_loading_0_\", \"png\", 72, width=NA, 1, 2);"
[78] "mSet<-PlotPLS3DLoading(mSet, \"pls_loading3d_0_\", \"json\", 1,2,3)"
[79] "mSet<-PLSDA.CV(mSet, \"L\",3, \"Q2\")"
[80] "mSet<-PlotPLS.Classification(mSet, \"pls_cv_0_\", \"png\", 72, width=NA)"
[81] "mSet<-PlotPLS.Imp(mSet, \"pls_imp_0_\", \"png\", 72, width=NA, \"vip\", \"Comp. 1\", 15,FALSE)
[82] "mSet<-PlotHCTree(mSet, \"tree_0_\", \"png\", 72, width=NA, \"euclidean\", \"ward.D\")"
[83] "mSet<-PlotHeatMap(mSet, \"heatmap_0_\", \"png\", 72, width=NA, \"norm\", \"row\", \"euclidean\
[84] "mSet<-Kmeans.Anal(mSet, 3)"
[85] "mSet<-PlotKmeans(mSet, \"km_0_\", \"png\", 72, width=NA, \"default\", \"F\")"
[86] "mSet<-PlotClustPCA(mSet, \"km_pca_0_\", \"png\", 72, width=NA, \"default\", \"km\", \"F\")"
[87] "mSet<-RF.Anal(mSet, 500,7,1)"
[88] "mSet<-PlotRF.Classify(mSet, \"rf_cls_0_\", \"png\", 72, width=NA)"
[89] "mSet<-PlotRF.VIP(mSet, \"rf_imp_0_\", \"png\", 72, width=NA)"
[90] "mSet<-PlotRF.Outlier(mSet, \"rf_outlier_0_\", \"png\", 72, width=NA)"
[91] "mSet<-SaveTransformedData(mSet)"
[92] "mSet<-PreparePDFReport(mSet, \"guest4008043714584141497\")\n"
```

---

The report was generated on Thu Apr 28 18:20:06 2022 with R version 4.0.2 (2020-06-22).
